# Pangenome analysis reveals the genetic basis for taxonomic classification of the Lactobacillaceae family

**DOI:** 10.1101/2023.05.16.541042

**Authors:** Akanksha Rajput, Siddharth M. Chauhan, Omkar S. Mohite, Jason C. Hyun, Omid Ardalani, Leonie J. Jahn, Morten OA Sommer, Bernhard O. Palsson

## Abstract

*Lactobacillaceae* represent a large family of important microbes that are foundational to the food industry. Many genome sequences of *Lactobacillaceae* strains are now available, enabling us to conduct a comprehensive pangenome analysis of this family. We collected 3,591 high-quality genomes from public sources and found that: 1) they contained enough genomes for 26 species to perform a pangenomic analysis, 2) the normalized Heap’s coefficient λ (a measure of pangenome openness) was found to have an average value of 0.27 (ranging from 0.07-0.37), 3) the pangenome openness was correlated with the abundance and genomic location of transposons and mobilomes, 4) the pangenome for each species was divided into core, accessory, and rare genomes, that highlight the species-specific properties (such as motility and restriction-modification systems), 5) the pangenome of *Lactiplantibacillus plantarum* (which contained the highest number of genomes found amongst the 26 species studied) contained nine distinct phylogroups, and 6) genome mining revealed a richness of detected biosynthetic gene clusters, with functions ranging from antimicrobial and probiotic to food preservation, but ∼93% were of unknown function. This study provides the first in-depth comparative pangenomics analysis of the *Lactobacillaceae* family.

## 1. Introduction

Lactic acid bacteria are important microorganisms in natural ecosystems as well as in the food industry (Duar et al., 2017). They are Gram-positive cocci and rods, unable to form spores in contrast to other Bacilli (de Almeida et al., 2021). Lactic acid bacteria share their ability to ferment carbohydrates anaerobically into lactate but can be divided into homofermenters, which produce only lactate as an end product, and heterofermenters, which additionally produce ethanol and carbon dioxide (Ayivi et al., 2020). The family of *Lactobacillaceae* contains 31 genera, of which 25 formerly belonged to the genus *Lactobacillus* (Zheng et al., 2020). It was hypothesized that lactic acid bacteria evolved from a common ancestor and adapted to a soil environment (Duar et al., 2017). Dramatic gene loss, in addition to horizontal gene transfer, has substantially shaped the differentiation and evolution of lactic acid bacteria into taxonomically different species, and high genetic variability and plasticity can be observed within individual species (Makarova et al., 2006).

The genetic variability of lactic acid bacteria can partly be explained by their different lifestyles and niche adaptations (O’Sullivan et al., 2009). These microbes are either free-living or associated with eukaryotic partners such as yeasts, plants, or animals (Duar et al., 2017). Interestingly, lactic acid bacteria form important relationships with humans and are among the most numerous bacteria positively interacting with humans (Makarova and Koonin, 2007). First, they are an essential part of the human microbiome and are present in multiple niches, including the vagina, oral cavity, and GI tract (Abou Chacra and Fenollar, 2021; Golshahi et al., 2021; Pessione, 2012). Second, lactic acid bacteria often dominate fermented food environments (Duar et al., 2017) and play crucial roles in food preservation (Zapaśnik et al., 2022) and flavor creation (Hu et al., 2022; Lorn et al., 2021). Indirectly, humans benefit from the association of lactic acid bacteria with plants, as they may play a role in plant health and growth promotion. Consequently, there is significant interest in understanding the ecology and evolution and the genetic and phenotypic capabilities of lactic acid bacteria to build a better basis for their practical applications. Some examples include probiotics (Raman et al., 2022), food starter cultures (Evivie et al., 2017), bioprotection agents (Bhat et al., 2012), and biocontrol (Strafella et al., 2020).

While these potential application areas are diverse, they share a common mechanism of action, which is the capability of lactic acid bacteria to modulate structures of multi-species microbial communities in ecological niches. In general, lactic acid bacteria create niche conditions that defend a specific environment or host from pathogens or spoilage organisms. Multiple mechanisms, linked to their metabolism and the production of bioactive compounds, form the basis for these effects. First, a number of metabolites like organic acids, including lactic acid, have been shown to have antimicrobial and antifungal activity (Siedler et al., 2019). Additionally, these metabolites have also been shown to directly interact with host organisms like humans and plants, promoting human health, for example, through anti-inflammatory pathways (Ohue-Kitano et al., 2018) or plant growth (Raman et al., 2022). Second, bioactive compounds such as peptide-based bacteriocins are produced by lactic acid bacteria and have been shown to have antimicrobial and antifungal properties (Sharma et al., 2022). Third, nutrient competition and rapid uptake of carbon sources (as well as micronutrients) have been shown to be important drivers for outcompeting fungal spoilage organisms (Siedler et al., 2020). While nutrient competition is difficult to assess based on genomic data, the metabolic capabilities as well as the potential production of bioactive compounds can be derived, to a large extent, from their genome sequences.

The genomics of lactic acid bacteria has been studied since 2001 when the first genomes were sequenced (Bolotin et al., 2001). Continuous efforts on individual genomes or pangenome analyses at the species or genus level (Carpi et al., 2022; Inglin et al., 2018) have been undertaken to characterize and understand the genetics of lactic acid bacteria as well as their evolution. In this study, using all publicly available genomes, we perform a family-wide pangenome analysis of lactic acid bacteria to map their functional genetic capabilities, including metabolic pathways and biosynthetic gene clusters. Thus, the large number of *Lactobacillaceae* genomes now available enables us to perform the first global comparative pangenome analysis at the family scale.

## 2. Methods

### 2.1. Data collection

For pangenomic analysis of the *Lactobacillaceae* family, we used the NCBI database to retrieve the genomic sequences (Sayers et al., 2022). The data retrieving and quality control steps include:

1. We retrieved 4,783 genomes of the *Lactobacillaceae* family from the NCBI database.
2. Next, we used The Genome Taxonomy Database Toolkit (GTDB-Tk) to re-annotate the taxonomy for all 4,783 genomes (Chaumeil et al., 2019).
3. Further, quality control and quality assurance (QC/QA) were done to get good-quality genomes. The QC/QA includes the taxonomy, number of contigs (<200), and N50(>50,000).
4. Finally, we get 3,591 high-quality genomes of the *Lactobacillaceae* family.

The 3,591 genomes were used for further pangenome and other downstream analyses.

### 2.2. Data processing

After extracting the high-quality genomes, we perform various analyses.

#### 2.2.1. Genome annotation

Initially, all the genomes were annotated using the Prokka software using stringent parameters (Seemann, 2014). In this study, we selected a group of 13 high-quality manually annotated genomes from the NCBI Genbank database (**Supplementary Table S5**). While running Prokka annotation we provided these 13 genomes as a priority to search annotations using the *--proteins* parameter. Further, the output of the Prokka annotation like GFF/ Genbank files is used for roary (pangenome), antiSMASH (biosynthetic gene clusters (BGCs)), and autoMLST (MLST tree).

#### 2.2.2. Pangenome construction

We used roary software to construct the pangenome (Page et al., 2015). Due to the high diversity among the *Lactobacillaceae family*, we selected 26 species with genomes >=30. The selected 26 species are *Lactiplantibacillus plantarum* (611)*, Pediococcus acidilactici* (193), *Lacticaseibacillus rhamnosus* (180), *Lacticaseibacillus paracasei* (172)*, Ligilactobacillus salivarius* (150)*, Limosilactobacillus reuteri* (100)*, Lactobacillus crispatus* (83)*, Pediococcus pentosaceus* (76)*, Oenococcus oeni* (72)*, Weissella confusai* (72)*, Lactobacillus delbrueckii* (67)*, Levilactobacillus brevis* (67)*, Leuconostoc mesenteroides* (62)*, Weissella cibaria* (57)*, Lactobacillus acidophilus* (54)*, Lactiplantibacillus pentosus* (52)*, Latilactobacillus sakei* (50)*, Limosilactobacillus fermentum* (49)*, Lactobacillus johnsonii* (47)*, Ligilactobacillus ruminis* (39)*, Lactobacillus paragasseri* (36)*, Leuconostoc inhae* (36)*, Lactobacillus helveticus* (32)*, Lactobacillus gasseri* (31)*, Lactobacillus iners* (30)*, Lentilactobacillus parabuchneri* (30). After running the roary we divided the pangenome into the core (>99%), accessory (<99% to >=15%), and rare (<15%) genomes. A list of the number of genes in all 03 categories in 26 species is provided in **Supplementary Table S2**.

#### 2.2.3. Biosynthetic gene clusters (BGCs) analysis

We detected BGCs across the selected 3,591 genomes of *Lactobacillaceae* family using antiSMASH v6.0.1 (Blin et al., 2021) (**Supplementary Table S6**). These detected BGCs can code for the biosynthesis of several secondary metabolites. To further identify the diversity of BGCs across different genomes, we used BiG-SCAPE v1.1.4 to compare the BGCs (Navarro-Muñoz et al., 2020). A combined distance metric was calculated between pairs of detected BGCs in this study and all the known BGCs from the MIBiG v2.0. The distance metric includes comparing protein domain strings across BGC genes in addition to domain sequence similarity. We used a cutoff of 0.3 on the combined distance to group BGCs into different gene cluster families (GCFs). If one of the known BGCs from the MIBiG database was found in the GCF, it was denoted as the known GCF. Using the distance matrix across all vs. all BGCs, a network was reconstructed using Cytoscape (Shannon et al., 2003) (Figure 5). The GCFs here represent different connected network components grouped based on the BGC type predicted using antiSMASH. The information on detected BGCs and GCFs across all genomes is enlisted in **Supplementary Table S7**. Further, we identified bacteriocins in the detected BGCs by calculating protein blast identity (>80%) against the database of 229 bacteriocins registered at the Bactibase (Hammami et al., 2010).

#### 2.2.4. Multi-locus species tree

The autoMLST software was used to reconstruct the multi-locus species for the members of the *Lactobacillaceae family* (Alanjary et al., 2019). The autoMLST software was used to reconstruct the phylogenetic tree for the individual 26 *Lactobacillaceae* species and the combined tree by taking the representative 307 species.

#### 2.2.5. Cluster of Orthologous Groups (COG) analysis

The pangenome reference fasta files from Roary for each species were processed using the eggNOG software to predict the further functional annotations and clustered ortholog groups (COG) (Huerta-Cepas et al., 2019). We used this software to annotate the genes of the *Lactobacillaceae* genomes among 27 different categories, for example, RNA processing and modification, Chromatin structure and dynamics, Energy production and conversion, Amino acid transport and metabolism, Nucleotide transport and metabolism, and many more as shown in **Supplementary Table S3**.

We performed Fischer’s exact test to identify the enrichment between the gene frequency group (CAR) and the COG functional category. For example, for category H in the core genome, Fischer’s exact test was applied to core vs. non-core genes and COG H vs. non-COG H genes. 26 species□×□3 frequency groups ×□20 COGs□=□1560 tests were conducted. Significance was determined based on FWER <□0.05 under Bonferroni correction or p-value <□7*10^− 5^. Log2 odds ratios (LOR) were computed between each frequency group and COG.

#### 2.2.6. Mobilome abundance and dispersion

In our compendium, we extracted genomic location, GC content, and gene length of all DNA modifying enzymes and mobilome. Then a case-by-case analysis was performed on *L. acidophilus*, *L. ruminis* and *L. parabuchneri* (species with the most closed pangenome) and *L. reuteri*, *L. helveticus,* and *L. crispatus* (species with the most open pangenome) to investigate whether there is a link between pangenome openness and elements with DNA modification properties. For these goals, Enzymes with DNA nuclease/ligase activity and mobilome were classified into ten functional groups, including 1) endonuclease, 2) CRISPR-Cas9, 3) transposase and prophages (mobilome), 4) recombinase, 5) repair system, 6) SOS system, 7) restriction-modification, 8) bacteriophage resistance, 9) toxin-antitoxin stability, and 10) integrase.

#### 2.2.7. Mash clustering

We utilized Mash, a program that quickly approximates nucleotide similarity between two genomes. Mash is an alternative to Pairwise calculation (correlation >95%) and is much faster than pairwise ANI, as described in the Mash manuscript by Ondov BD et al. (Ondov et al., 2016). Further, the Mash-generated phylogenetic trees are comparable to phylogenetic trees based on core genomes. We integrated it into an in-house Python script to create a table of pairwise distances for all 4,783 genomes collected (11,522,400 total). This table was filtered to include only genomes for which metadata from PATRIC was available (2,447) and then transformed into a square matrix of pairwise distance values. We converted the Mash distance from this square matrix into a similarity matrix using Pearson’s correlation coefficient as the metric to improve the clustering results and resulting visuals. These values were then subtracted from 1 to convert them back into distance measures in accordance with the protocol outlined by Abram et al. (Abram et al., 2021). This distance matrix was clustered in Python using the “clustermap’’ function in Seaborn with the “ward” method. This Mash clustering methodology was repeated for each of the top 6 pangenome species (by genome content).

#### 2.2.8. Heap’s law

Heap’s law was used to calculate the pangenomes’ openness and closeness, as Tettelin et al. proposed (Tettelin et al., 2008). We used Heap’s law to calculate the pangenome status of the genomes. Heap’s law is formulated as P=kN^λ^, where P is the pangenome size, N is the number of genomes, and k and λ are fitting parameters. The exponent λ determines the openness/closeness of the pangenomes. To compare differently-sized pangenomes, we randomly sampled 30 strains from each species for the Heaps’ Law fitting and repeated the process 50 times. This generated a range of λ values, from which we computed the mean and standard deviation. We call this value the “mean normalized λ” and use it to compare the different pangenomes. We have categorized the pangenomes into two categories as per *Hyun et al*.(Hyun et al., 2022), closed pangenome (λ<0.3) and intermediate open (λ>0.3).

Most of the data acquisition, curation, processing, and storage were carried out using a Snakemake (Mölder et al., 2021) based workflow that included rules for various steps of taxonomic identification, phylogenetic analysis, genome annotation, and pangenome reconstruction.

## 3. Results

### 3.1. Data selection and pangenome analysis workflow to identify qualified Lactobacillaceae genomes

We focused on characterizing the diverse *Lactobacillceae* family using various in-depth analytic methods, including pangenomics, phylogroup analysis, biosynthetic gene cluster (BGC) detection, and gene classification by clusters of orthologous genes (COGs) (**Figure 1A**). A total of 4,783 genomes of all assembly levels from the family *Lactobacillaceae* were downloaded from the NCBI RefSeq database on January 15, 2022. To achieve an accurate large-scale pangenome analysis, we used strict filters like the assembly quality of the starting dataset, accurate taxonomic definition, and consistent annotations of all genomes. Thus, the 4,783 genomes were processed based on the number of contigs (<200) and N50 value (>50 KB), which resulted in 3,591 medium (fragmented) to high-quality (complete) genomes after filtering out 1,192 poor-quality genomes (**Figure 1C**). The remarkable diversity of the *Lactobacillaceae* family is depicted by a wide range of genome sizes ranging from 1.2 Mbp (*Fructobacillus sp.* S1-1) to 3.96 Mbp (*Lactiplantibacillus pentosus* DZ35), with an average genome length of 2.4 Mbp (**Figure 1B**).

**Figure 1.**
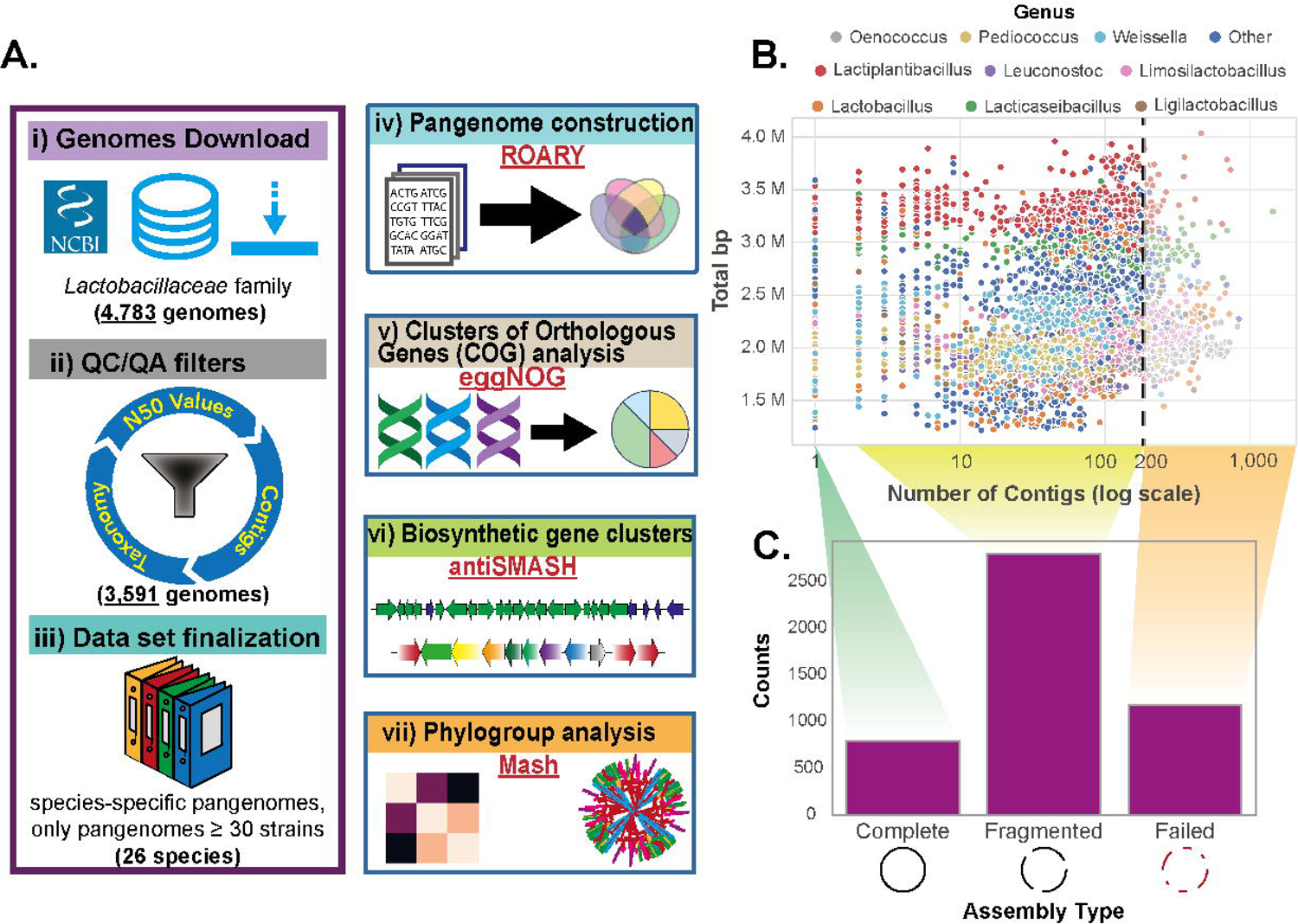
Workflow for data curation and subsequent pangenome analysis. A) Graphical summary of workflow and methodologies used. Genomes were downloaded from NCBI and filtered using N50 values, contigs, and GTDB taxonomy as quality control metrics. 26 species had ≥30 genomes available and were then selected for pangenome generation and analysis. The pangenome generation step outputs a gene presence/absence matrix, which we denote with **P**. This binary matrix lists gene as rows and strains as columns, with an entry of 1 if a gene is present in a given strain, and 0 if a gene is absent in a given strain. Information about the pangenome, phylogeny, secondary metabolites, clusters of orthologous genes, and Mash clustering analysis was used as a basis for characterizing the Lactobacillaceae family. B) Scatter plot showing the relationship between the number of contigs (log scale) and total base pair length of the 4,783 unfiltered Lactobacillaceae genomes, which were initially downloaded. The color of the dots represents various genera. Genomes that passed the quality control step had < 200 contigs, with their genome size varying from 1.2 Mbp to 4 Mbp. This genome size is generally correlated with genus classification. C) Bar plots depicting the assembly quality of Lactobacillaceae genomes downloaded from NCBI into 3 categories of complete genomes, high-quality fragmented genomes (< 200 contigs), and QC failed genomes with more than 200 contigs.

Given the substantial genomic diversity of the *Lactobacillaceae* family, we used a genome-based taxonomy database, GTDB (Chaumeil et al., 2022), to identify the genus and species for all 3,591 selected genomes (**Supplementary Table S1**). We detected a total of 344 species split across 33 genera. An additional 44 genomes were not assigned to any GTDB species, thus representing a few additional “orphan” species. The taxonomic reclassification using GTDB showed few differences from the NCBI genus definition, particularly for some species in three genera of *Lactobacillus*, *Lacticaseibacillus*, and *Lapidilactobacillus* (**Supplementary Figure S1**).

To assess the phylogenetic diversity across the whole family, we reconstructed an autoMLST-based (Alanjary et al., 2019) phylogenetic tree of the 307 strains that were classified as representatives at NCBI, spanning most of the captured genera and species (we excluded the 37 remaining species as their representative strains were not found at NCBI) (**Supplementary Figure S2**). The branching pattern within the resulting tree clearly shows significant phylogenetic diversity within and across the various genera in this family, with the *Lactobacillus* genus being the most diverse. Therefore, for the subsequent comparative pangenome analysis, we shortlisted the species with the highest number of genomes, resulting in 26 major species-specific pangenomes spanning 12 different genera, all of which had ≥30 medium to high-quality genomes available within the collected dataset.

We then constructed 26 species-specific pangenomes (a list can be found in **Supplementary Table S2**). *Lactiplantibacillus plantarum* had the highest number of strains (611), whereas *Lactobacillus iners* and *Lentilactobacillus parabuchneri* shared the fewest number of genomes (30 each). As previously mentioned, the 26 selected species span 12 different genera of the *Lactobacillaceae* family, with eight from *Lactobacillus* and four being single species that were sole representatives of their genus: *Oenococcus* (*oeni*), *Levilactobacillus* (*brevis*), *Lentilactobacillus* (*parabuchneri*), and *Latilactobacillus* (*sakei*).

We generated pangenomes for these 26 major species using Roary (Page et al., 2015) and divided the genes into the core, accessory, and rare (CAR) genomes, based on the gene frequency. The core genome consisted of genes present in at least 99% of all strains within the pangenome, while the rare genome consisted of genes present in less than 15% of all strains within the pangenome. The accessory genome consisted of genes between those ranges (15% - 99%). We then performed Mash clustering (Ondov et al., 2019, 2016) to showcase the genomic similarity across the *Lactobacillaceae* family (**Supplementary Figure S3**). Having 611 genomes available, we revealed nine potential phylogroups within *Lactiplantibacillus plantarum.* COG-based enrichment analysis was also performed to determine the functional annotation of the genes in the CAR genomes for all 26 species. Furthermore, antiSMASH (Blin et al., 2021) was run to discover all BGC predictions of secondary metabolite secretion within the *Lactobacillaceae* family. Thus, in this current study, we generated 26 *Lactobacillaceae* pangenomes from high-quality, publicly available strains and analyzed their pathways, BGCs, and phylogroups. The workflow described in this subsection is summarized in **Figure 1**.

### 3.2. Members of the *Lactobacillaceae* family have moderately open pangenomes

An open or closed pangenome status indicates a high or low probability, respectively, of finding a new gene family upon adding a new genome for pangenome analysis. Highly open pangenomes are typically associated with species inhabiting multiple environments and showing evidence of the exchange of genetic material (Medini et al., 2005; Reis and Cunha, 2021). The openness of a pangenome is characterized mathematically through Heaps’ Law, which describes the rate at which new genes are found when sampling an increasing number of strains within a pangenome (Hyun et al., 2022; Tettelin et al., 2008). This model is based on a power law characterized by its exponent parameter, which we denote as λ. This λ exponent value varies between 0 and 1, with 0 indicating a completely closed pangenome (with no new genes found as more strains are examined) and λ = 1 indicating a fully open pangenome (with a constant number of new genes found in each added strain). The numerical value of λ is therefore used to convey the openness of a given pangenome (Tettelin et al., 2008). A normalization step was also performed to compare these openness values between differently-sized pangenomes (see **Methods**) to generate normalized λ values for all pangenomes in this study.

The openness of the 26 pangenomes displayed a notable range in their diversity. 11 of the 26 species had a moderately open pangenome with normalized λ values > 0.3. 13 of the 26 species had a less open pangenome with normalized λ values between 0.2 and 0.3 (**Figure 2A**). Only two species had more closed pangenomes, *Lentilactobacillus parabuchneri,* with a normalized λ value of 0.171±0.004 and *Lactobacillus acidophilus,* with a normalized λ value of 0.077±0.021. *Limosilactobacillus reuteri* had the most open pangenome (Li et al., 2023), with a normalized λ value of 0.378±0.015, followed by *Lactobacillus crispatus,* with a normalized λ value of 0.364±0.017. The genera with the most open pangenomes were *Limosilactobacillus*, *Lactobacillus*, and *Levilactobacillus*. However, it is worth noting that many Lactobacillus species also had minimally open pangenomes, such as *L. acidophilus, L. gasseri*, and *L. iners*, which further showcases diversity in openness of pangenomes across the species in the *Lactobacillus* genus.

**Figure 2.**
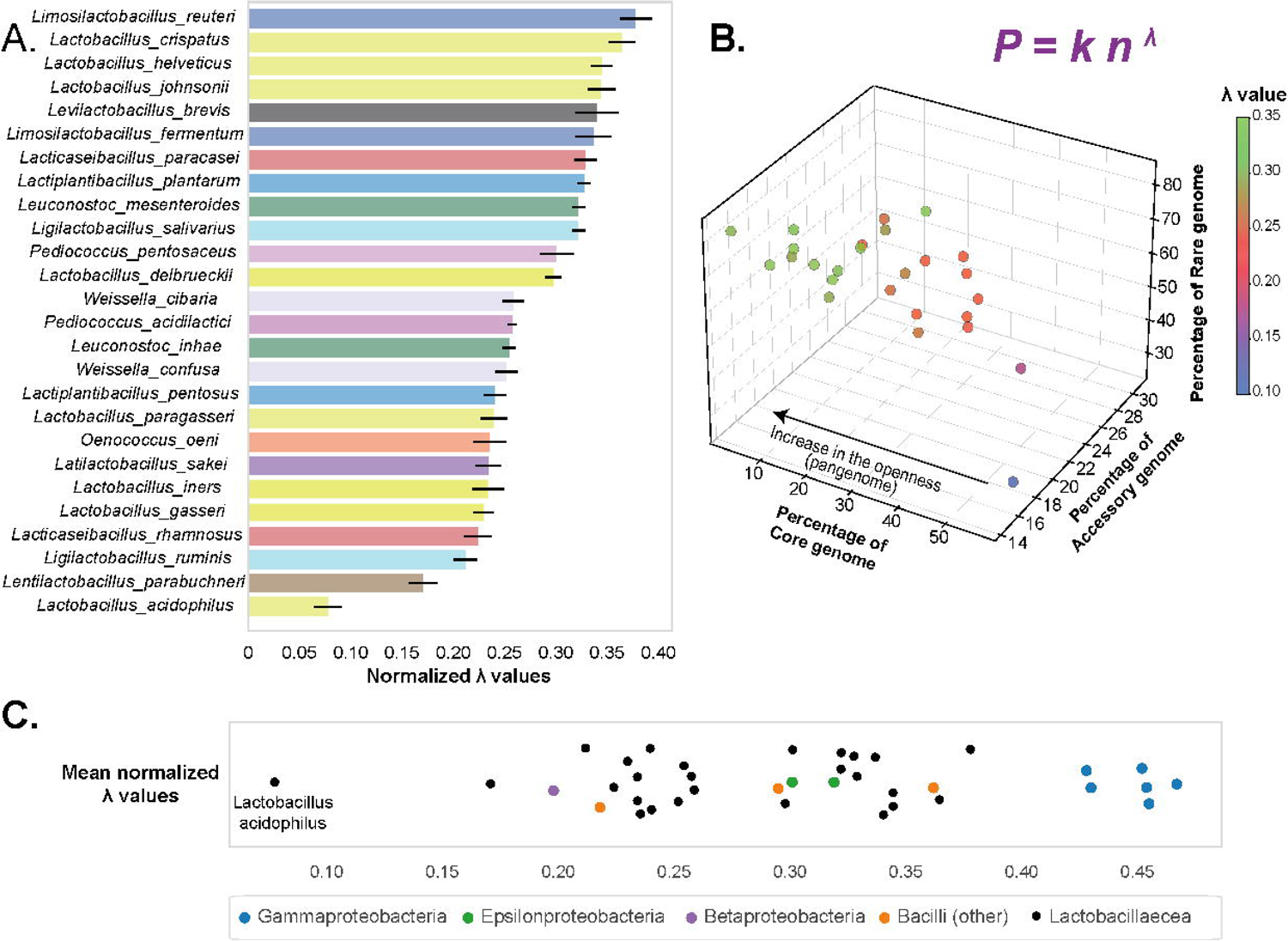
Pangenome status of Lactobacillaceae genomes. A) A bar plot showing rank-ordered, normalized λ values for each of the 26 pangenomes generated. The color of the bars depicts a specific genus of the Lactobacillaceae family. The λ values showcase moderately open pangenomes for most species, with genus-level correlation with openness. Lactobacillus acidophilus stands out as a clear outlier with its minimally open pangenome B) The 3D-scatter plot depicts the correlation between the percentage of genes in the core, accessory, and rare genomes with that of the λ values (color scaled) of 26 Lactobacillaceae species. We found a notable trend between the percentage of total genes in the core genome and the λ value of that species. C) A comparison of the mean normalized λ values of the Lactobacillaceae pangenomes calculated in this study vs. λ values calculated for other groups of bacteria from a previous study (Hyun et al., 2022). While none of the Lactobacillaceae have as open a pangenome as the gammaproteobacteria, they still occupy a much larger range than previously thought (the smallest reported bacilli pangenome previously had a λ value of 0.218), especially when looking at the pangenomes of Lactobacillus acidophilus and Lentilactobacillus parabuchneri.

We further evaluated the correlation between a species’ pangenome openness and its CAR genomes’ size. We found that the percentage of the genes in the core genomes was linked to the openness of the pangenomes (**Figure 2B**). *Lactobacillaceae* pangenomes with a low percentage of core genes possessed higher λ values, which signifies more openness of the pangenome. A small λ value indicates the same genes being found repeatedly in multiple sample strains, with few new genes added to a pangenome with each strain added. For example, species like *Lactobacillus crispatus, Limosilactobacillus reuteri, Lacticaseibacillus paracasei,* and *Lactiplantibacillus plantarum* have a low percentage of total genes in their core genomes (i.e., 9.24%, 10.04%, 6.48%, and 2.77%, respectively) and high λ values (i.e., 0.345 ± 0.032, 0.344 ± 0.021, 0.316 ± 0.016, 0.301 ± 0.011) (**Figure 2B**). Likewise, species with low λ values, such as *L. acidophilus* and *L. parabuchneri* had a high percentage of total genes in their core genomes.

We also compared the *Lactobacillaceae* pangenome openness to other bacteria such as Gammaproteobacteria, Epsilonproteobacteria, Betaproteobacteria, and (non-*Lactobacillaceae*) Bacilli from a previous comparative pangenomics study (Hyun et al., 2022). Through this comparison, we found that *Lactobaciilaceae* have moderately open pangenomes and λ values generally in line with other Bacilli strains. However, *Lactobacillus acidophilus* is shown to be an outlier, with a significantly smaller λ value (and thus a correspondingly more conservative pangenome) than any other species (**Figure 2C**).

In summary, the pangenomes of the 26 major *Lactobacillaceae* species have moderately open pangenomes but contain notable diversity, especially within the *Lactobacillus* genus.

### 3.3. Elucidating the characteristics of core, accessory, and rare genomes of *Lactobacillaceae*

The CAR genomes play an important role in defining the pangenome characteristics of a microbial species. Therefore, we explored the pangenome characteristics of all 26 *Lactobacillaceae* species individually. CAR gene numbers vary across the 26 species (**Supplementary Figures S4-S29**).

We describe the pangenome characteristics of *L. plantarum* in detail, as it had the highest number of genomes (611) in our compendium. *L. plantarum* is a critical member of the genus *Lactiplantibacillus* and is commonly found in fermented food products and anaerobic plant matter (Surve et al., 2022). We calculated the individual gene frequencies and generated a cumulative gene distribution plot (**Figure 3A**) (Hyun et al., 2022). This plot for the *L. plantarum* pangenome displays a sharp change in slope near the edges where both the core and rare genome cutoffs are located, whereas the region defined as the accessory genome is relatively flat by comparison. These characteristics are generally found in the other 25 *Lactobacillaceae* pangenomes studied (**Supplementary Figures S4-S29**).

**Figure 3.**
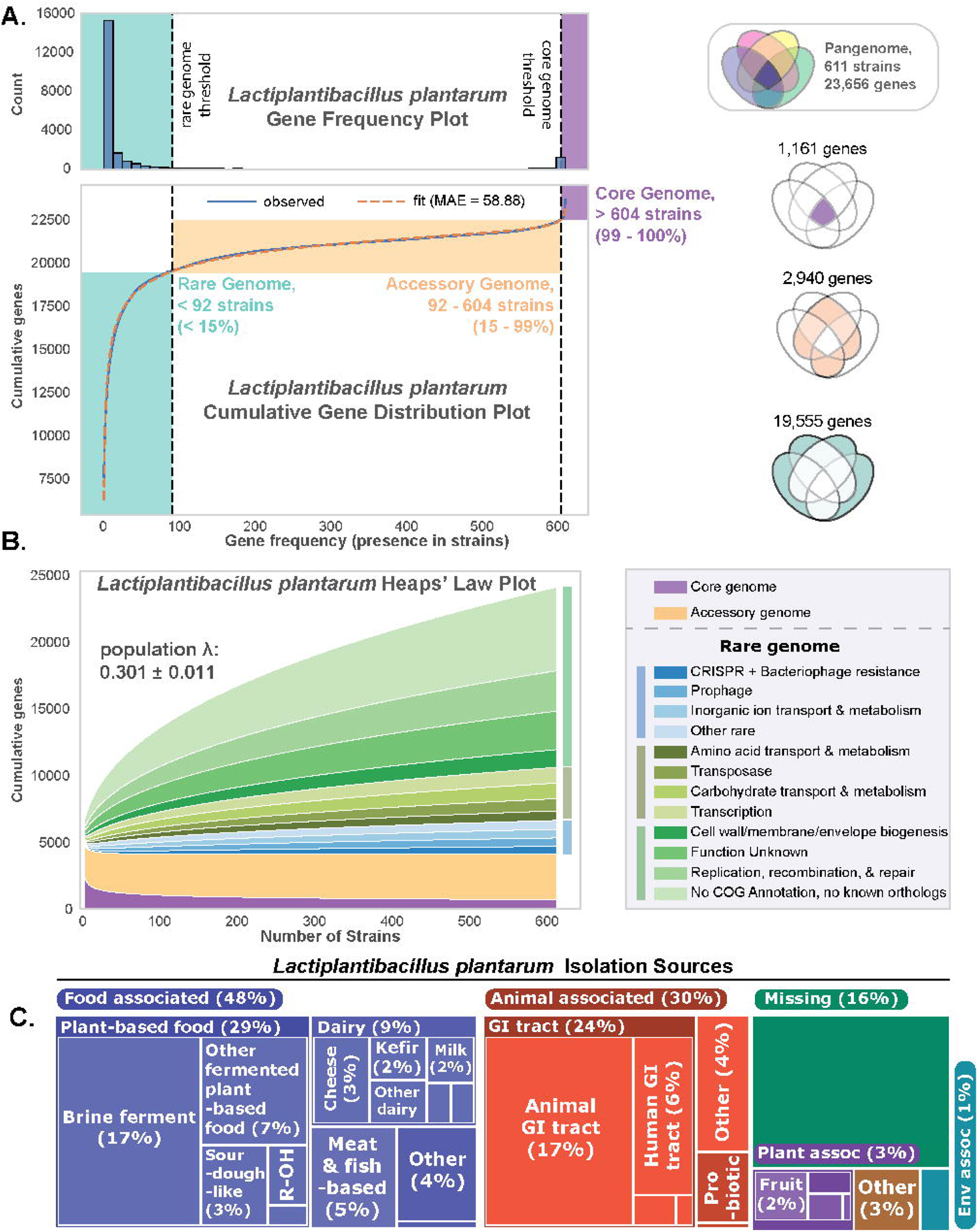
The pangenomic analysis of Lactiplantibacillus plantarum. A) Cumulative gene frequency distribution function for 611 strains of the L. plantarum pangenome. The gene frequency distribution function is fitted to a double-exponential model (with median absolute error or MAE = 58.88), from which the corresponding cumulative gene distribution function is generated (Hyun et al., 2022). The genes in the pangenome are divided into the core (comprising 1,161 genes), accessory (comprising 2,940 genes), and rare (comprising 19,555 genes) genomes. B) Heaps plot of L. plantarum showing the number of genomes vs. genes present. 14 curves are shown, representing the core genome, accessory genome, and subsections of the rare genome. While the core and accessory genome sizes quickly stabilize, the rare genome (and all its highlighted subsections) keeps increasing via a positive power-law relationship. This plot also shows the calculated population λ value for this species’ pangenome at 0.301±0.011 (note: this is not the same as the normalized λ value used in Figure 2 for comparison between pangenomes, which was obtained via sampling (see Methods). Instead this uses the whole pangenome population for this species to generate the population λ value). C) The treemap shows the environmental niche from which the 611 L. plantarum strains were isolated.

These sharp changes in slope near the elbows of the cumulative gene distribution function represent defining characteristics of the pangenome. The frequency of the core genes is much higher than those in the accessory genome. This increased frequency, as evidenced by the near-vertical slope by the elbow of the plot, is indicative of core genes being present in (nearly) all strains. The near omnipresence of these 1,161 core genes in the *L. plantarum* pangenome, despite these strains inhabiting a diverse set of niches, reveals that these genes are unlikely to be lost due to mutations, adaptations to the environment, or other evolutionary events. This observation also applies to the remaining *Lactobacillaceae* pangenomes, which incidentally have a similar number of core genes to *L. plantarum*, averaging 1,197±273 genes. The Heaps’ plot (**Figure 3B**) showcases that, while *L. plantarum* has a moderately open pangenome (population λ value of 0.301±0.011), the core genome itself is quite conserved and does not change much beyond a sample size of 300 strains.

The *L. plantarum* accessory genome has 2,940 genes with varying presence across the 611 strains. This sizable range suggests that these genes (many of which are involved in nucleotide metabolism, coenzyme metabolism, translation machinery, and energy production) provide varying fitness levels, likely environment-dependent. We thus examined the niche distribution of the strains in *L. plantarum*. We found that the plurality of the genomes in the *L. plantarum* pangenome was isolated from plant-based food (Yu et al., 2021), followed by the gastrointestinal tract (Echegaray et al., 2023) and milk-based food (Ngamsomchat et al., 2022) (**Figure 3C**). The genes in the accessory genome led to the genetic basis for defining phylogroups of *L. plantarum*, as detailed in a later subsection. It is worth noting that *L. plantarum* has the largest accessory gene pool amongst the 26 pangenomes in our compendium, with the average accessory genome profile consisting of 1,523±595 genes. It is also worth noting that the accessory genome does not change much beyond a sample size of 200 strains, as shown in Heap’s plot (**Figure 3B**).

In contrast, most of the 19,555 genes in the rare genome were present only in a handful of the 611 strains, with a rare gene present in only three strains on average (**Figure 3A**, blue-green area). Many of these rare genes are either transporters (3.5%), glycosyl hydrolases and glycosyltransferases (4.3%), transposases (4.6%), phage-related genes (5.4%), transcriptional regulation genes (5.7%), or genes with no functional annotations (48.7%). This observation suggests that they represent each strain’s unique experiences and history. Namely, these genes are likely the results of rare horizontal gene transfer events, mutations within duplicated and/or transposed genes or represent potential “battle scars” from phage fights in the strain’s past.

In summary, we found that the 26 species-specific pangenomes within the *Lactobacillaceae* family have a set of 1,197±273 core genes, with *L. plantarum*, in particular, consisting of 1,161 core genes (comprising 38% of the average genome). On average, most of the niche adaptations for these species come from their accessory gene profile, which contains 1,523±595 genes. These accessory genes generally code for enzymes involved in nucleotide metabolism, coenzyme metabolism, translation machinery, and energy production and suggest adaptations for the various environments in which these strains thrive. The rare genome contains many genes found in <15% of genomes and averages about three genes per strain. Thus, these genes are truly rare and represent unique or rare events in the history of that particular strain. Cumulative gene distribution plots of the remaining 25 *Lactobacillaceae* genera are provided in **Supplementary Figures S4-S29**.

### 3.4. All parts of the pangenome of the 26 species are composed of functionally diverse genes

As suggested by our analysis of the *L. plantarum* pangenome, COG annotations of genes can aid in identifying the root cause of pangenome openness and understanding species adaptations to their environment. We utilized eggNOG (Cantalapiedra et al., 2021; Huerta-Cepas et al., 2019) to determine the COG annotations for all genes in the CAR genomes. We then performed a Fisher’s Exact Test (Bonferonni correction, family-wise error rate of 0.05) to confirm the statistical enrichment of the assigned functions within the CAR genomes across all 26 species. The list of all 27 COG categories is provided in **Supplementary Table S3**.

Of the 26 COG categories, 14 are statistically enriched within the core genome, of which three categories are enriched in more than 20 of the 26 species (**Figure 4A**). These three categories are translational, ribosomal structure and biogenesis (J, 26/26); nucleotide transport and metabolism (F, 25/26); and post-translational modification, protein turnover, and chaperones (O, 22/26), and are among the categories most consistently enriched within core genomes, as previously observed across multiple microbial pathogens (Hyun et al., 2022). Among the J category, enriched genes included ribosomal 50S (RpmA-G, RpIB-X) and 30S (RpsC-S) proteins; tRNA modification (SpoU, TrmL); and 23S (MrnC, RluA) and 16S rRNA (RsuA) modification. In the case of the F category, both the pyrimidine biosynthesis (Adk, PyrG) and salvage (Upp) pathways, as well as the purine biosynthesis (PrsA) and salvage (Apt) pathways were found to be enriched in 25/26 species except *L. acidophilus*. For the O category, we discovered co-chaperonin GroES and ATP-dependent proteases (HflB, ClpA, ClpP, HsIV) to be enriched in 22/26 species (**Supplementary Table S4**). Interestingly, the carbohydrate transport and metabolism function (G) is only found in *Leuconostoc inhae*, which is a lactic acid bacterium isolated from kimchi, a fermented vegetable food produced in Korea (log2odds ratio of 0.93) (Kim et al., 2003). Furthermore, the cell motility category (N) was enriched in the core genome of the two important *Lactobacillaceae* species typically isolated from the human intestine: *Lactobacillus helveticus* and *Ligilactobacillus ruminis*. More specifically, the genes in these categories encoded for a type IV pilus system (T4PS) (PulE-O) and a flagellum hydrolase (FlgJ), which have a role in microbiota-host interaction (Ligthart et al., 2020). Thus, the core genomes of the 26 species were primarily enriched with genes involved in information storage and processing functions (i.e., J) followed by essential metabolic functions (i.e., F).

**Figure 4.**
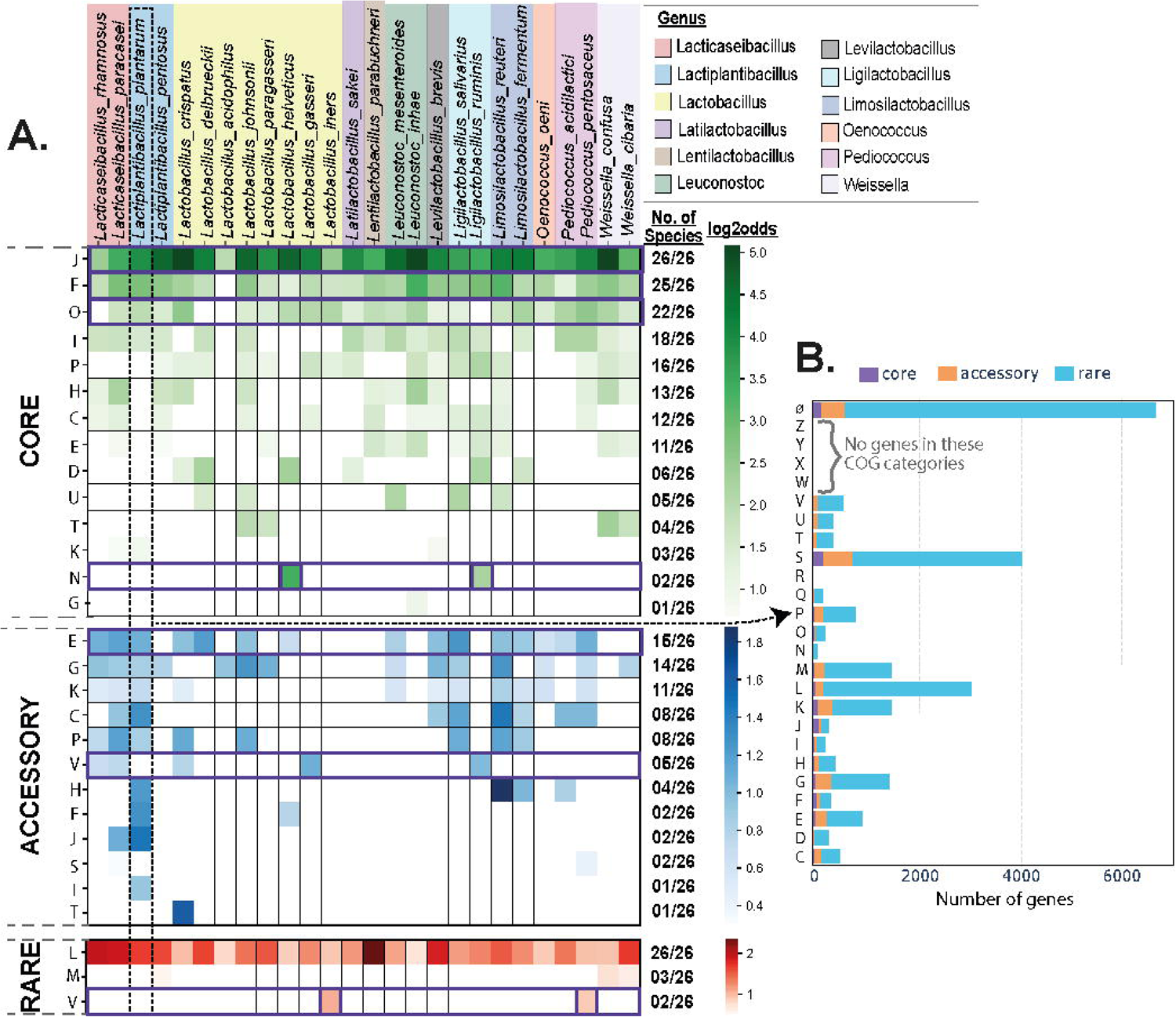
The Clusters of Orthologous Genes (COG) category analysis of the Lactobacillaceae species. A) Heat map showing the log2odd ratio among the core, accessory, and rare pangenome for the 26 species. The top horizontal scale shows the species name highlighted with a specific genus. The color intensity of the squares is as per the log2odds ratio. The right vertical axis scale shows the frequency of the occurrence among species with respect to all 26 species. Only enrichments with a p-value < 0.05 are displayed. ‘S’ and ‘-’ categories are not shown in the heatmap. B) The barplot depicts the COG functions among core, accessory, and rare pangenomes in L. plantarum. The barplot is between the number of genes vs. COG functions. Note: “ Ø ” refers to genes with no COG annotation and no known orthologs.

For the accessory genome, we discovered 12 enriched COG functional categories. The amino acid transport and metabolism function (E) was the most enriched among the accessory genomes (in 15/26 species). Specifically, the genes involved in glutamine biosynthesis (i.e., GlnA); serine transporters (i.e., PotE); and ABC-type polar amino acid transport system (i.e., GlnQ) were enriched in E. Interestingly, in the accessory genomes, the carbohydrate transport and metabolism (G) is reported in 14/26 species. Further, the defense mechanism category (V) is enriched in the accessory genomes of five *Lactobacillaceae* species. The common genes involved in these five species are the ABC-type transport system (CcmA, YadH, MdlB); ABC-type antimicrobial peptide transport system (SalY); and the type I restriction-modification system (HsdM). These ‘V’ category functions involve antibiotic resistance and innate immunity in bacteria (Feng et al., 2020; Huo et al., 2019). Additionally, we discovered that lipid transport and metabolism (I) and signal transduction mechanisms (T) were enriched solely within *Lactiplantibacillus plantarum* and *Lacticaseibacillus crispatus* (log2odds ratio of 0.90 and 1.61, respectively). Our analysis found that accessory genomes were mostly enriched with various metabolism-related functions (i.e., E and G) and transporter genes.

In the case of the rare genome, we identified only three enriched COG categories. The replication, recombination, and repair (L) function were maximally enriched in all 26 species. Two other COG functions, cell wall/membrane/envelope biogenesis (M) and ‘V’, were also identified in rare genomes. Interestingly the common genes in the rare genomes are beta-lactamase class A (PenP); type I and type II restriction-modification system (HsdM, YeeA); abortive infection bacteriophage resistance (AbiF); ABC-type antimicrobial peptide transport system (SalY); and ABC-type multidrug transport system (YadH). Therefore, rare genomes were mostly enriched with information storage and processing functions (i.e., L). Additionally, the rare genomes are mostly enriched in unknown COG function (S) or no COG annotation (Ø) compared to core and accessory genomes, suggesting that the genes in these rare genomes are poorly characterized and warrant further investigation.

We also evaluated the COG category for the 26 species individually (**Figure 4B** and **Supplementary Figures S30-S54**). For example, in *L. plantarum*, we found that the maximum percentage of the functions is not predicted by eggNOG. However, the most preferred and known functions are replication, recombination, & repair ‘L’, transcription ‘K’, and cell wall/membrane/envelope biogenesis ‘M’ (**Figure 4B**).

Taken together, the results show that the ‘J’ function is enriched in all the core genomes of the 26 species, which include rRNA and tRNA modification and biogenesis. For the accessory genome, we did not find any function conserved among all 26 species, whereas, for the rare genomes, we identified the ‘L’ function conserved among all 26 species, which might be responsible for the genetic diversity among species. Importantly, we find the ‘V’ function in rare genomes, specifically the restriction modification and abortive bacteriophage resistance systems.

### 3.5. Abundance and genome-wide dispersion of elements involved in genomic structural changes correlate with pangenome openness

To understand the underlying reasons for the varied openness of the *Lactobacillaceae* pangenome, we analyzed the abundance and distribution of all the genetic elements that are associated with structural changes in the genome. Here, we selected genes from ten functional categories for detailed assessment: 1) endonuclease, 2) CRISPR-Cas9, 3) transposase and prophages (mobilome), 4) recombinase, 5) repair system, 6) SOS system, 7) restriction-modification, 8) bacteriophage resistance, 9) toxin-antitoxin stability, and 10) integrase. All these functions have been reported to have DNA cleavage and/or recombination activity, resulting in genome instability in terms of losing or acquiring new genes (Darmon and Leach, 2014) (**Figure 5A**). Many of these functions belong to COG categories V (enriched in both rare and accessory genomes), L (only enriched in the rare genome), and S (only enriched in the rare genome).

**Figure 5.**
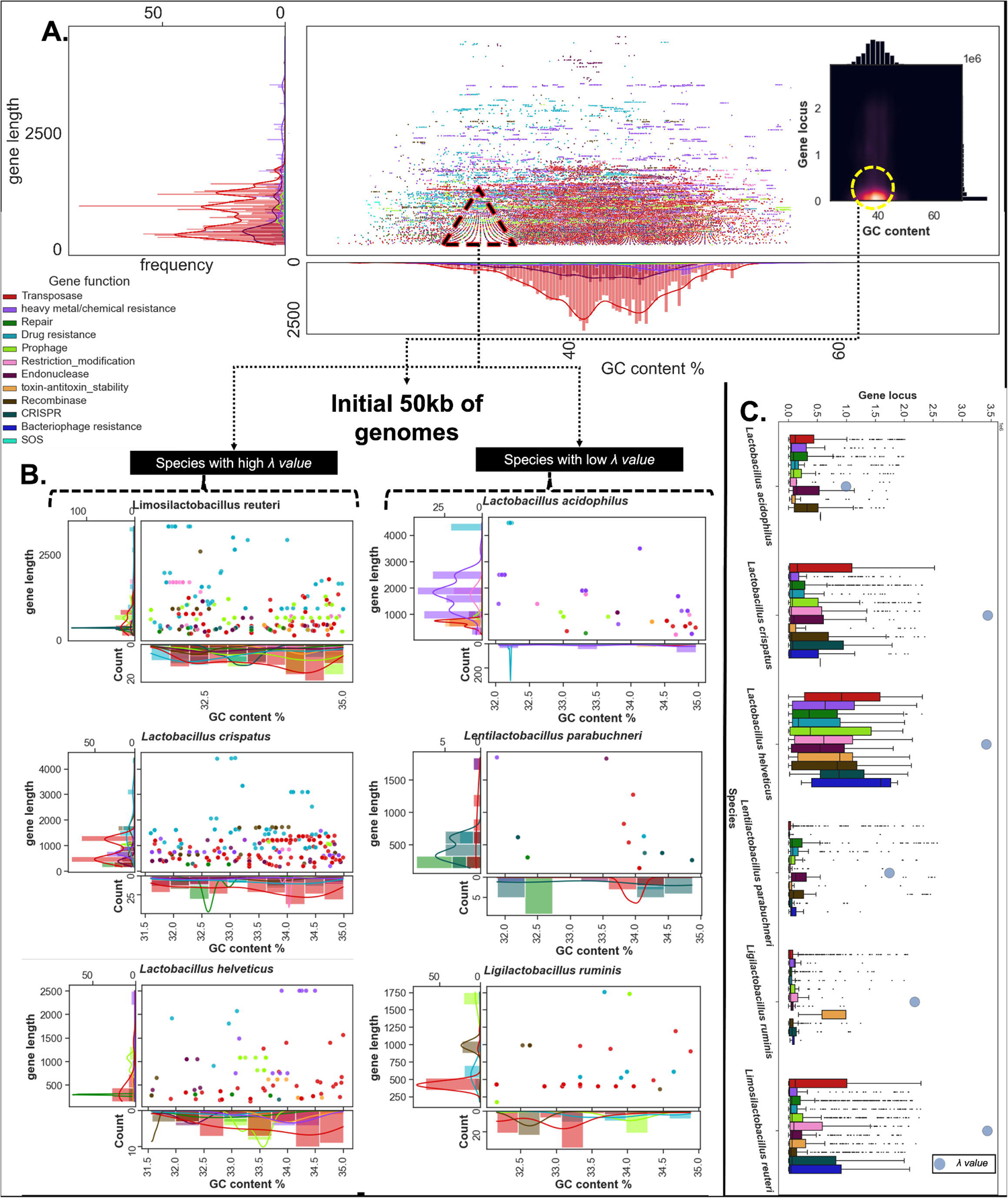
Role of mobilome in genome instability, and pangenome openness. A) A scatter plot shows the GC content and gene length of enzymes involved in structural changes in the genomes of Lactobacillaceae. The color of the dots represents the functions of enzymes. B) A zoom-in on a region with low GC content and located in the first 50k base pairs downstream from the origin of the replication on Lactobacillaceae genomes for species with the most open and the most closed pangenome. C) Boxplots show the distribution of gene locus across the genomes of selected species. Gene functions are represented by the color bar, and λ values are represented by the gray dots.

Genes belonging to these structural categories were selected from all the genomes across 26 species of *Lactobacillaceae*. Gene length and GC content of transposable elements (TEs) have been shown to play an important role in mobility (Boissinot, 2022; Feschotte and Pritham, 2006). We assessed the variation in the GC content and gene length across the genes belonging to different categories. By plotting gene length and GC content of the mobilome, several symmetrical patterns were observed by dividing TEs by different GC content (**Figure 5A**). This observation shows that *Lactobacillaceae* TEs have seven preferred niches based on GC content across *Lactobacillaceae* genomes. For example, we found the GC niches at 42% and 46% optimal for TEs, possibly allowing for HGT or gene disruptions in these areas. Furthermore, we also noted that many of the hypothetical genes from the rare genomes are collocated with TEs from these two optimal GC niches.

We further analyzed the genes with low GC content (31.5-35%) enriched in the first 50kbp downstream from the origin of replication of all *Lactobacillaceae* genomes. We discovered the species with the most closed pangenomes, like *Lactobacillus acidophilus*, *Lentilactobacillus parabuchneri, and Lactobacillus ruminis,* show a lower number of TEs and other elements in this region. In comparison, the number of TEs and other elements increases with the openness in the pangenome in species like *Lactobacillus crispatus, Limosilactobacillus reuteri,* and *Lactobacillus helveticus* (**Figure 5B**). For example, a group of genes in this important region primarily coded for functions like TEs, prophages, SOS, and heavy metal/ chemical resistance (**Figure 5B**). We also found that SOS functional genes in the first 50 kbp region of the genome were specifically enriched at the GC content of 32.2% in the *L. acidophilus* species.

Next, we investigated if the scattering pattern of these elements across genomic locations was correlated with pangenome openness. Interestingly, the species with a dispersed distribution of TEs throughout their genomes have the most open pangenomes (**Figure 5C**). In contrast, species with closed pangenomes have TEs accumulated in the 50kbs downstream of the origin of replication. We find the same observation using the CRISPR system. For *L. helvticus* species, most of the categories were dispersed across the genome, whereas for the two other species i.e. *L. reuteri* and *L. helveticus*, only a few of the categories were dispersed (e.g., TEs and CRISPRs). In the *L. ruminis* species, all but the toxin-antitoxin-related genes were accumulated in the first 50kbp region. Overall, these insights show the high importance of the abundance and dispersion of different genetic elements in the species’ pangenome status (openness and closeness). Further investigations into this hypothesis may reveal the underlying forces behind species diversification and adaptation.

### 3.6. Mash-based analyses of the *Lactiplantibacillus plantarum* pangenome reveal nine distinct phylogroups

We utilized Mash (Ondov et al., 2016), a program that approximates the similarity between two genomes based on their nucleotide content, alongside an in-house Python script, to create a distance matrix for all 3,591 *Lactobacillacea* genomes in our compendium. This matrix was then clustered using hierarchical clustering to produce a heatmap illustrating the population structure of all analyzed strains (**Supplementary Figure S3**). From there, we focused on the genomes of *L. plantarum*, the largest species-specific pangenome in our compendium (consisting of 611 strains). Clustering this submatrix (consisting of only *L. plantarum* genomes) using hierarchical clustering produced a heatmap that differentiated nine distinct clusters within the species (**Figure 6A**). We hypothesize that these clusters represent distinct phylogroups within this species. Interestingly, one of these clusters, Cluster A, is quite distinct from all other Mash clusters (**Figure 6B**). While most of the Mash distances between *L. plantarum* strains are between 0.005 and 0.017, the Cluster A strains have Mash distance values between 0.025 and 0.040 against strains from other clusters. This result may suggest that this subgroup of strains comes from either a distinct evolutionary history or unique environments and may form separate subspecies within *L. plantarum*. This result has been reported previously in literature focusing on *L. plantarum*, where a divergent set of strains seems to emerge, forming its phylogenetic clade (Li et al., 2022).

**Figure 6.**
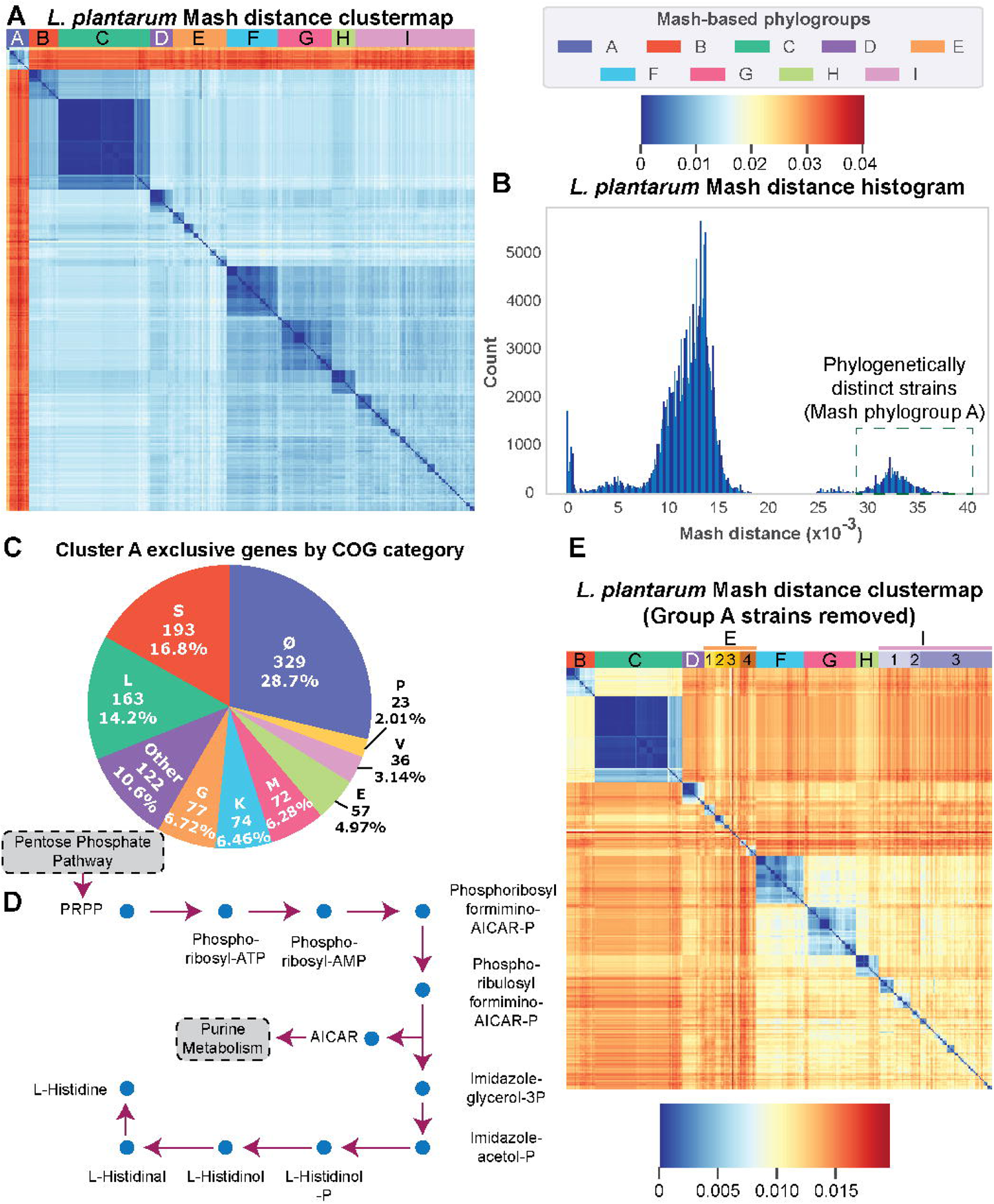
Mash-based analysis of L. plantarum reveals nine phylogroup candidates. *A) A clustermap of Mash distances for all* L. plantarum *strains. Nine different Mash clusters are formed, which may represent potential phylogroups for this species. B) A histogram of raw Mash distances between all* L. plantarum *strains. The Mash distances between Mash-phylogroup A and the rest of the strains (highlighted in the green box) in the* L. plantarum *pangenome suggest this group of strains is quite distinct from the rest of the strains, either a phylogenetically distant phylogroup or as an entirely separate subspecies of* L. plantarum*. C) A pie chart showcasing the distribution of exclusive genes found within Cluster A by COG category. The top 9 COG categories are highlighted in this chart, with the remainder being grouped into the “Other” section (note: “ Ø ” refers to genes with no COG annotation and no known orthologs). D) A metabolic map showcasing the PRPP to L-histidine pathway enriched within the Cluster A exclusive genes. E) A clustermap of Mash distances for all* L. plantarum *strains, with the Cluster A strains removed. All the former clusters in Panel A are the same, except for Cluster E, which splits into 4 clades, and Cluster I, which splits into 3 clades*.

Looking further into the gene content of Cluster A strains, we find 1,146 exclusive genes mapped to known orthologs using eggNOG (Huerta-Cepas et al., 2019). The COG category breakdown reveals that 329 (28.7%) of these genes are not functionally annotated in addition to not having any known orthologs, with an additional 193 (16.8%) mapping to established orthologous genes of unknown function. The largest known grouping for these exclusive genes belongs to ‘L’ (163 genes, 14.2%), followed by ‘G’ (77 genes, 6.72%), ‘K’ (74 genes, 6.46%), ‘M’ (72 genes, 6.28%), ‘E’ (57 genes, 4.97%), ‘V’ (36 genes, 3.14%), and ‘P’ (23 genes, 2.01%) (**Figure 6C**). A Fisher’s Exact Test was performed (Benjamini–Hochberg correction, false-discovery rate at 0.01), which found that these Cluster A exclusive genes were enriched for genes in the KEGG-defined histidine metabolic pathway, specifically for the histidine biosynthesis KEGG module responsible for converting PRPP into histidine (**Figure 6D**). Upon closer examination, we found that genes encoding this module are also found in other phylogroup candidate strains. Still, those genes are not the same as the ones exclusive to Cluster A, most likely coding for isozymes. It should be noted that most of these exclusive genes have minimal-to-no functional annotation and warrant further investigation to fully understand this phylogenetically distinct subgroup within *L. plantarum*.

As reported earlier, our analysis *via* Mash clustering finds nine phylogroups. Of those, Mash Cluster A is the most distinct and may represent a more distantly related set of strains. As such, we reran our Mash analysis after excluding these strains and found that the same Mash clusters are still present, except Mash Cluster E, which splits into four clades, and Mash Cluster I, which splits into three clades (**Figure 6E**). This suggests robustness within the remaining Mash phylogroups.

In summary, our Mash-based analysis finds nine distinct phylogroups within *L. plantarum*. One of these phylogroups, Cluster A, is quite distinct from the rest and may form its subspecies. There are 1,146 genes exclusively found in Cluster A strains which contribute to these strains being more genetically distant from the rest of the *L. plantarum* strains. 522 (45%) of these genes have no known putative function assigned to them. Of those with a known COG annotation, many are either recombinases, transposases, membrane-related, or sugar metabolism-related genes. A Fisher’s Exact Test finds that these exclusive genes are also enriched for genes spanning the histidine biosynthesis pathway, specifically in relation to converting from PRPP into histidine. A closer look at these genes reveals that they are in fact, isozymes that are separate from the other histidine biosynthesis genes found in the remaining *L. plantarum* strains.

### 3.7. Genome mining of the rare genome reveals mostly uncharacterized secondary metabolite BGCs across *Lactobacillaceae*

Secondary metabolites play an important role in microbial life. Their production is encoded by biosynthetic gene clusters (BGCs). Genome mining tools such as antiSMASH can predict various types of BGCs in the genome and thus help identify novel bioactive compounds (Blin et al., 2021). The most studied BGCs within lactic acid bacteria primarily encode for bacteriocins that have antimicrobial properties towards specific species and strains (Donia et al., 2014; Soltani et al., 2021; Zacharof and Lovitt, 2012). Nisin is one of the best-known examples of bacteriocins with wide applications in the food industry (de Arauz et al., 2009).

We predicted 6,950 BGCs across 3,117 genomes, with the remaining 474 genomes having no BGCs detected (Figure 7). The average number of BGCs across the whole family of *Lactobacillaceae* was 1.94, with 51 of the 345 species initially detected possessing no BGCs. The average number of BGCs varied among all collected species across the family.

**Figure 7.**
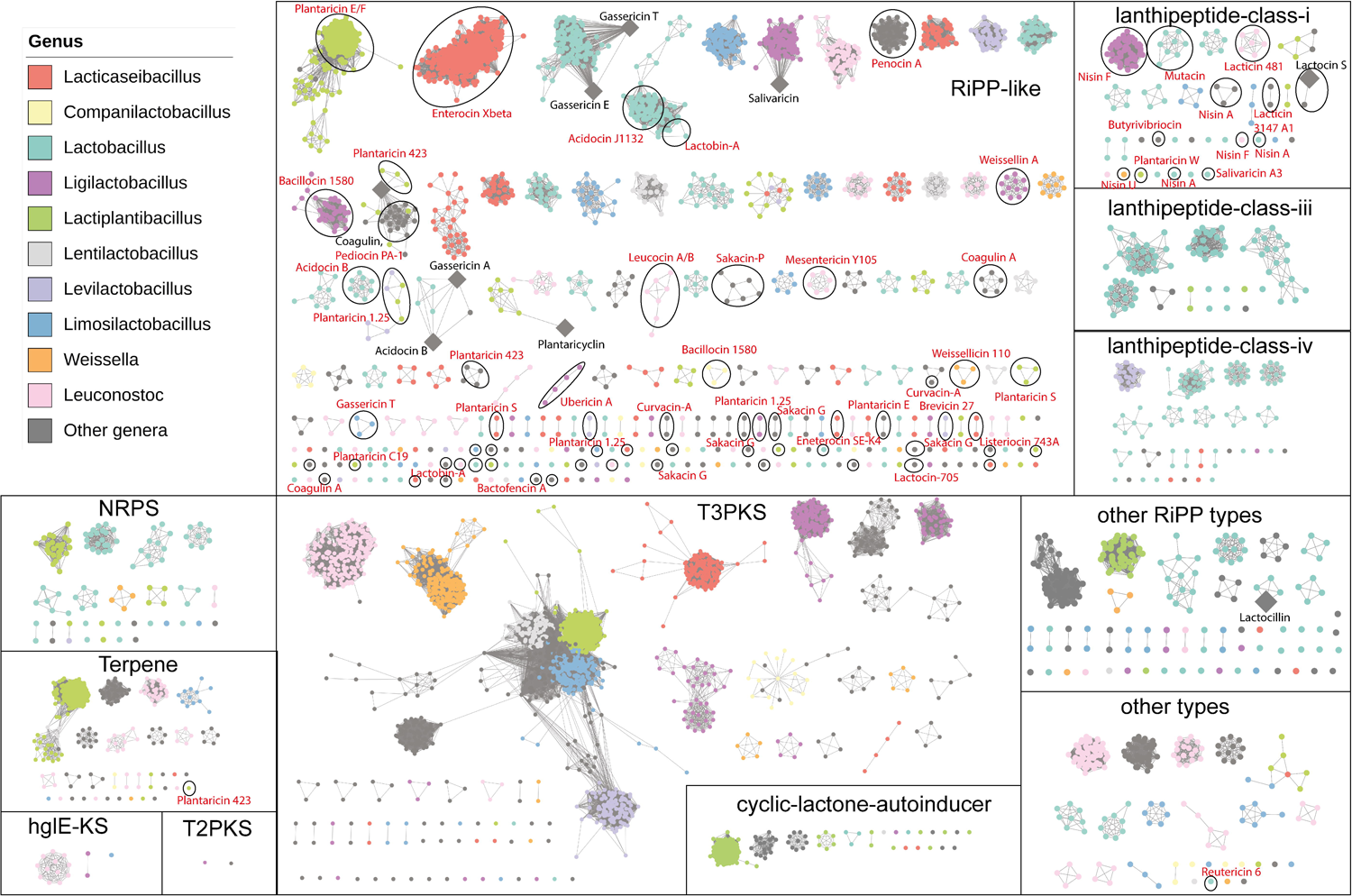
Overview of the biosynthetic gene clusters in the Lactobacillaceae family. The BGC map shows the distribution of different BGC classes among the members of the Lactobacillaceae family. The color represents the genus, the nodes represent BGCs, and the edges show similarities between BGCs. Many of the BGCs are annotated using top Blastp hit against the Bactibase database of bacteriocins (mimimun similarity of 50% is considered).

Among the 26 species where detailed pangenome analysis was possible, *Pediococcus acidilactici* possessed the lowest number of average BGCs (∼0.1 BGCs on average across 193 genomes, with 175 of them having no detected BGCs), whereas *Lactiplantibacillus plantarum* reported the highest number of average BGCs (∼4 BGCs on average across 611 genomes). The most abundant types of BGCs included RiPP-like (2,322), T3PKS (2,319), terpene (788), cyclic-lactone-autoinducer (643), lanthipeptide-class-iv (130), RRE-containing (90), lanthipeptide-class-i (82), lanthipeptide-class-iii (78), and beta lactone (72). Some BGCs were hybrid types, involving more than one type of BGC in the region. The top category included a hybrid of the types cyclic-lactone-autoinducer and T3PKS (33), which could lead to two separate metabolites or a hybrid biosynthesis.

We compared all BGC families using BiG-SCAPE (Navarro-Muñoz et al., 2020) to generate a similarity network to identify distinct BGCs (**Figure 7**). We detected 532 distinct BGC families predicted to encode enzymes for different secondary metabolites. Most of these families were present in one or a few genomes, thus representing BGCs as a feature of the rare genome. When compared against the standard database of known BGCs, we could identify only seven of these families coding for known putative BGCs in the MIBIG database, whereas we further detected around 30 families with known bacteriocins from Bactibase (Hammami et al., 2010) and other BGCs from the literature search. We also found that 281 families contained only one BGC, thus representing the unique potential of these genomes.

The most abundant family with 1,331 BGCs coded for uncharacterized T3PKS clusters widely present across 109 species, including *Lactiplantibacillus plantarum* (605 genomes)*, Limosilactobacillus reuteri* (98)*, Pediococcus pentosaceus* (76), and *Oenococcus oeni* (72), among others. This T3PKS coded for the core biosynthetic gene encoding hydroxymethylglutaryl-CoA synthase (Okoye et al., 2022). The second largest family of 638 BGCs coded for a terpene was detected mainly across the *Lactiplantibacillus plantarum* (606) species. Even though there was no similar MIBIG entry for this BGC, it is well known to code for a yellow-colored C(30) carotenoid 4,4’-diaponeurosporene with the antioxidant activity (Garrido-Fernández et al., 2010; Kim et al., 2020).

Various RiPP-like type BGCs consist of unspecified RiPPs in antiSMASH that encode for several bacteriocins such as plantaracins, gassericins, acidocins, penocins, pediocins, and weissellins, among others. As some of these bacteriocins were not included in the MIBIG database, we further mined for them across all BGCs using 229 bacteriocins from the Bactibase database (by using protein blast percent identity of 50%) (Hammami et al., 2010). In total, 1080 BGCs detected across 995 of the genomes had at least one gene with amino acid similarity of greater than 50% with one of the 54 bacteriocins from Bactibase (**Supplementary Figure S56**). Here, we counted only the top hit of the blast PID above 50%. Among these, 478 genomes of *L. plantarum* possessed BGCs to code for bacteriocins plantaricin E, a well-known antimicrobial peptide produced by most of the *L. plantarum* strains, with applications in food preservation and potential therapeutics. Some of the other detected BGCs with known bacteriocins encoded for Enterocin Xbeta (89 genomes from *Lacticaseibacillus spp.*), acidocin J1132 (53 genomes from *Lactobacillus acidophilus* genomes), nisin F (49 genomes from *Ligilactobacillus salivarius*), penocin A (46 genomes from *Pediococcus pentosaceus*), Bacillocin 1580 (38 genomes from *Ligilactobacillus ruminis*), lactacin-F (38 genomes from *Lactobacillus spp.*). Even after annotating BGCs with similarly known bacteriocins, we still find only 1796 of the 6950 BGCs (∼26%) with an assigned known secondary metabolite either from MIBIG database or Bactibase. Thus, a relatively large number of undiscovered BGCs spread acrsoss the rare genome, whereas many of the known BGCs coded for bacteriocins with industrial potential.

## 4. Discussion

In the last few years, characterization of the pangenomes of individual species has demonstrated that the pangenomes represent an important source of knowledge that remains to be extracted. This study expanded this observation to the 26 species of the *Lactobacillaceae* family that are highly important to the food industry. The quality of starting dataset and taxonomic assignments are critical for such a large-scale comparative analysis. In this study, we used high-quality genomes and GTDB-based taxonomic definitions for consistent comparisons. The overall workflow covers many tools in data curation, phylogenetic placement, pangenome reconstruction, functional annotation, genome mining, and detailed pangenome calculations. Together, this workflow led to the discovery of key features of the CAR genomes of the 26 selected species of *Lactobacillaceae*. Particularly, the large number of genes in the rare genomes represents huge genetic diversity across the strains within each species. This finding indicates unique and uncharacterized features for each of the sequenced strains and their potential applications. The combined workflow deployed here will pave the way for future in-depth pangenome analytics for several other bacterial species.

Prior pangenome studies of lactic acid bacteria were focused on a few of the selected species, such as *L. plantarum* (127 genomes) (Carpi et al., 2022)*, L.delbrueckii* (41 genomes) (Kim et al., 2021), *L. acidophilus* (46 genomes) (Huang et al., 2021), *L. paracaseii* (34 genomes) (Smokvina et al., 2013) and *Lactobacillus spp.* (Inglin et al., 2018). The current global analysis of all publicly available genomes finds 26 species eligible for structured pangenome analysis. The re-annotation of taxonomy based on GTDB revealed that some members of Lactobacillaceae were misclassified at NCBI, which is also in accordance with previous findings (Zheng et al., 2020). The pangenomes of most *Lactobacillaceae* species were found to be moderately open compared to other bacterial species. The vast genome diversity of the species was also discussed in prior pangenome studies of a few of the *Lactobacillaceae* (Ksiezarek et al., 2022; Zhang et al., 2020). Thus the larger size of the rare genomes observed in this study represents an important marker for characterizing the genomic diversity of lactic acid bacteria.

The COG-based enrichment analysis showed that the core genomes among the 26 species were primarily enriched with translational and post-translational related functions. Previously, the translation-related genes were also reported to be core in the *Lactobacillaceae* gene pool (Kant et al., 2011) and 12 different microbial species (Hyun et al., 2022). Interestingly, two important species, *L. helveticus* and *L. ruminis,* found in the human gut, possess the genes of T4PS and flagella formation (*flgJ*). T4PS is responsible for motility and attachment to epithelial surfaces and has a significant role in pathogenicity (Lieberman et al., 2015). Likewise, flagella also play an important role in pathogenicity and help the bacteria to reach different sites in the host (Herlihey et al., 2014). Further, the genes within the accessory genomes show enrichment in metabolism-related functions such as amino acid and carbohydrate metabolism. The accessory genomes also included genes from the Type I restriction-modification system and ABC-transport system specific to antimicrobial peptides and multi-drugs, which explains phage resistance (Suárez et al., 2009) and antibiotic resistance in *Lactobacillus* species (Orelle et al., 2019). Therefore, the COG enrichments suggest that the core and accessory genomes possess functionally diverse genes.

At the outset, we expected to observe the most closed and open pangenome in niche specialized and nomadic species, respectively. However, the *Limosilactobacillus reuteri* (mainly isolated from the GI tract) had the most open pangenome within our compendium, whereas *Lactobacillus acidophilus* (isolated from several different niches including the GI tract, dairy products, plant-based foods, and a few environmental isolates) had the most closed pangenome. For a better understanding of diversification forces within *Lactobacillaceae*, we investigated the abundance and dispersion of genes related to the mobilome (transposable elements and prophages) and DNA cleavage/recombination activity. These genes were reported to actively contribute to horizontal gene transfer (HGT) and gene loss due to their mobile and replicative nature (Darmon and Leach, 2014). Likewise, we found that hypothetical proteins, which are commonly known as pseudogenes, are collocated with TEs. This collocation may result from the TEs gene disruption activity. Here, we discovered a correlation between the abundance or genomic dispersion of these elements and pangenome openness. Therefore, the openness of a pangenome is likely to be correlated to the mobilome and genome instability rather than ecological niches. Future investigations into this hypothesis may help elucidate key evolutionary forces behind *Lactobacillaceae* diversification.

The diversity of the pangenomes at the species level indicates the need for analyzing species at a finer phylogroup level. Here, the Mash-based distances across whole genome sequences of the *L. plantarum* pangenome with 611 strains were used to identify nine distinct phylogroup candidates, one more genomically divergent than the others (Carpi et al., 2022). A COG category distribution across various phylogroups revealed phylogroup-specific functions. Of the 1,146 genes exclusive to this divergent phylogroup (Cluster A), 28.7% did not map to any functionally known gene orthologs, suggesting a highly underexplored functionality of the phylogroup. Many known genes carry out functions like replication and repair, cell wall/membrane/envelope biogenesis, transcription, carbohydrate metabolism, and amino acid metabolism. A Fisher’s Exact Test on the exclusive genes revealed that these 29 strains, which make up this phylogroup, also had their own set of isozymes responsible for histidine biosynthesis, separate from the genes which code for the same pathway in the other strains. The detailed analysis of phylogroup-specific genes further supports the need for looking at different phylogroups within species for better classification.

Secondary metabolites play an important role in microbial defense along with other properties. In the *Lactobacillaceae* family, several organisms produce specific bacteriocins, which are small peptides that are ribosomally synthesized and post-translationally modified (RiPPs). In this study, we found that various species of the *Lactobacillaceae* family had specific bacteriocins. For example, most of the *Lactiplantibacillus plantarum* genomes contained plantaricin-encoding BGCs. These species-specific bacteriocins give them specific antimicrobial properties that play a significant role in food preservation e.g. plantaricins (Choi et al., 2021; Kawahara et al., 2022). Apart from bacteriocins, a larger family of T3PKS BGCs was observed across most species, encoding the enzyme hydroxymethylglutaryl-CoA synthase. Additionally, terpene-type BGC coding for carotenoid 4,4’-diaponeurosporene was abundantly present in *Lactiplantibacillus plantarum,* which has antioxidant properties. We also found that many of the detected BGC families had no known secondary metabolites, thus highlighting a novel potential for their bioactivity. The high number of unique BGCs further represent characteristics of the rare genome and are potentially considered signatures for individual strains.

## 5. Conclusion

Going forward, extracting the pangenomic characteristics of bacterial species will be an important method for providing insights into evolution and adaptation and the potential for genetic diversity in the group. Our analysis suggests that most *Lactobacillaceae* species have a moderately open pangenome with larger rare genome sizes, reflecting the diversity and high probability of finding novel functional genes. Further, the openness of pangenomes could be correlated with many mobile genetic elements, which pave the way for studying HGT events and their impact on genomic diversity. Additionally, the nine phylogroups identified based on the genomic similarity within the *L. plantarum* species show the need for finer resolution of species analysis. The rare BGCs in the rare genome represent the unique bioactive potential for each strain in food preservatives, probiotics, and antimicrobials, among other applications. In summary, using a comprehensive collection of available sequences, our study represents the first comprehensive comparative pangenome analysis of 26 identifiable species in the *Lactobacillaceae* family.

## Acknowledgments

We thank Marc Abrams for reviewing the manuscript and providing constructive suggestions. We thank Matin Nuhamunada for providing inputs on the Snakemake workflow used for data processing.

