## Supplementary Figure for "Pangenome analysis reveals the genetic basis for taxonomic classification of the Lactobacillaceae family"

### Supplementary Figures

***Supplementary Figure S1. Sankey diagram representing reclassification of genera from NCBI to GTDB taxonomy***

***Supplementary Figure S2: Phylogenetic tree of 307 representative genomes from different species across 33 genera.*** *The top 10 genera are represented by different colors whereas grey represents other genera with the text denoting the number of other genera in each phylogenetic clade (total of 33 genera). The bars represent the genome length.*

***Supplementary Figure S3: A clustermap of Mash distances for all filtered Lactobacillaceae strains (3,591 total). Clusters are generally species or genus-specific, with some clusters being a composition of 2 or more different genera or species.***

***Supplementary Figure S4: Gene CDF plot for Lacticaseibacillus paracasei.***
***Supplementary Figure S5: Gene CDF plot for Lacticaseibacillus rhamnosus.***
***Supplementary Figure S6: Gene CDF plot for Lactiplantibacillus pentosus.***
***Supplementary Figure S7: Gene CDF plot for Lactiplantibacillus plantarum.***
***Supplementary Figure S8: Gene CDF plot for Lactobacillus acdiophillus.***
***Supplementary Figure S9: Gene CDF plot for Lactobacillus crispatus.***
***Supplementary Figure S10: Gene CDF plot for Lactobacillus delbrueckii.***
***Supplementary Figure S11: Gene CDF plot for Lactobacillus gasseri.***
***Supplementary Figure S12: Gene CDF plot for Lactobacillus helveticus.***
***Supplementary Figure S13: Gene CDF plot for Lactobacillus iners.***
***Supplementary Figure S14: Gene CDF plot for Lactobacillus johnsonii.***
***Supplementary Figure S15: Gene CDF plot for Lactobacillus paragasseri.***
***Supplementary Figure S16: Gene CDF plot for Latilactobacillus sakei.***
***Supplementary Figure S17: Gene CDF plot for Lentilactobacillus parabuchneri.***
***Supplementary Figure S18: Gene CDF plot for Leuconostoc inhae.***
***Supplementary Figure S19: Gene CDF plot for Leuconostoc mesenteroides.***
***Supplementary Figure S20: Gene CDF plot for Levilactobacillus brevis.***
***Supplementary Figure S21: Gene CDF plot for Ligilactobacillus ruminis.***
***Supplementary Figure S22: Gene CDF plot for Ligilactobacillus salivarius.***
***Supplementary Figure S23: Gene CDF plot for Limosilactobacillus fermentum.***
***Supplementary Figure S24: Gene CDF plot for Limosilactobacillus reuteri.***
***Supplementary Figure S25: Gene CDF plot for Oenococcus oeni.***
***Supplementary Figure S26: Gene CDF plot for Pediococcus acidilactici.***
***Supplementary Figure S27: Gene CDF plot for Pediococcus pentosaceus.***
***Supplementary Figure S28: Gene CDF plot for Weissella cibaria.***
***Supplementary Figure S29: Gene CDF plot for Weissella confusa.***

***Supplementary Figure S30: The barplot depicts the COG functions among core, accessory, and rare pangenome in Lacticaseibacillus paracasei. The barplot is between***

***the number of genes vs COG functions. Note: " - " refers to genes with no COG annotation.***

***Supplementary Figure S31: The barplot depicts the COG functions among core, accessory, and rare pangenome in Lacticaseibacillus rhamnosus. The barplot is between the number of genes vs COG functions. Note: " - " refers to genes with no COG annotation.***

***Supplementary Figure S32: The barplot depicts the COG functions among core, accessory, and rare pangenome in Lactiplantibacillus pentosus. The barplot is between the number of genes vs COG functions. Note: " - " refers to genes with no COG annotation.***

***Supplementary Figure S33: The barplot depicts the COG functions among core, accessory, and rare pangenome in Lactobacillus acdiophillus. The barplot is between the number of genes vs COG functions. Note: " - " refers to genes with no COG annotation.***

***Supplementary Figure S34: The barplot depicts the COG functions among core, accessory, and rare pangenome in Lactobacillus crispatus. The barplot is between the number of genes vs COG functions. Note: " - " refers to genes with no COG annotation.***

***Supplementary Figure S35: The barplot depicts the COG functions among core, accessory, and rare pangenome in Lactobacillus delbrueckii. The barplot is between the number of genes vs COG functions. Note: " - " refers to genes with no COG annotation.***

***Supplementary Figure S36: The barplot depicts the COG functions among core, accessory, and rare pangenome in Lactobacillus gasseri. The barplot is between the number of genes vs COG functions. Note: " - " refers to genes with no COG annotation.***

***Supplementary Figure S37: The barplot depicts the COG functions among core, accessory, and rare pangenome in Lactobacillus helveticus. The barplot is between the number of genes vs COG functions. Note: " - " refers to genes with no COG annotation.***

***Supplementary Figure S38: The barplot depicts the COG functions among core, accessory, and rare pangenome in Lactobacillus iners. The barplot is between the number of genes vs COG functions. Note: " - " refers to genes with no COG annotation.***

***Supplementary Figure S39: The barplot depicts the COG functions among core, accessory, and rare pangenome in Lactobacillus johnsonii. The barplot is between the number of genes vs COG functions. Note: " - " refers to genes with no COG annotation.***

*Supplementary Figure S40: The barplot depicts the COG functions among core, accessory, and rare pangenome in Lactobacillus paragasseri. The barplot is between the number of genes vs COG functions. Note: “ - ” refers to genes with no COG annotation.*

*Supplementary Figure S41: The barplot depicts the COG functions among core, accessory, and rare pangenome in Latilactobacillus sakei. The barplot is between the number of genes vs COG functions. Note: “ - ” refers to genes with no COG annotation.*

*Supplementary Figure S42: The barplot depicts the COG functions among core, accessory, and rare pangenome in Lentilactobacillus parabuchneri. The barplot is between the number of genes vs COG functions. Note: “ - ” refers to genes with no COG annotation.*

*Supplementary Figure S43: The barplot depicts the COG functions among core, accessory, and rare pangenome in Leuconostoc inhae. The barplot is between the number of genes vs COG functions. Note: “ - ” refers to genes with no COG annotation.*

*Supplementary Figure S44: The barplot depicts the COG functions among core, accessory, and rare pangenome in Leuconostoc mesenteroides. The barplot is between the number of genes vs COG functions. Note: “ - ” refers to genes with no COG annotation.*

*Supplementary Figure S45: The barplot depicts the COG functions among core, accessory, and rare pangenome in Levilactobacillus brevis. The barplot is between the number of genes vs COG functions. Note: “ - ” refers to genes with no COG annotation.*

*Supplementary Figure S46: The barplot depicts the COG functions among core, accessory, and rare pangenome in Ligilactobacillus ruminis. The barplot is between the number of genes vs COG functions. Note: “ - ” refers to genes with no COG annotation.*

*Supplementary Figure S47: The barplot depicts the COG functions among core, accessory, and rare pangenome in Ligilactobacillus salivarius. The barplot is between the number of genes vs COG functions. Note: “ - ” refers to genes with no COG annotation.*

*Supplementary Figure S48: The barplot depicts the COG functions among core, accessory, and rare pangenome in Limosilactobacillus fermentum. The barplot is between the number of genes vs COG functions. Note: “ - ” refers to genes with no COG annotation.*

*Supplementary Figure S49: The barplot depicts the COG functions among core, accessory, and rare pangenome in Limosilactobacillus reuteri. The barplot is between the number of genes vs COG functions. Note: “ - ” refers to genes with no COG annotation.*

***Supplementary Figure S50:** The barplot depicts the COG functions among core, accessory, and rare pangenome in Oenococcus oeni. The barplot is between the number of genes vs COG functions. Note: " - " refers to genes with no COG annotation.*

***Supplementary Figure S51:** The barplot depicts the COG functions among core, accessory, and rare pangenome in Pediococcus acidilactici. The barplot is between the number of genes vs COG functions. Note: " - " refers to genes with no COG annotation.*

***Supplementary Figure S52:** The barplot depicts the COG functions among core, accessory, and rare pangenome in Pediococcus pentosaceus. The barplot is between the number of genes vs COG functions. Note: " - " refers to genes with no COG annotation.*

***Supplementary Figure S53:** The barplot depicts the COG functions among core, accessory, and rare pangenome in Weissella cibaria. The barplot is between the number of genes vs COG functions. Note: " - " refers to genes with no COG annotation.*

***Supplementary Figure S54:** The barplot depicts the COG functions among core, accessory, and rare pangenome in Weissella confusa. The barplot is between the number of genes vs COG functions. Note: " - " refers to genes with no COG annotation.*

***Supplementary Figure S55:** A clustermap of Mash distances for all L. plantarum strains with the phylogenetically distinct Mash Cluster A removed. All the clusters remain stable, with Cluster E splitting into 4 subclusters and Cluster I splitting into 3 subclusters.*

***Supplementary Figure S56:** Biosynthetic gene clusters with BLAST hits (50%) in Bactibase.*

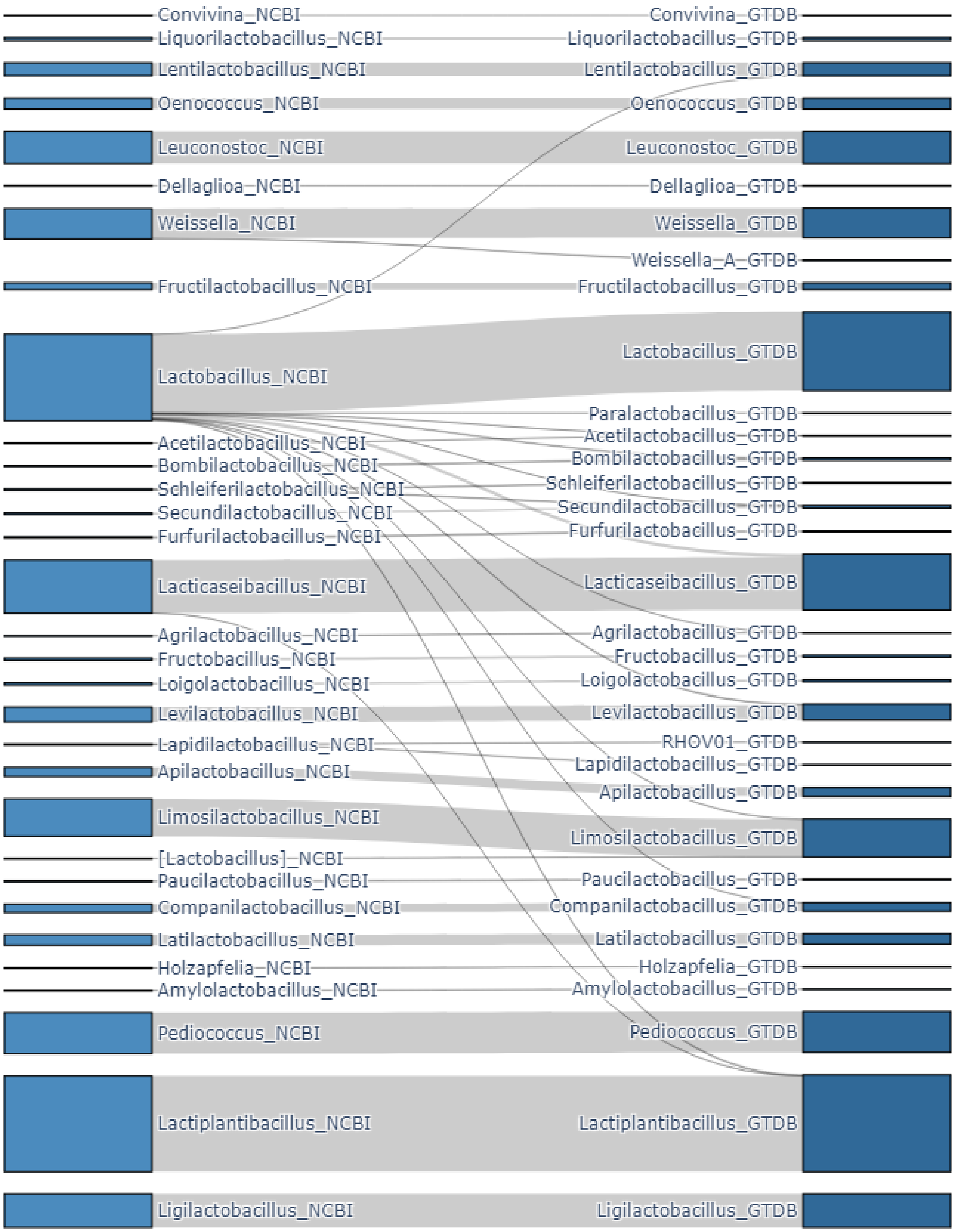


***Supplementary Figure S1. Sankey diagram representing reclassification of genera from NCBI to GTDB taxonomy***

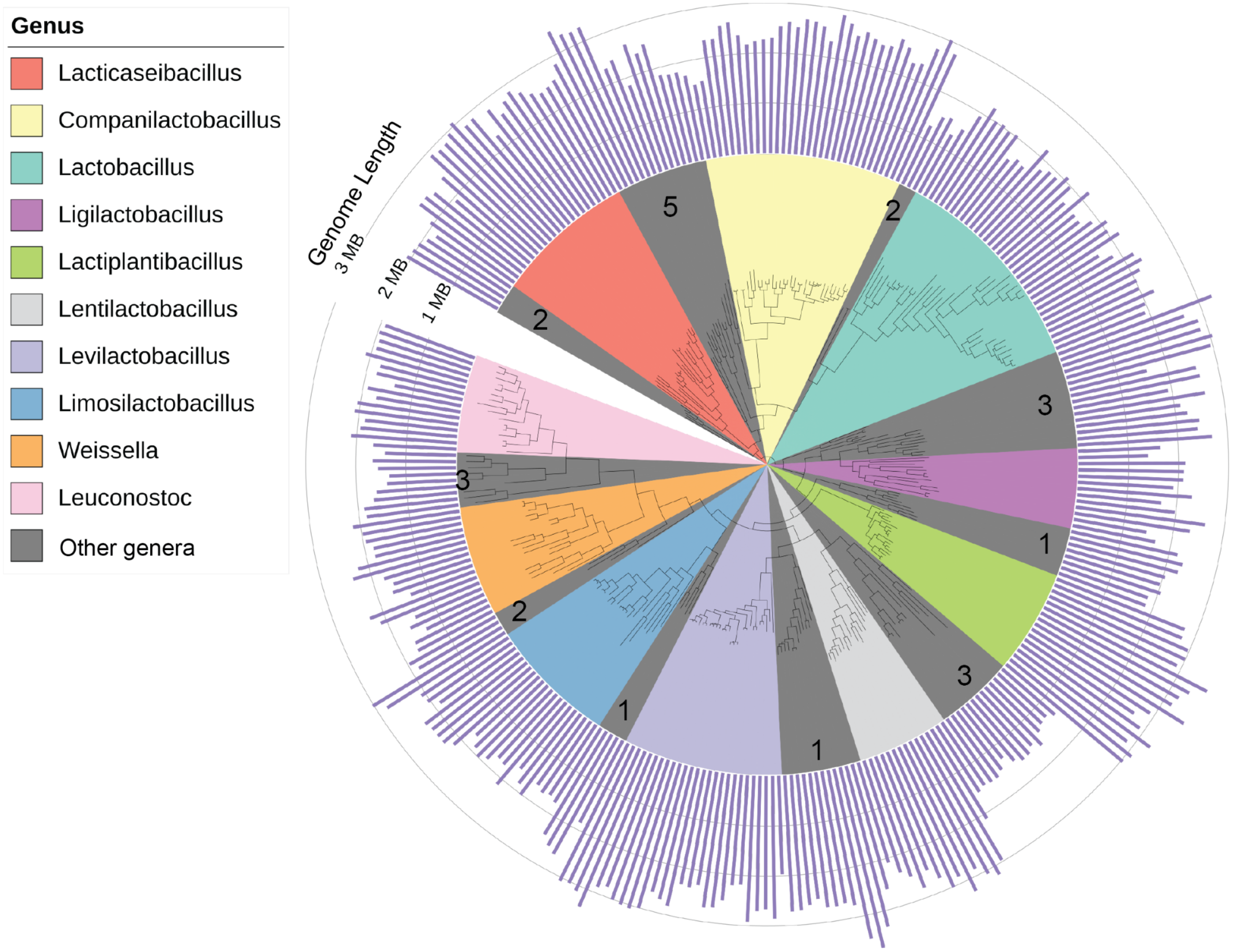


***Supplementary Figure S2: Phylogenetic tree of 307 representative genomes from different species across 33 genera.*** *The top 10 genera are represented by different colors whereas grey represents other genera with the text denoting the number of other genera in each phylogenetic clade (total of 33 genera). The bars represent the genome length.*

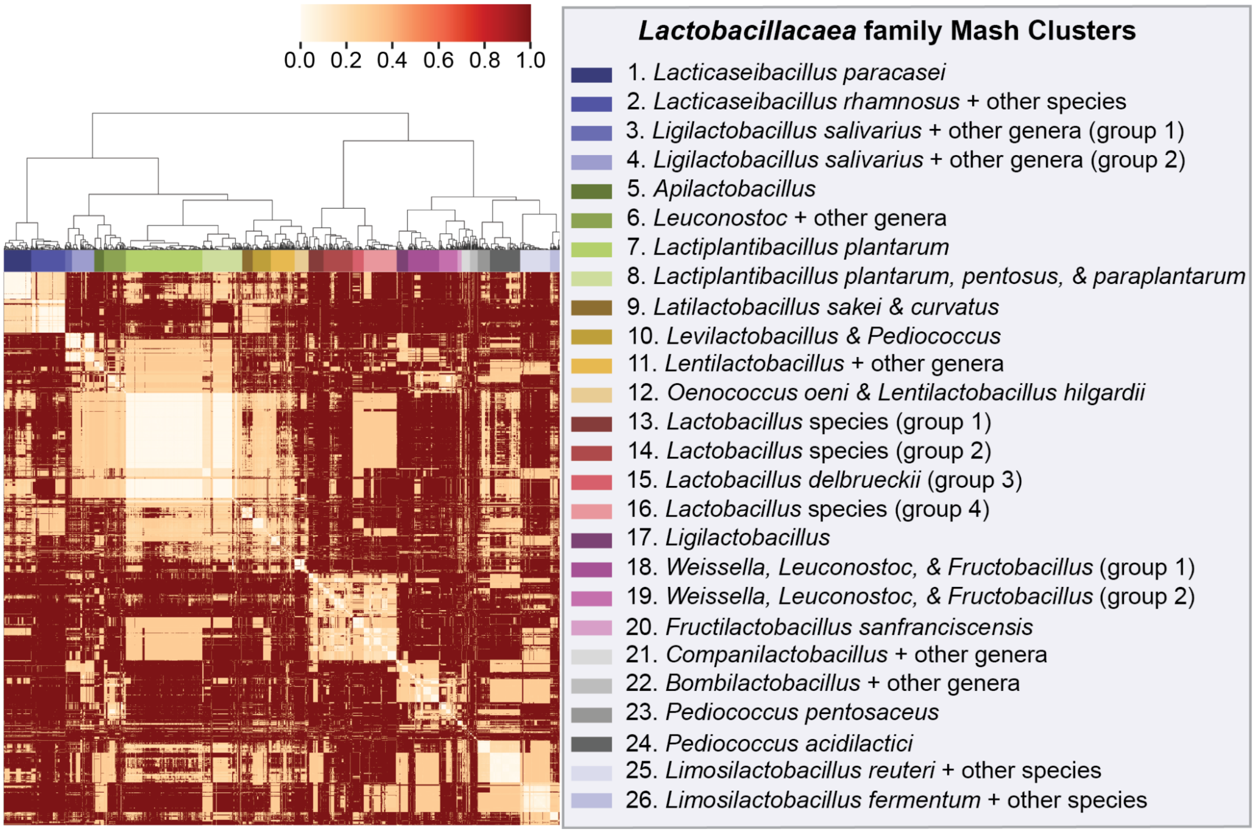


***Supplementary Figure S3: A clustermap of Mash distances for all filtered Lactobacillaceae strains (3,591 total). Clusters are generally species or genus-specific, with some clusters being a composition of 2 or more different genera or species.***

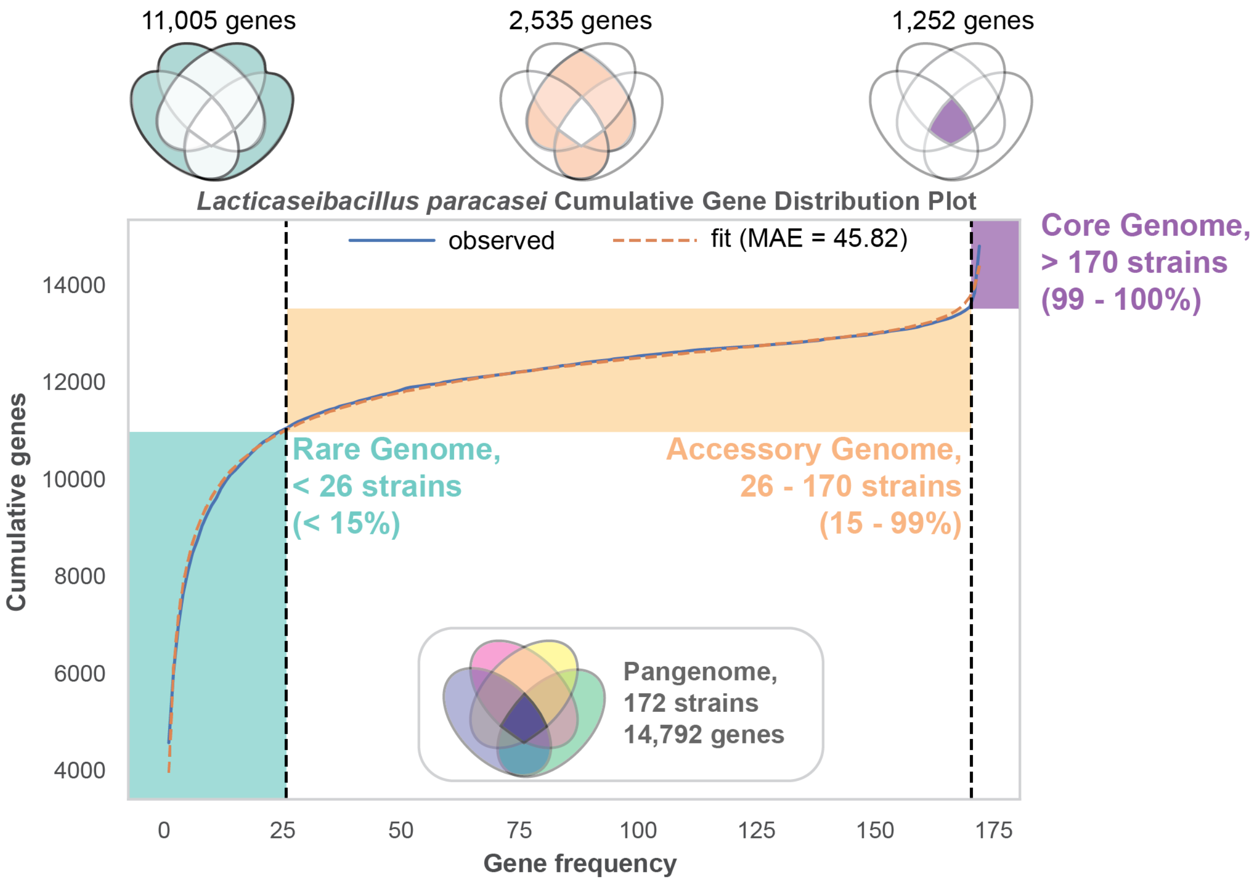


***Supplementary Figure S4: Gene CDF plot for Lacticaseibacillus paracasei.***

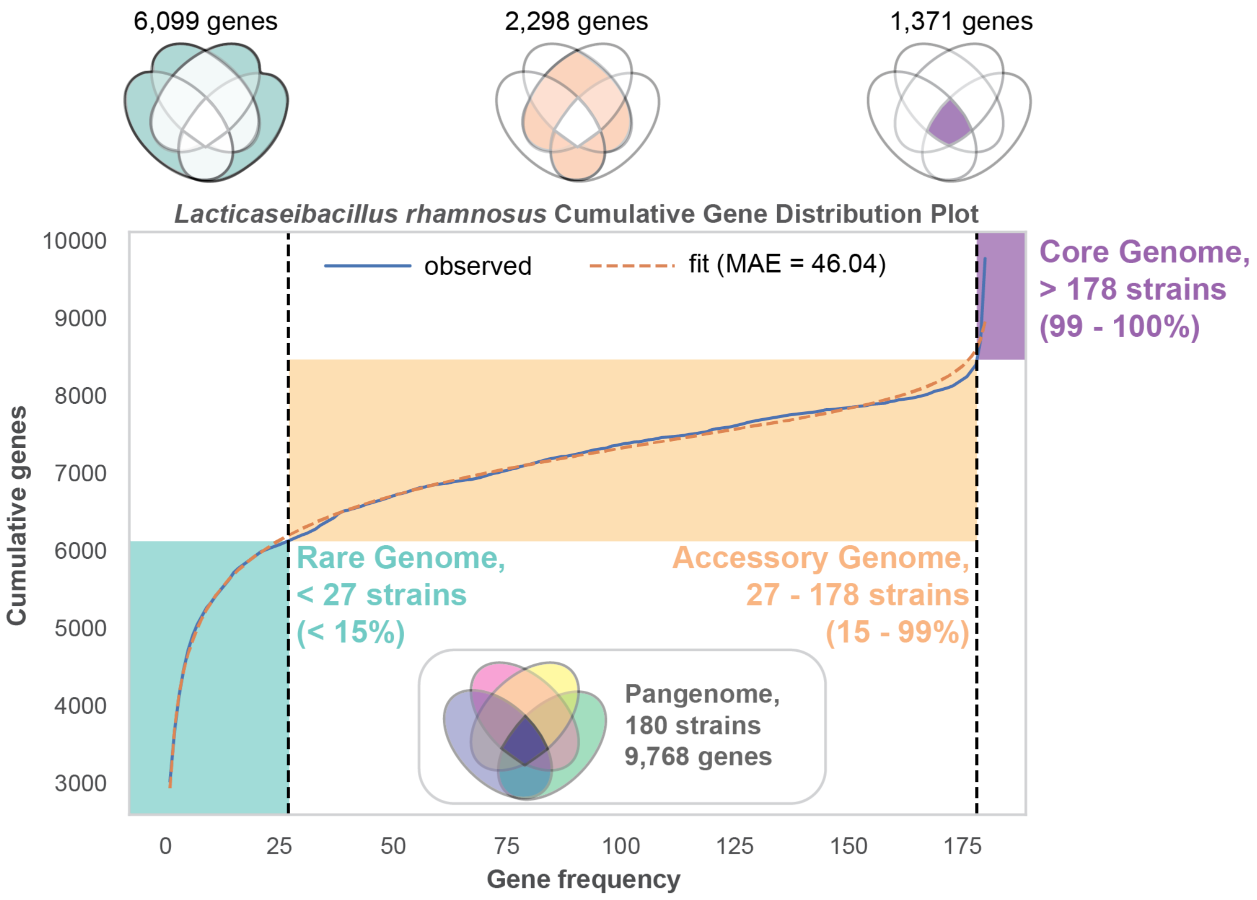


***Supplementary Figure S5: Gene CDF plot for Lacticaseibacillus rhamnosus.***

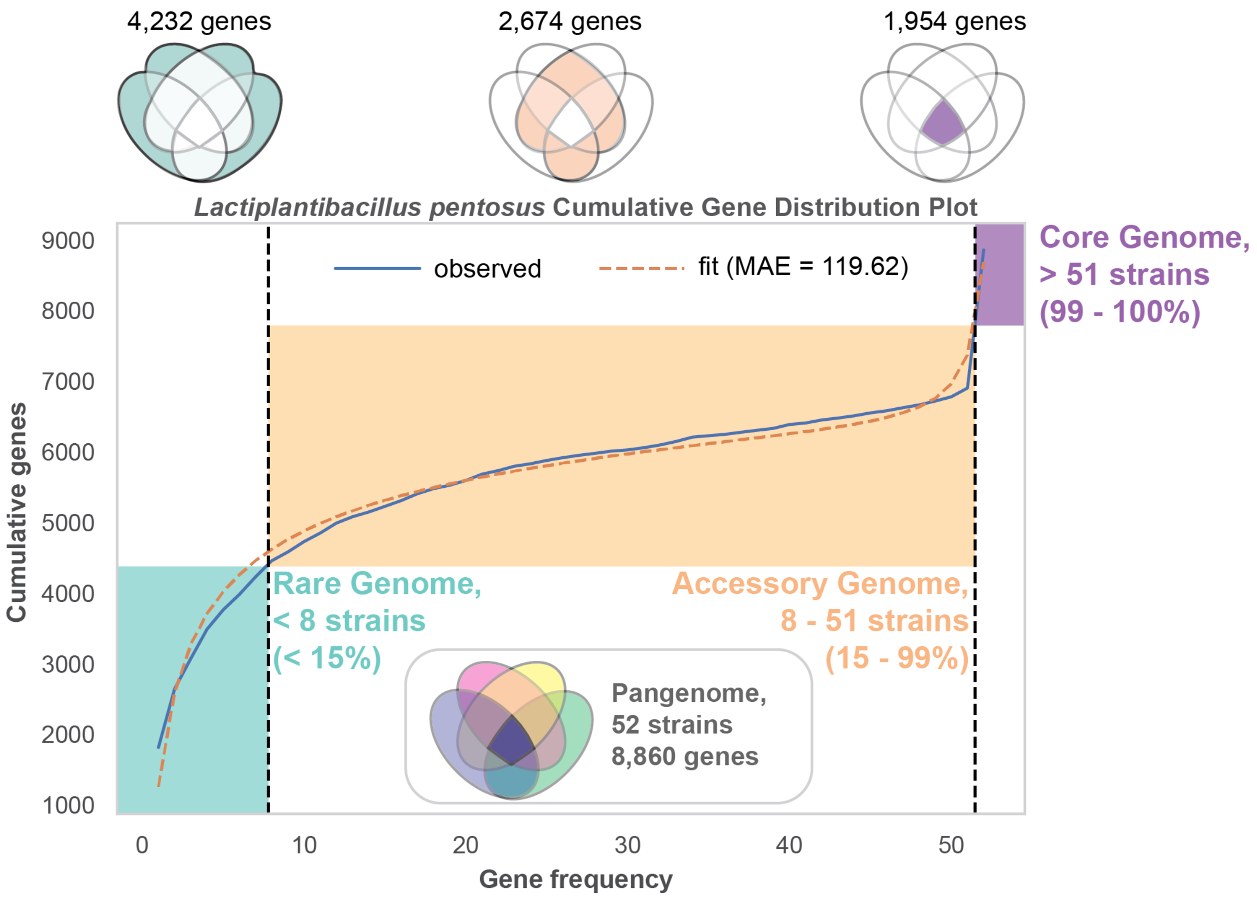


***Supplementary Figure S6: Gene CDF plot for Lactiplantibacillus pentosus.***

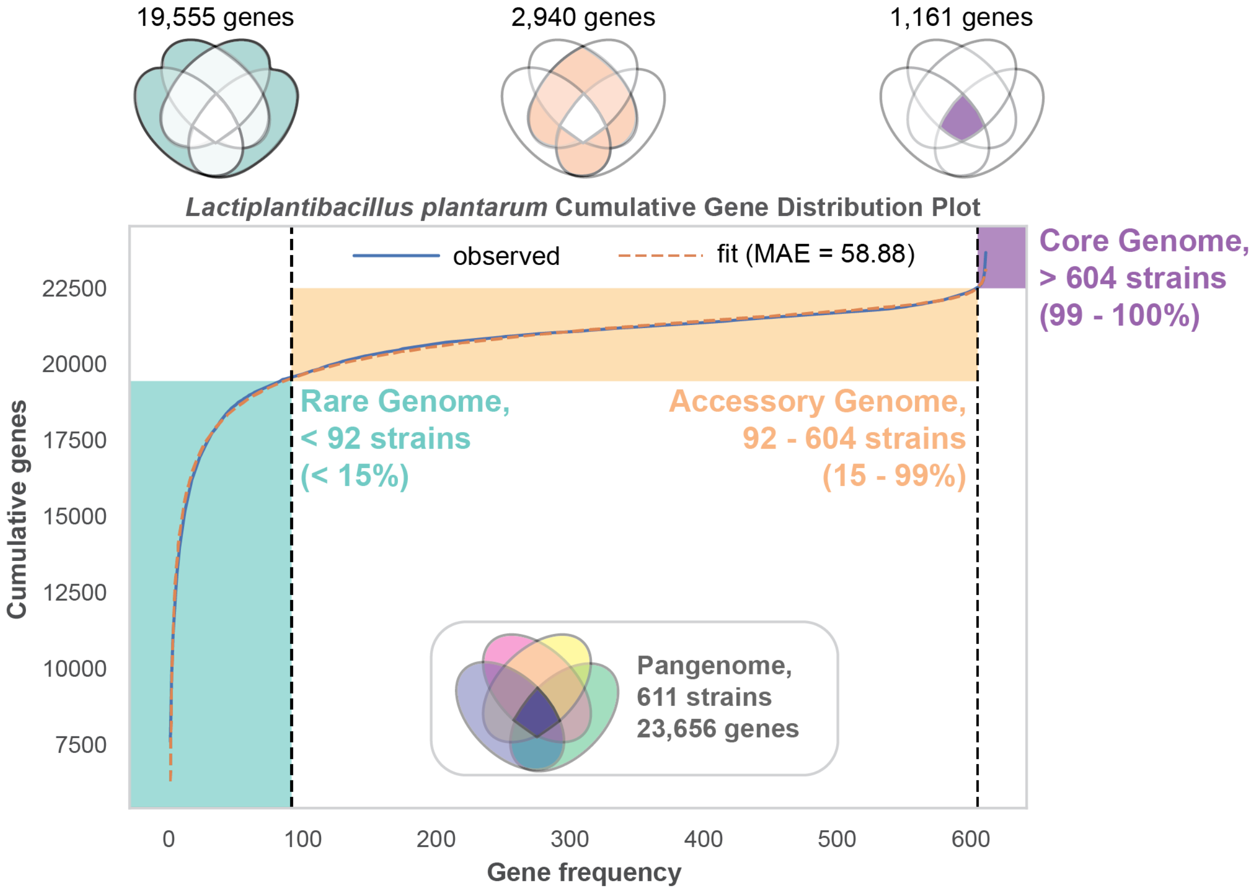


***Supplementary Figure S7: Gene CDF plot for Lactiplantibacillus plantarum.***

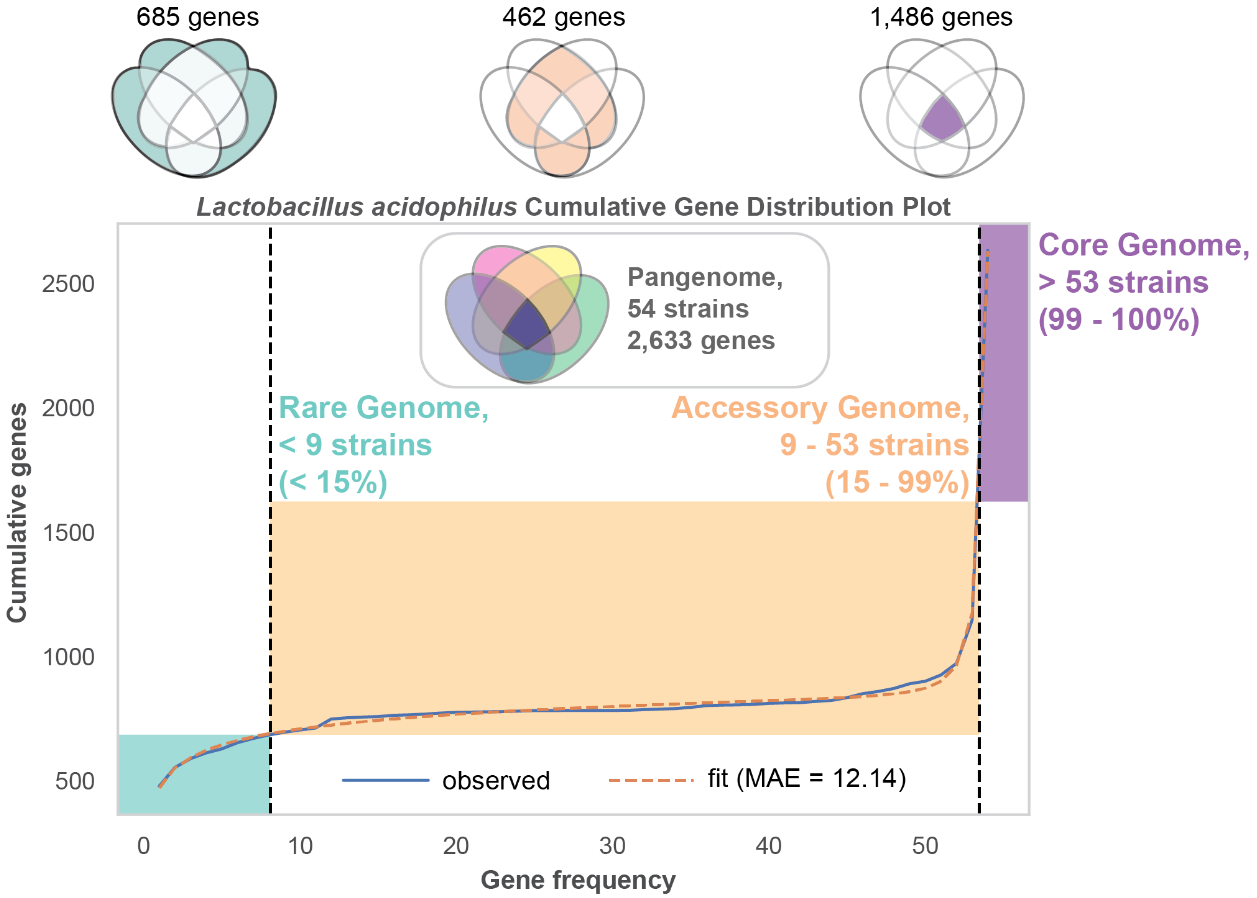


***Supplementary Figure S8: Gene CDF plot for Lactobacillus acdiophillus.***

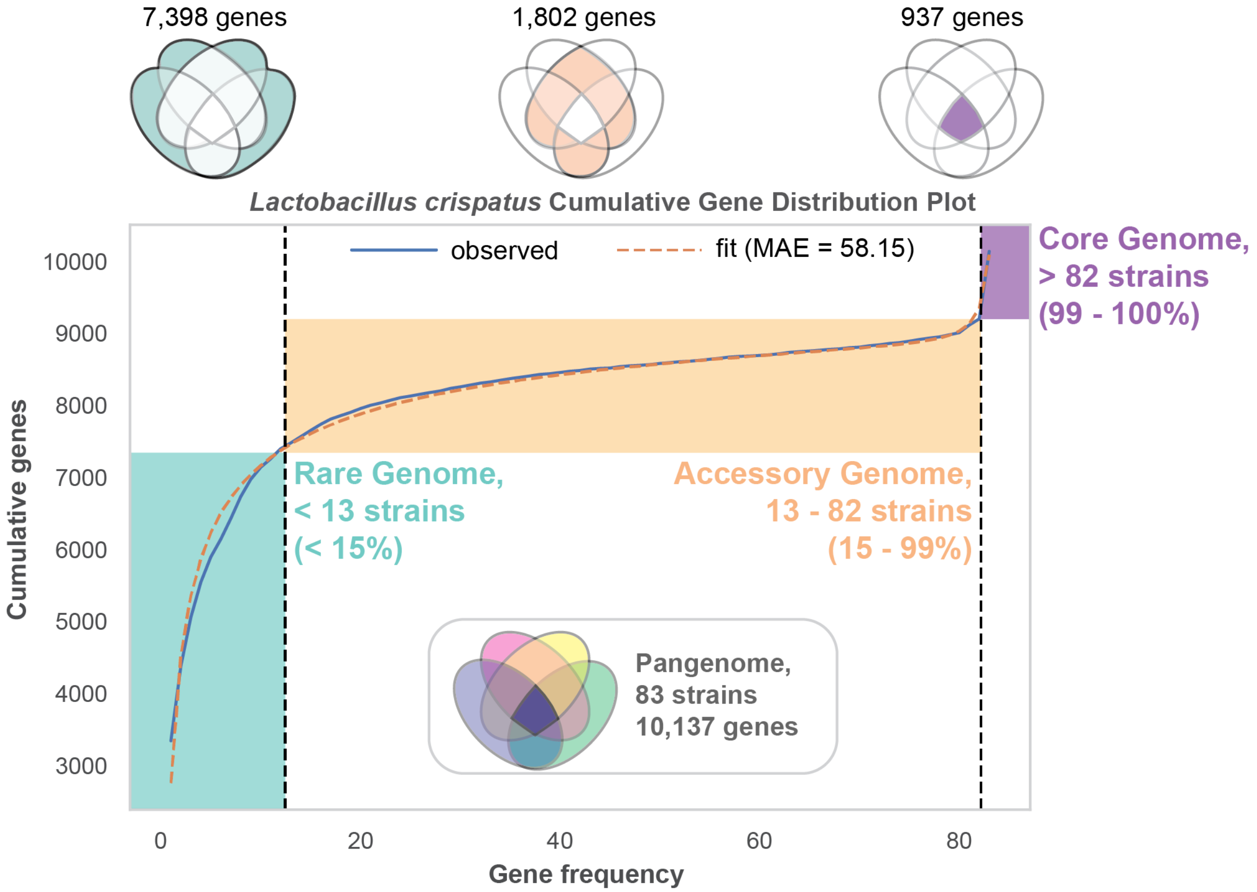


***Supplementary Figure S9: Gene CDF plot for Lactobacillus crispatus.***

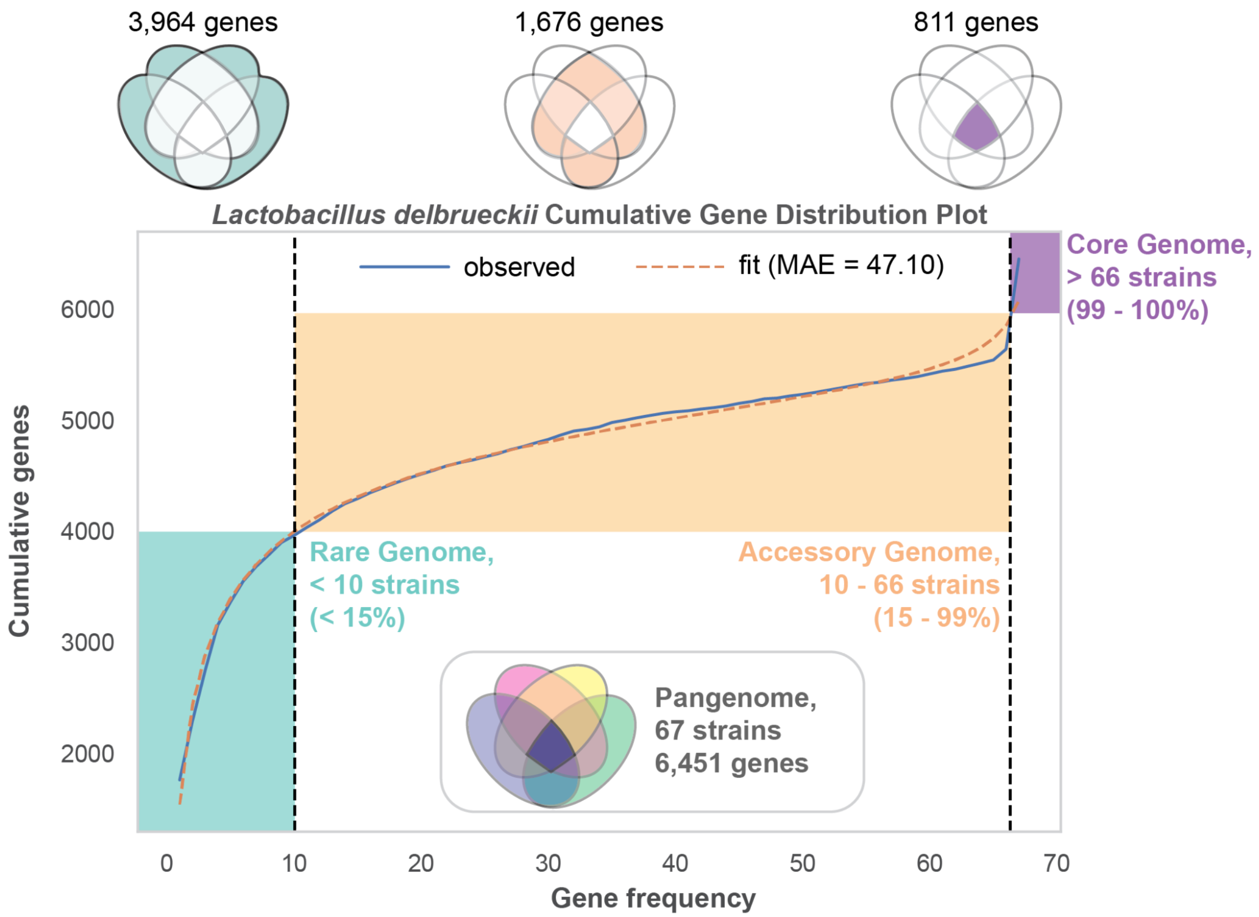


***Supplementary Figure S10: Gene CDF plot for Lactobacillus delbrueckii.***

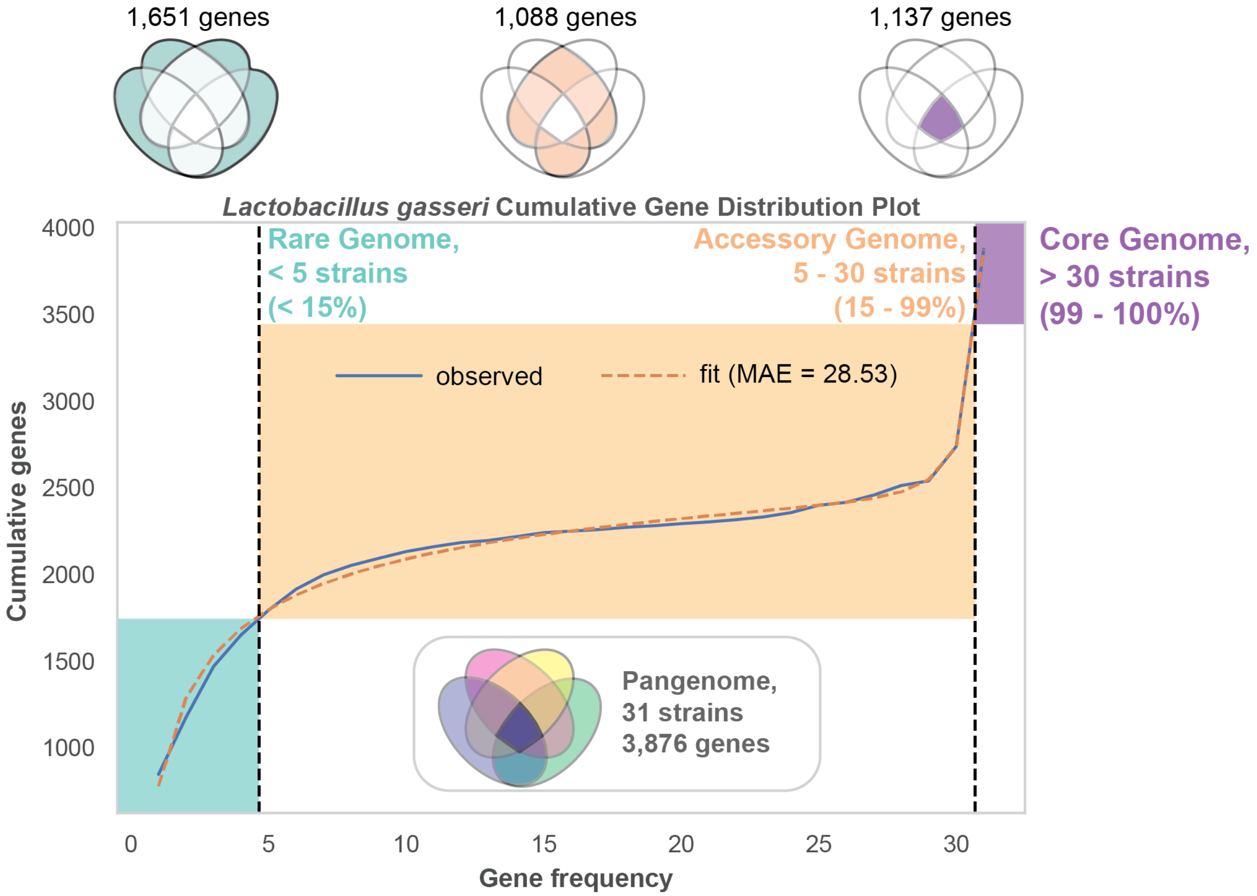


***Supplementary Figure S11: Gene CDF plot for Lactobacillus gasseri.***

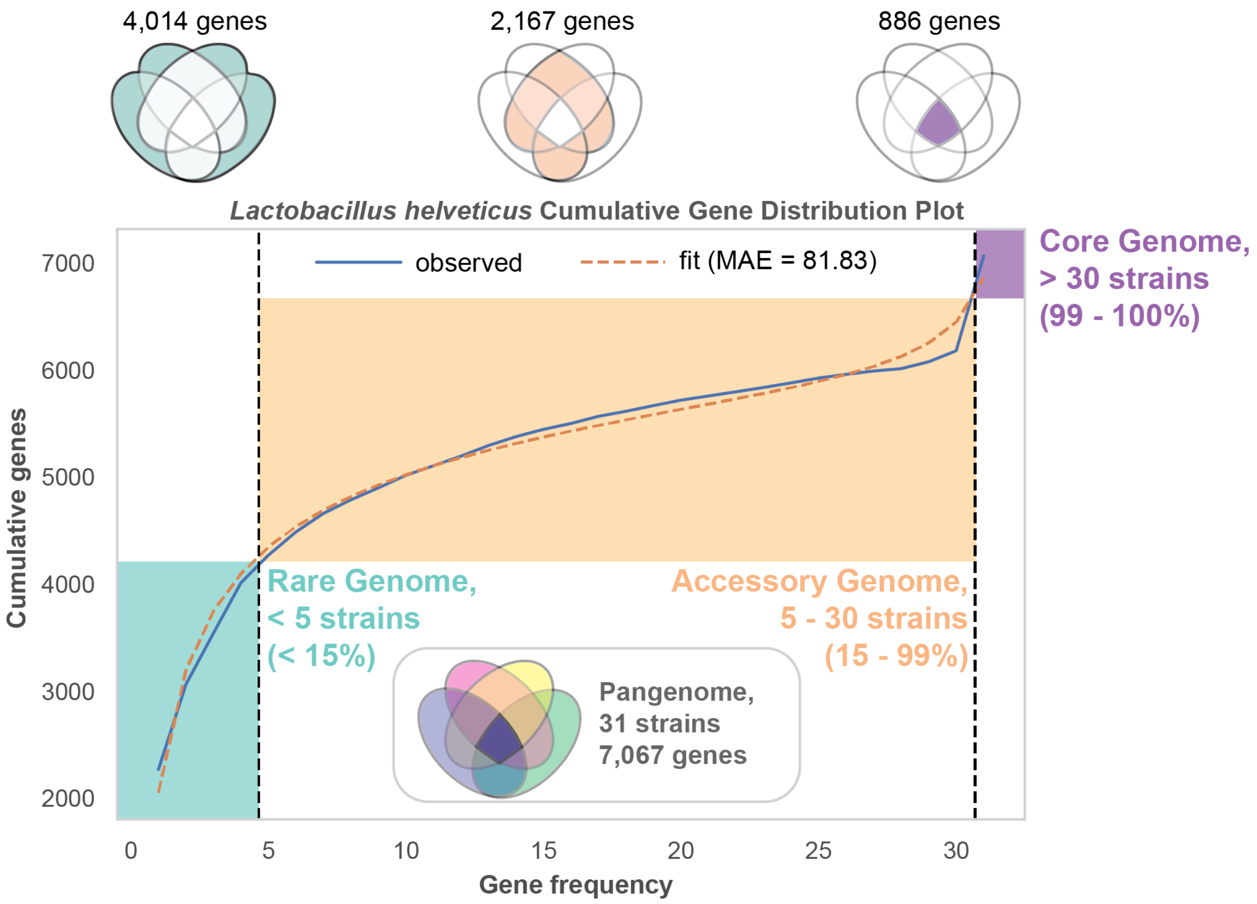


***Supplementary Figure S12: Gene CDF plot for Lactobacillus helveticus.***

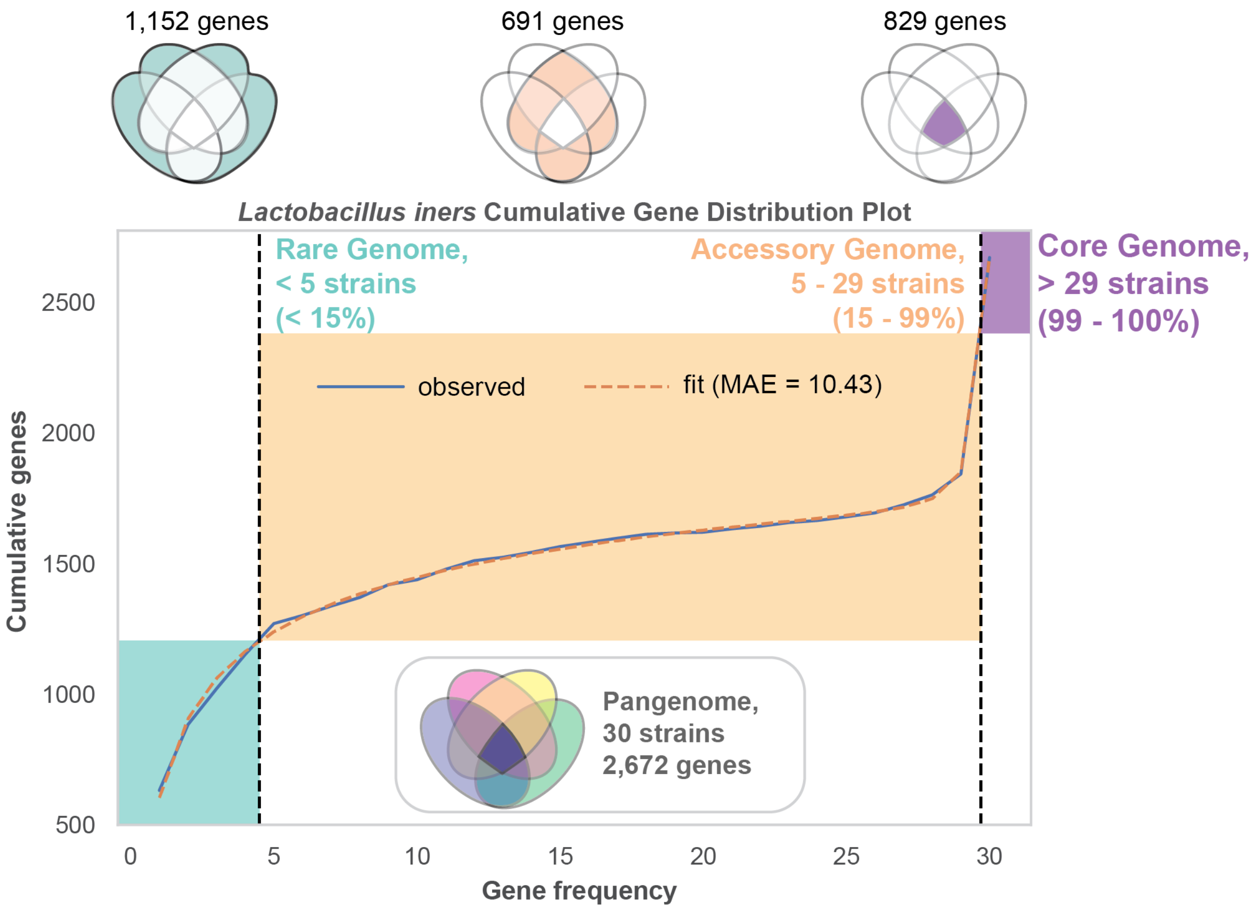


***Supplementary Figure S13: Gene CDF plot for Lactobacillus iners.***

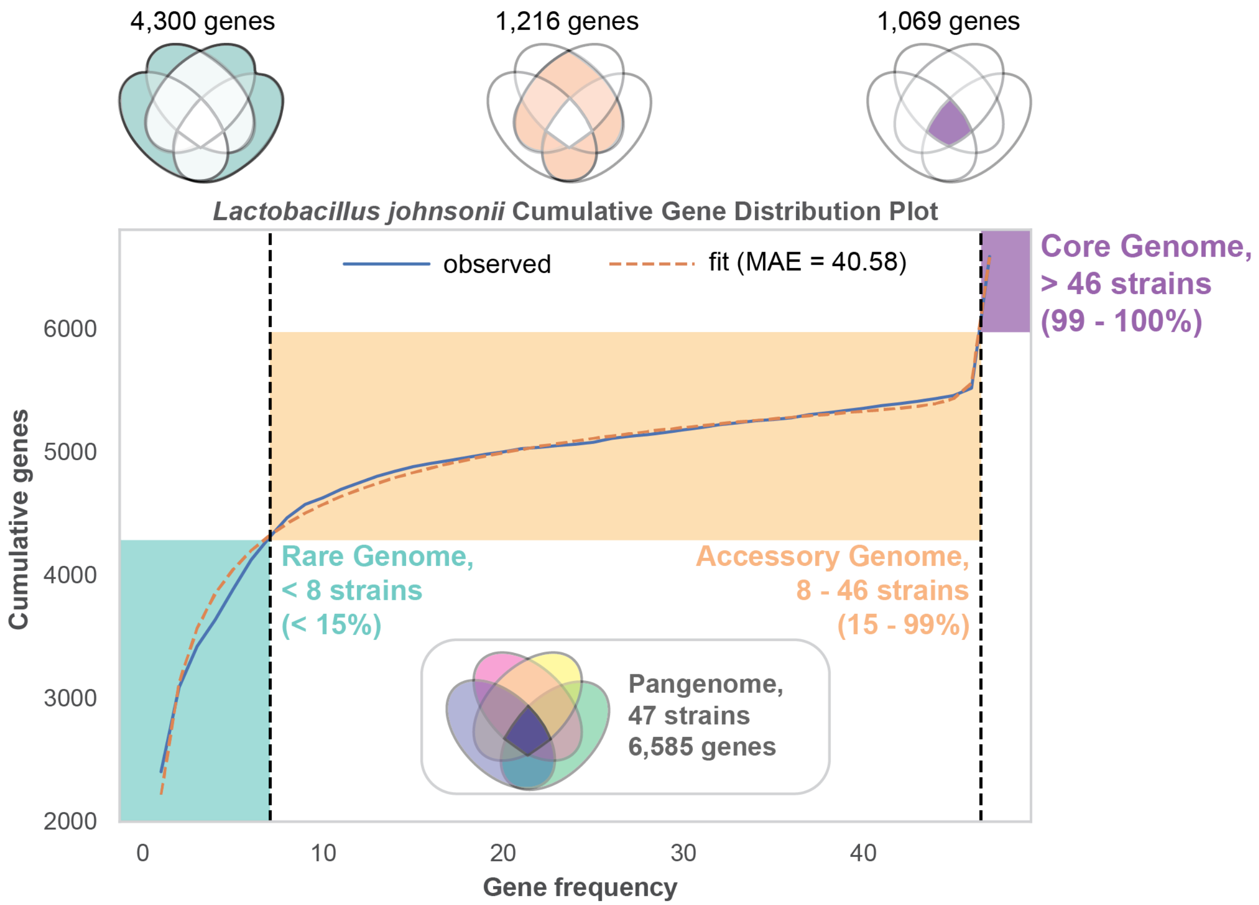


***Supplementary Figure S14: Gene CDF plot for Lactobacillus johnsonii.***

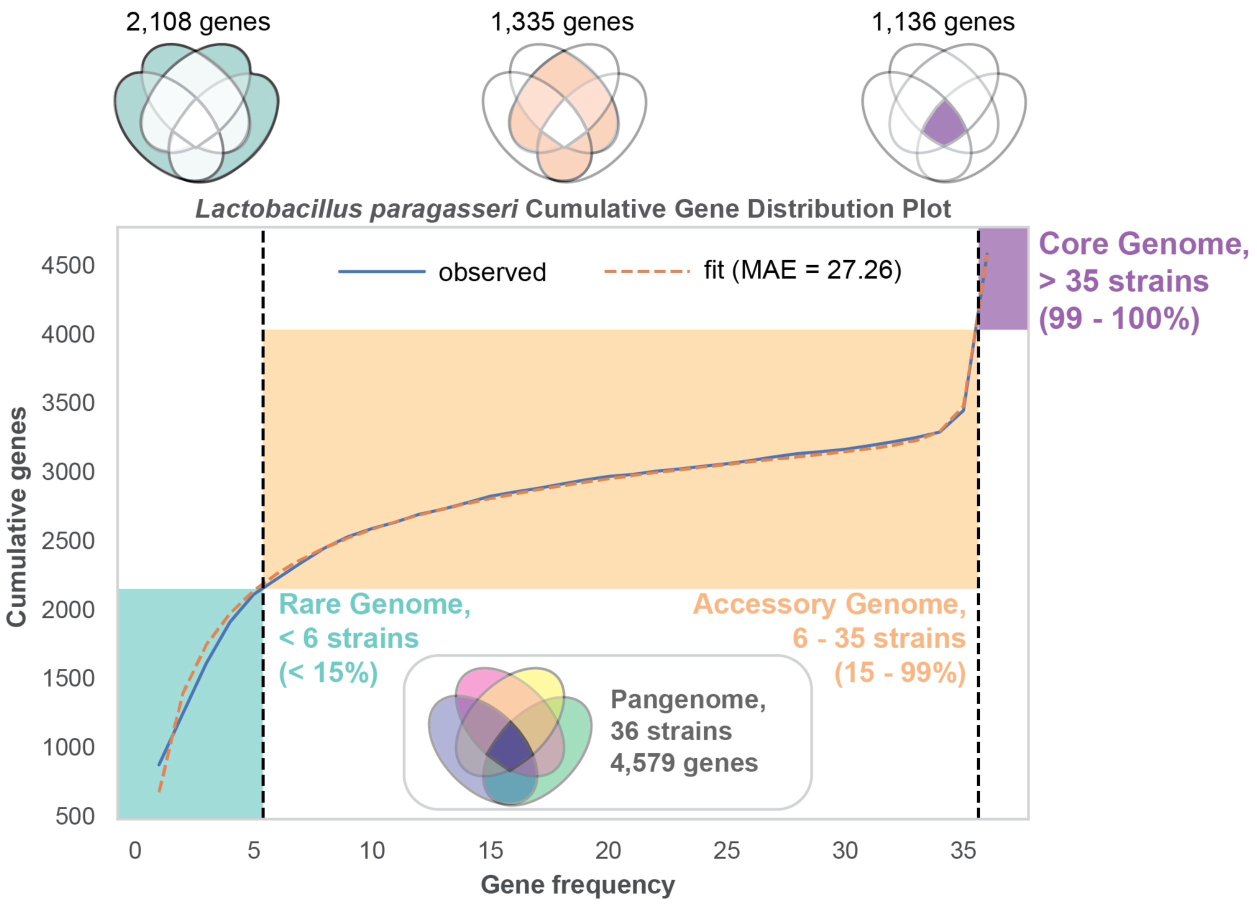


***Supplementary Figure S15: Gene CDF plot for Lactobacillus paragasseri.***

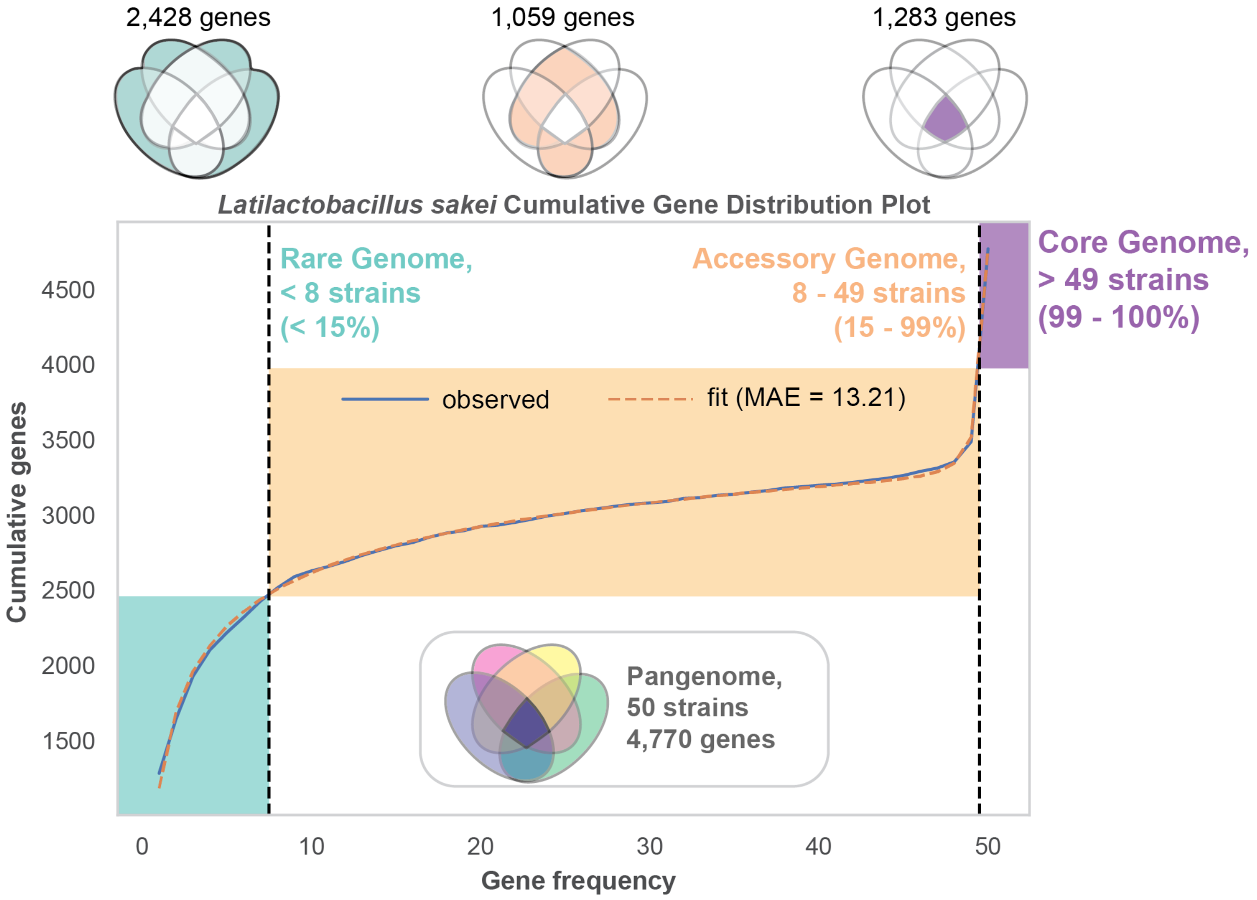


***Supplementary Figure S16: Gene CDF plot for Latilactobacillus sakei.***

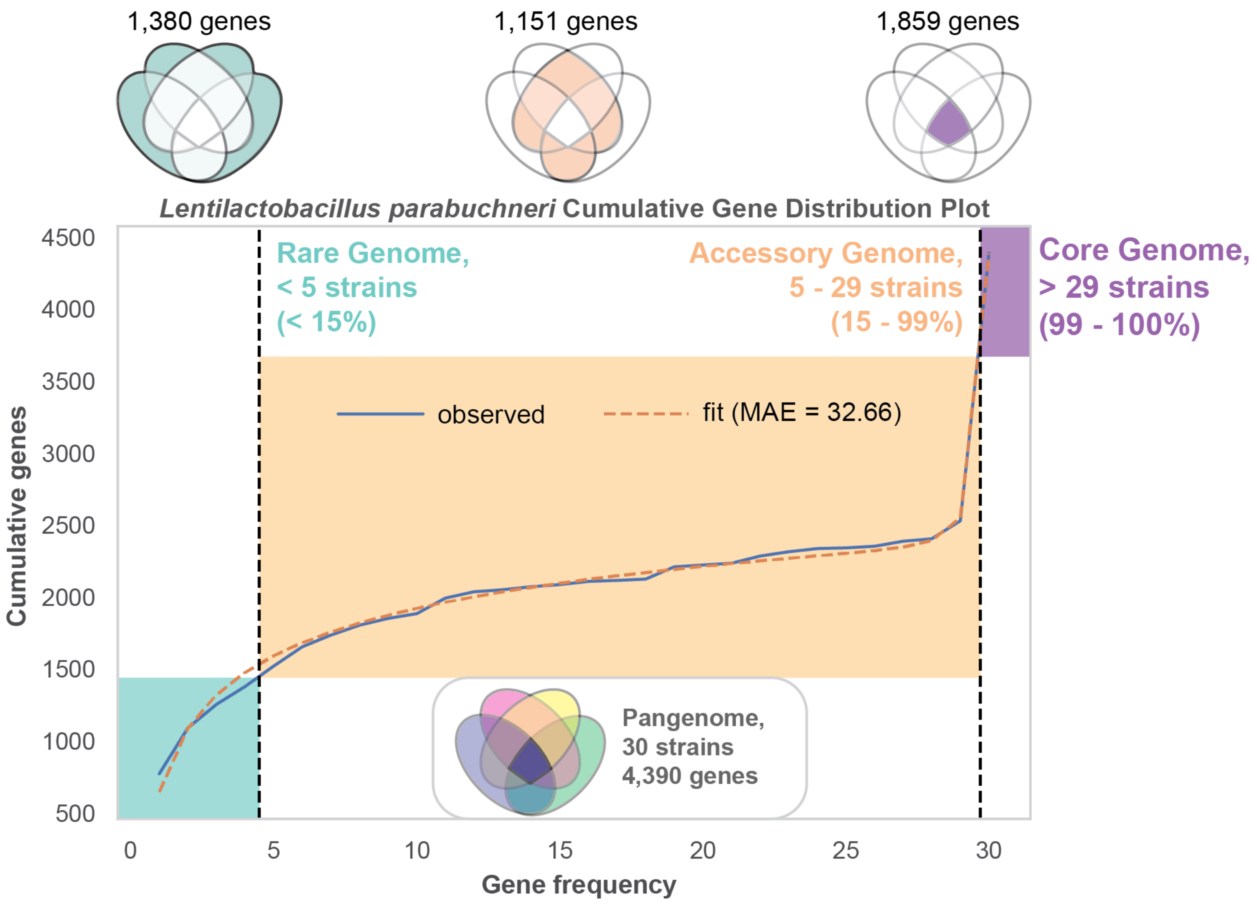


***Supplementary Figure S17: Gene CDF plot for Lentilactobacillus parabuchneri.***

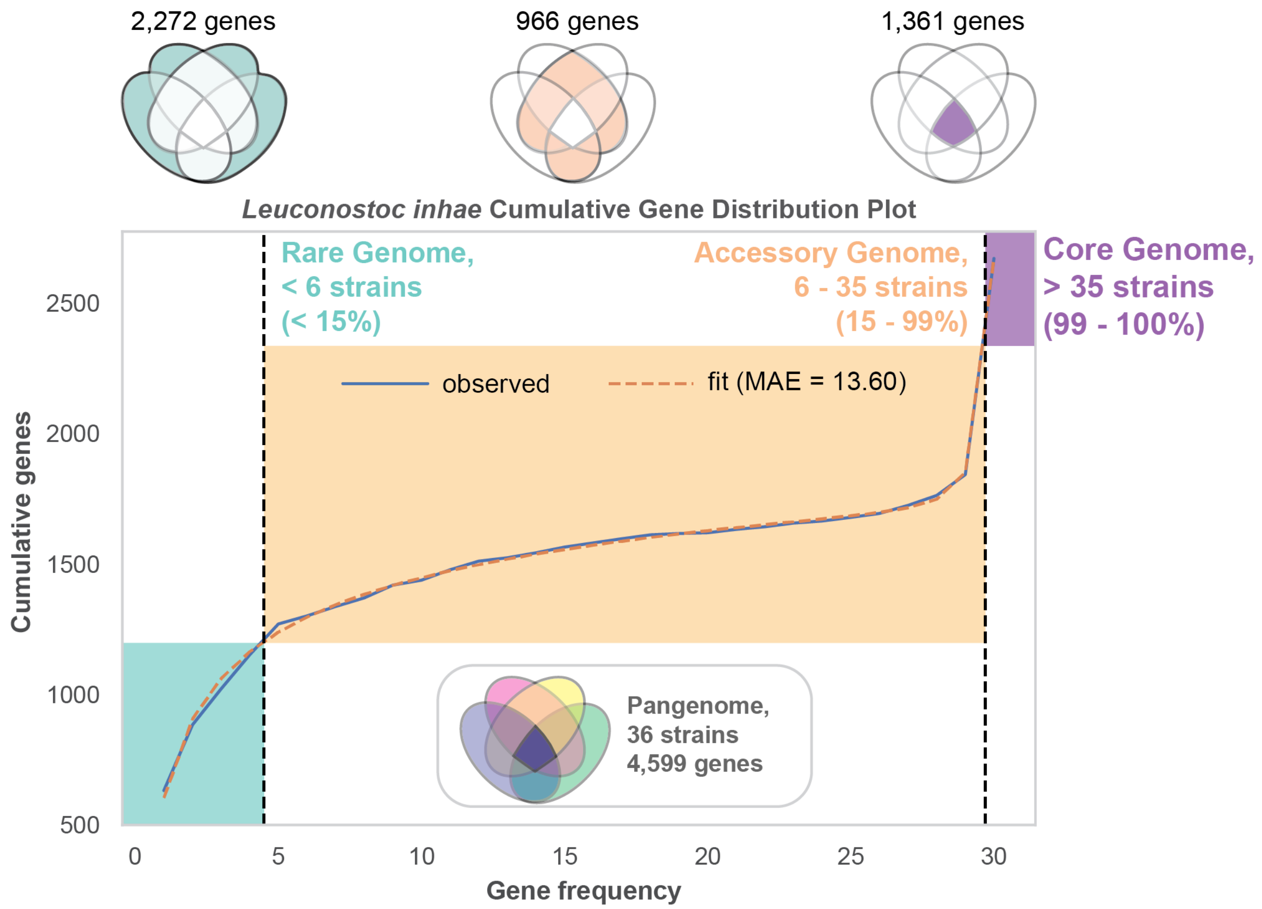


***Supplementary Figure S18: Gene CDF plot for Leuconostoc inhae.***

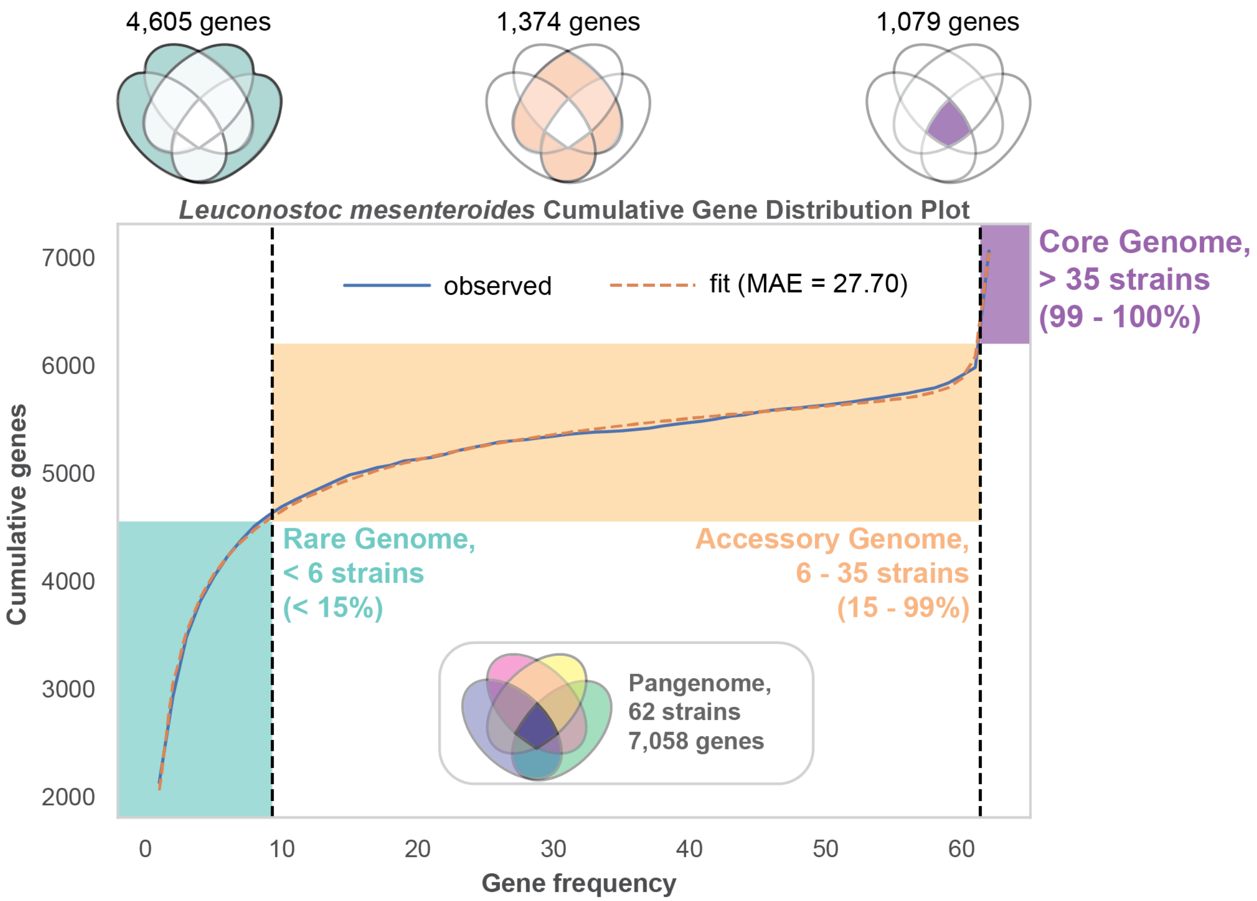


***Supplementary Figure S19: Gene CDF plot for Leuconostoc mesenteroides.***

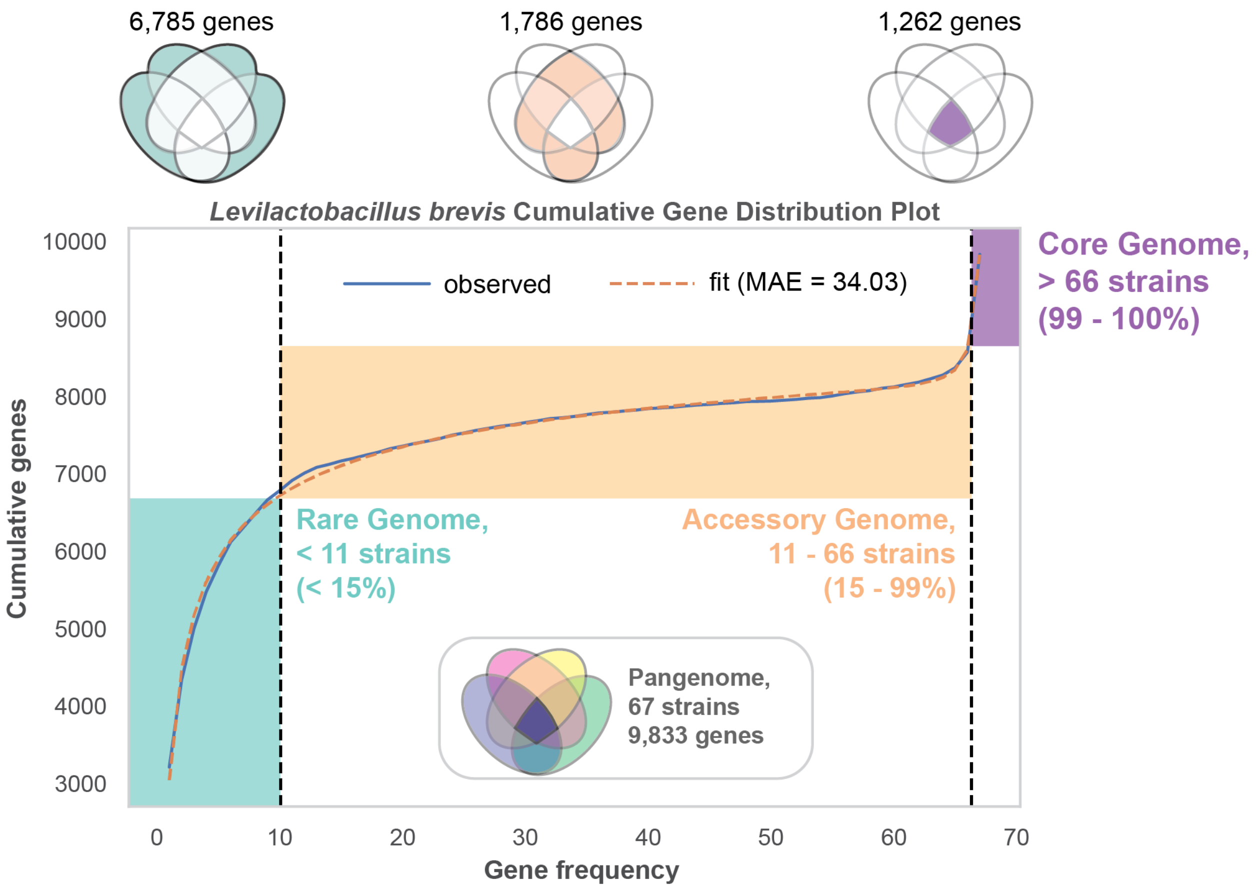


***Supplementary Figure S20: Gene CDF plot for Levilactobacillus brevis.***

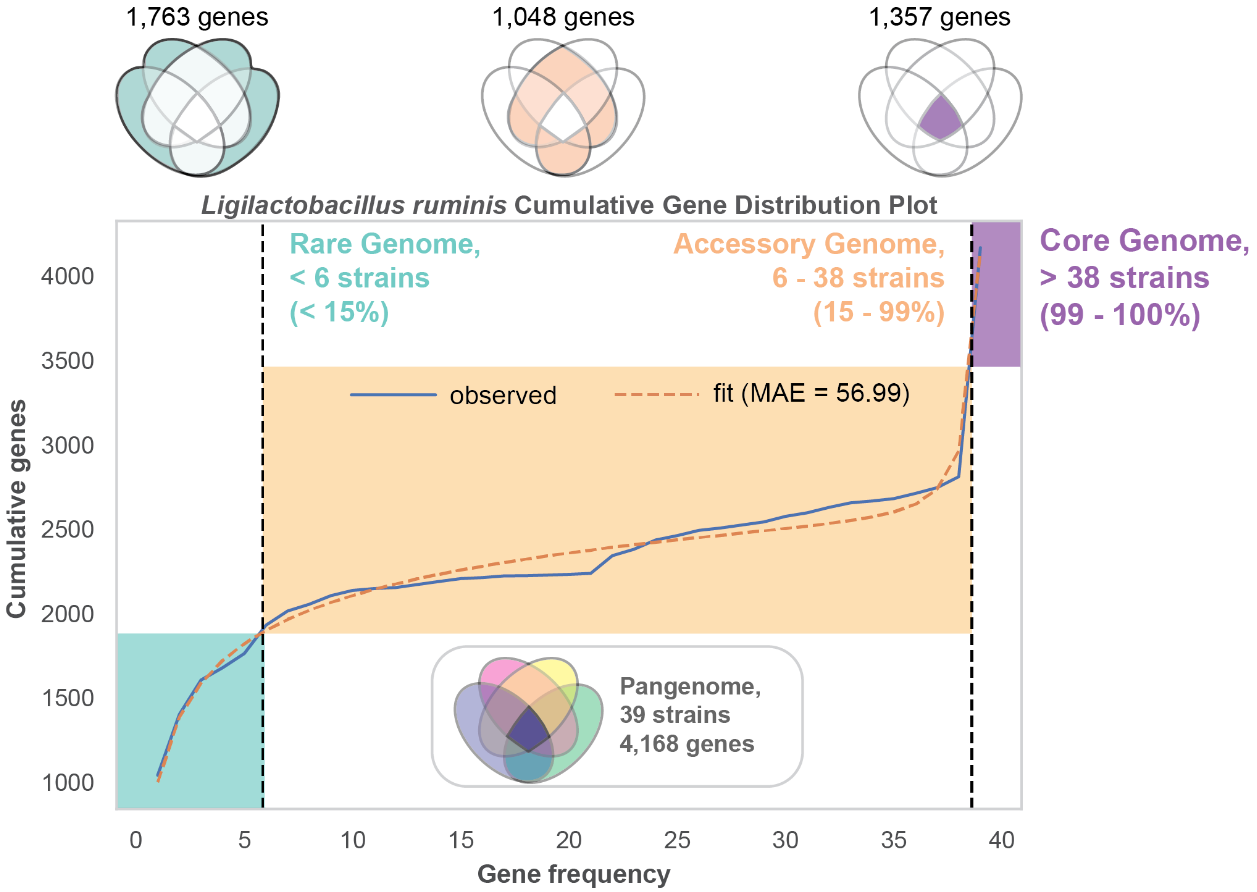


***Supplementary Figure S21: Gene CDF plot for Ligilactobacillus ruminis.***

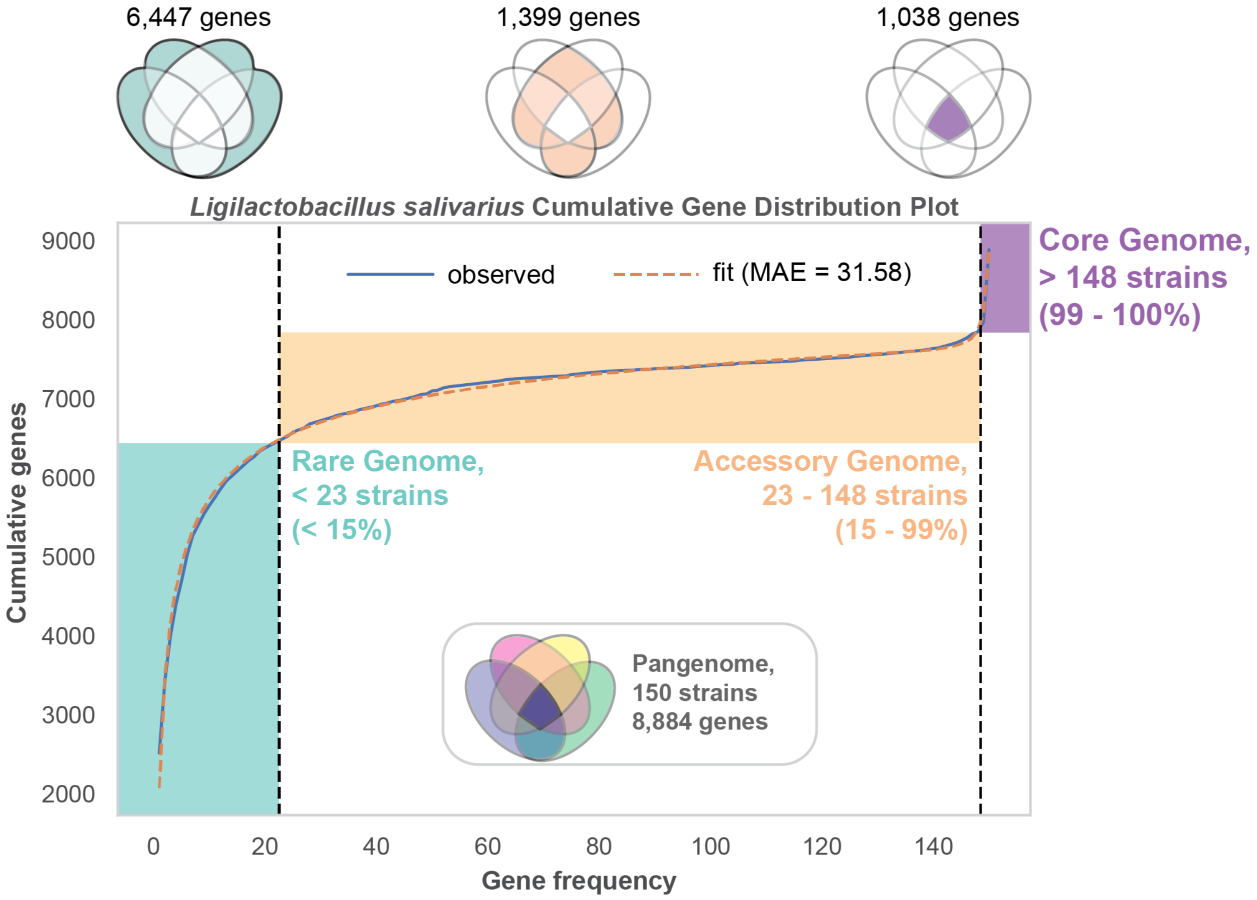


***Supplementary Figure S22: Gene CDF plot for Ligilactobacillus salivarius.***

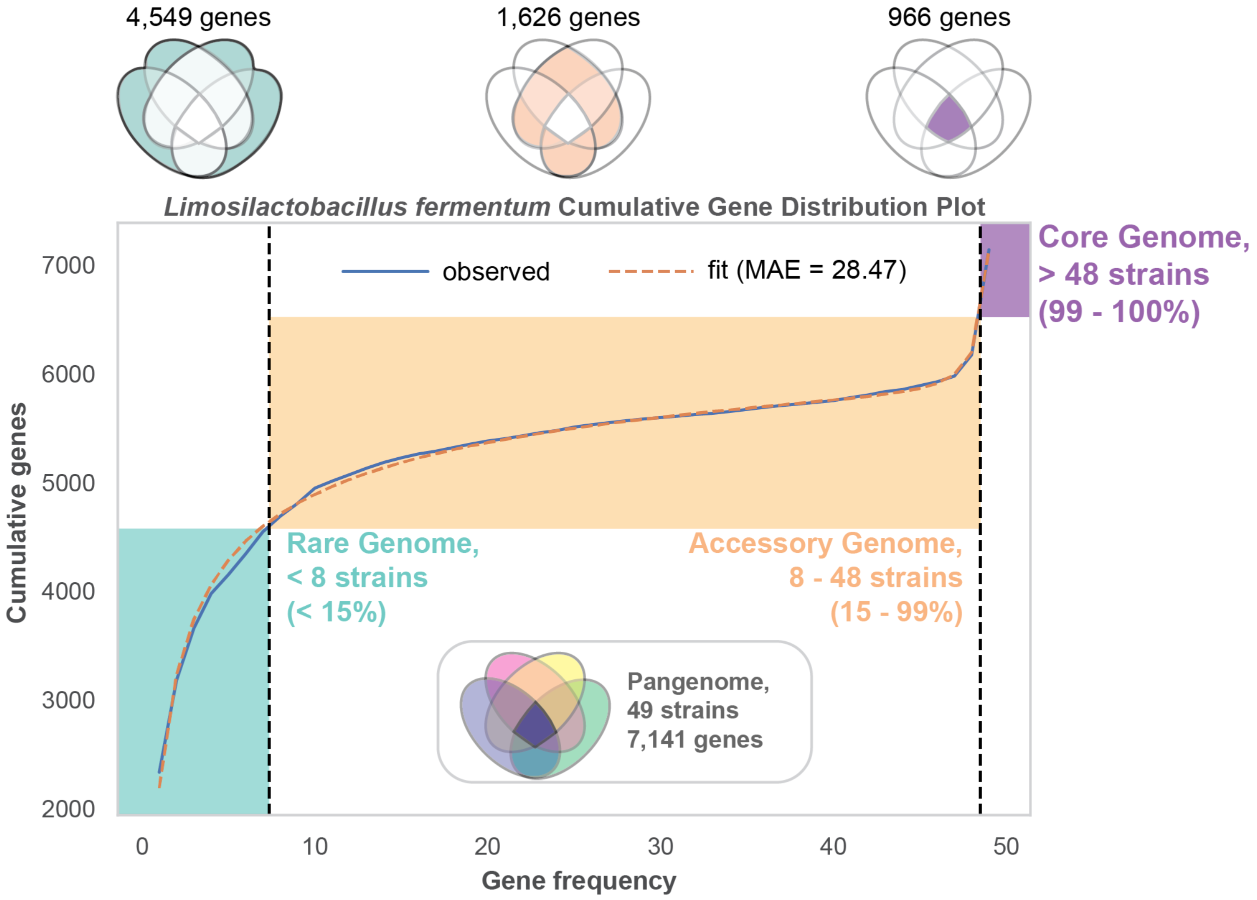


***Supplementary Figure S23: Gene CDF plot for Limosilactobacillus fermentum.***

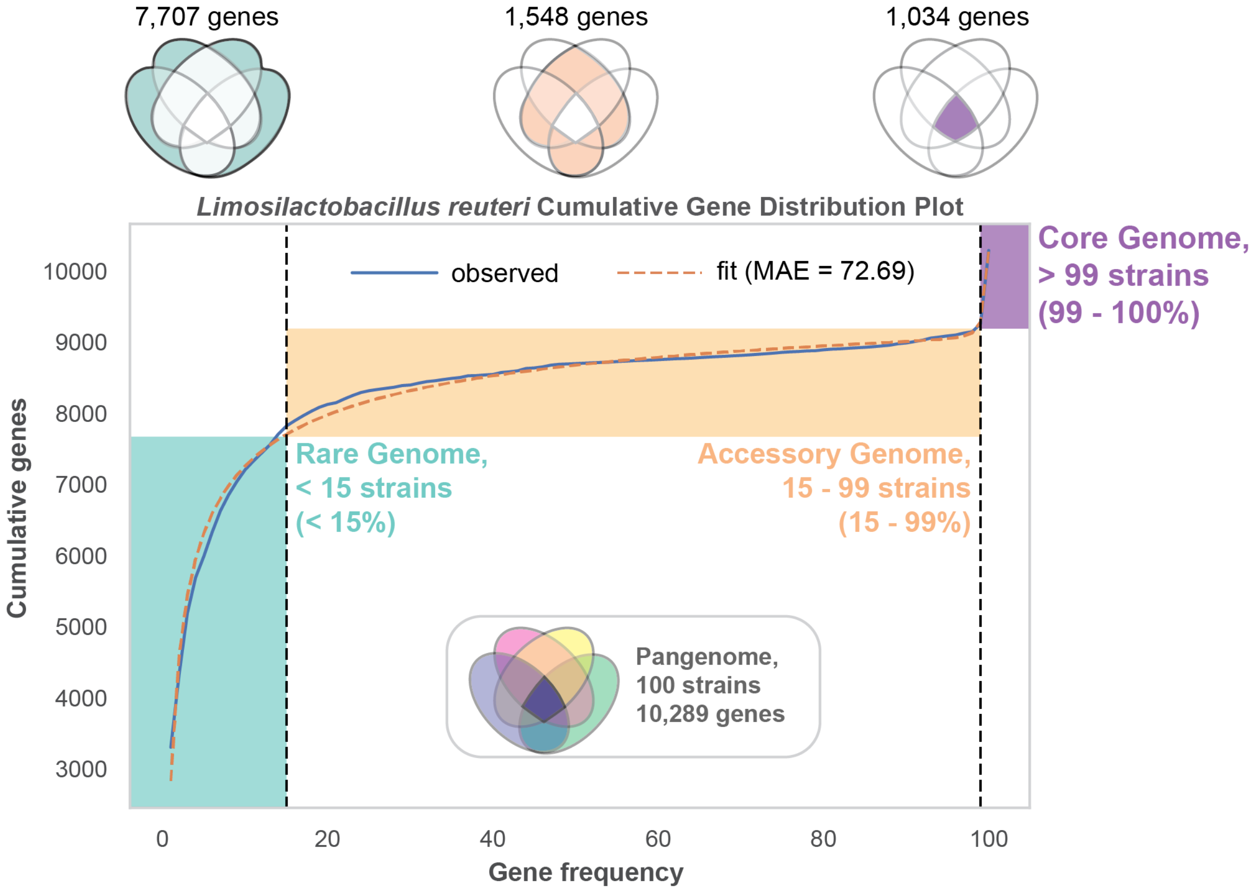


***Supplementary Figure S24: Gene CDF plot for Limosilactobacillus reuteri.***

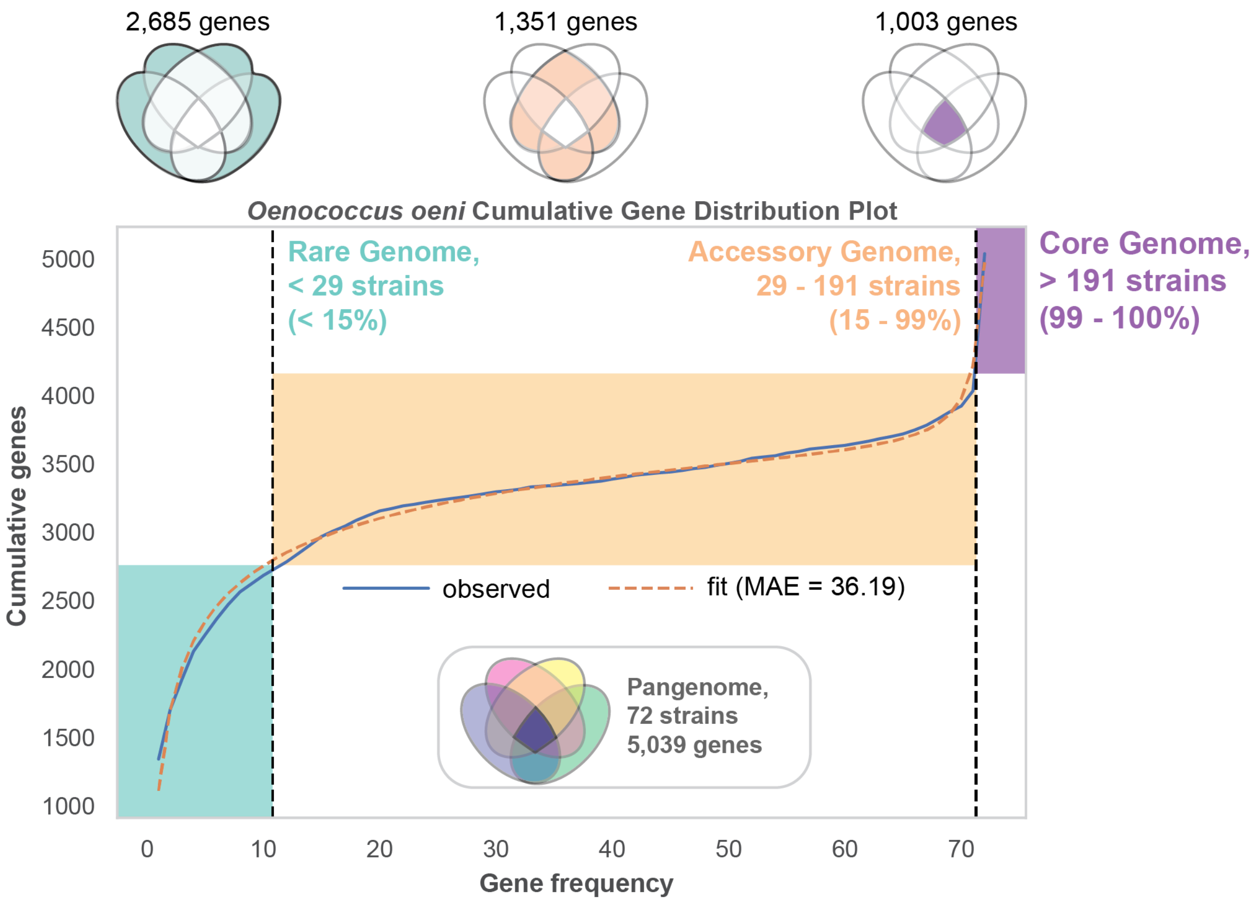


***Supplementary Figure S25: Gene CDF plot for Oenococcus oeni.***

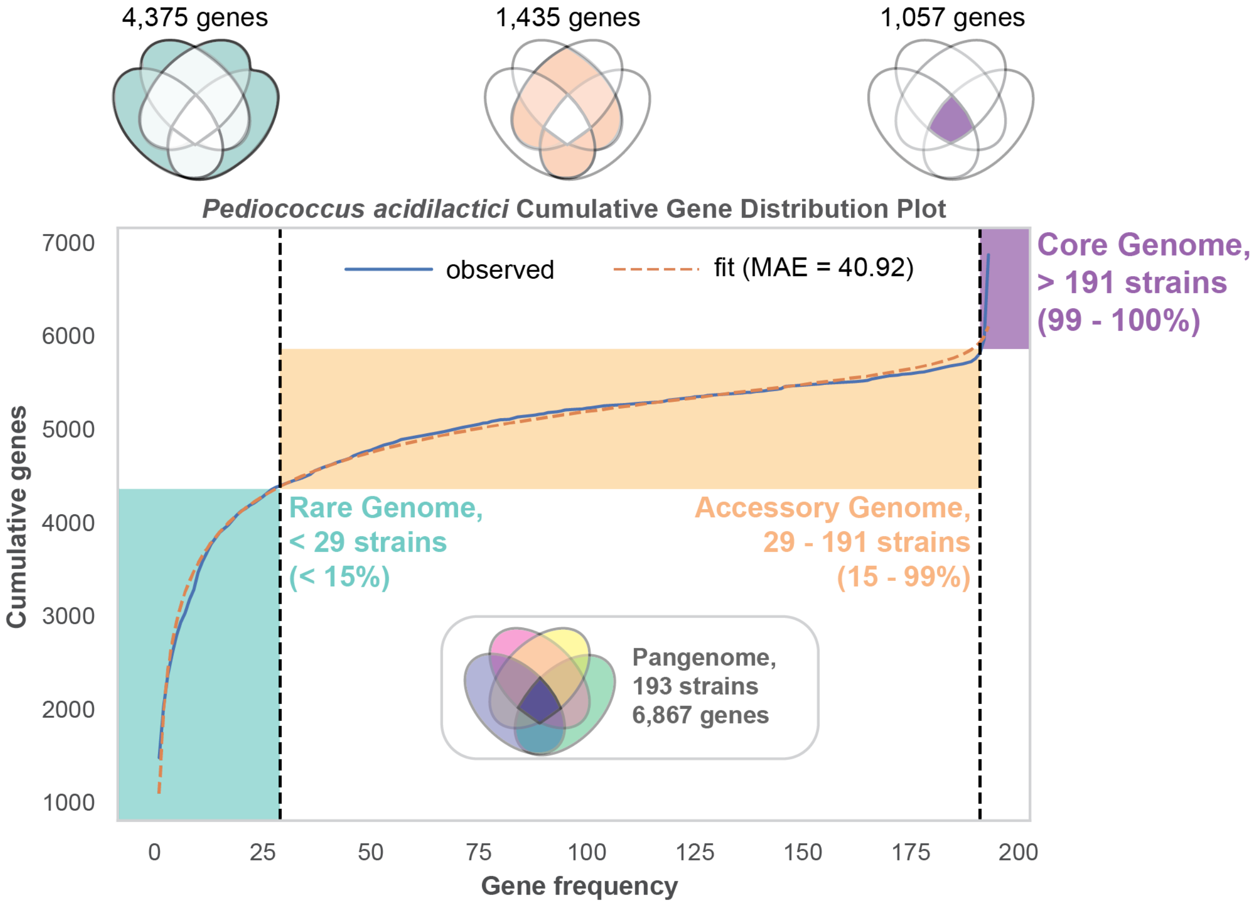


***Supplementary Figure S26: Gene CDF plot for Pediococcus acidilactici.***

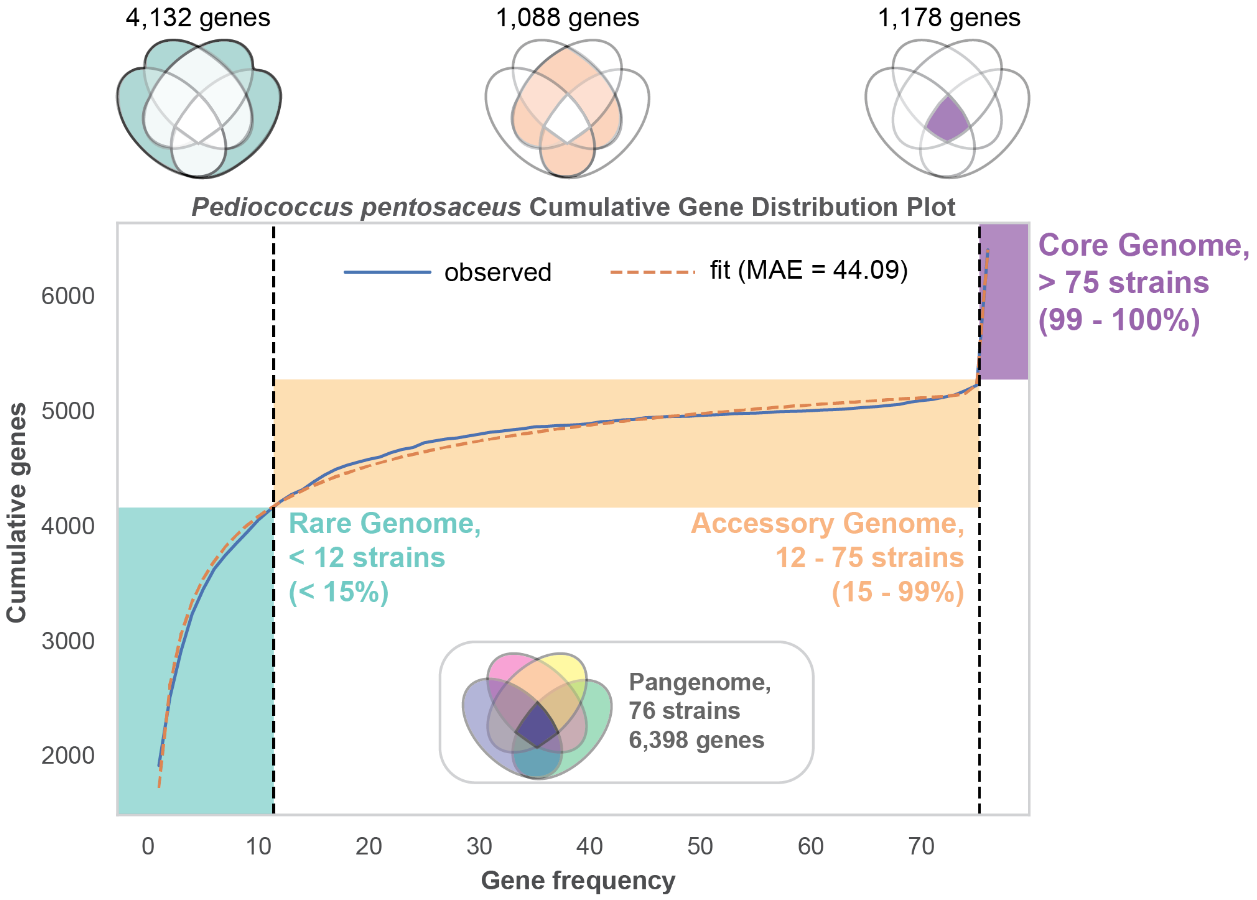


***Supplementary Figure S27: Gene CDF plot for Pediococcus pentosaceus.***

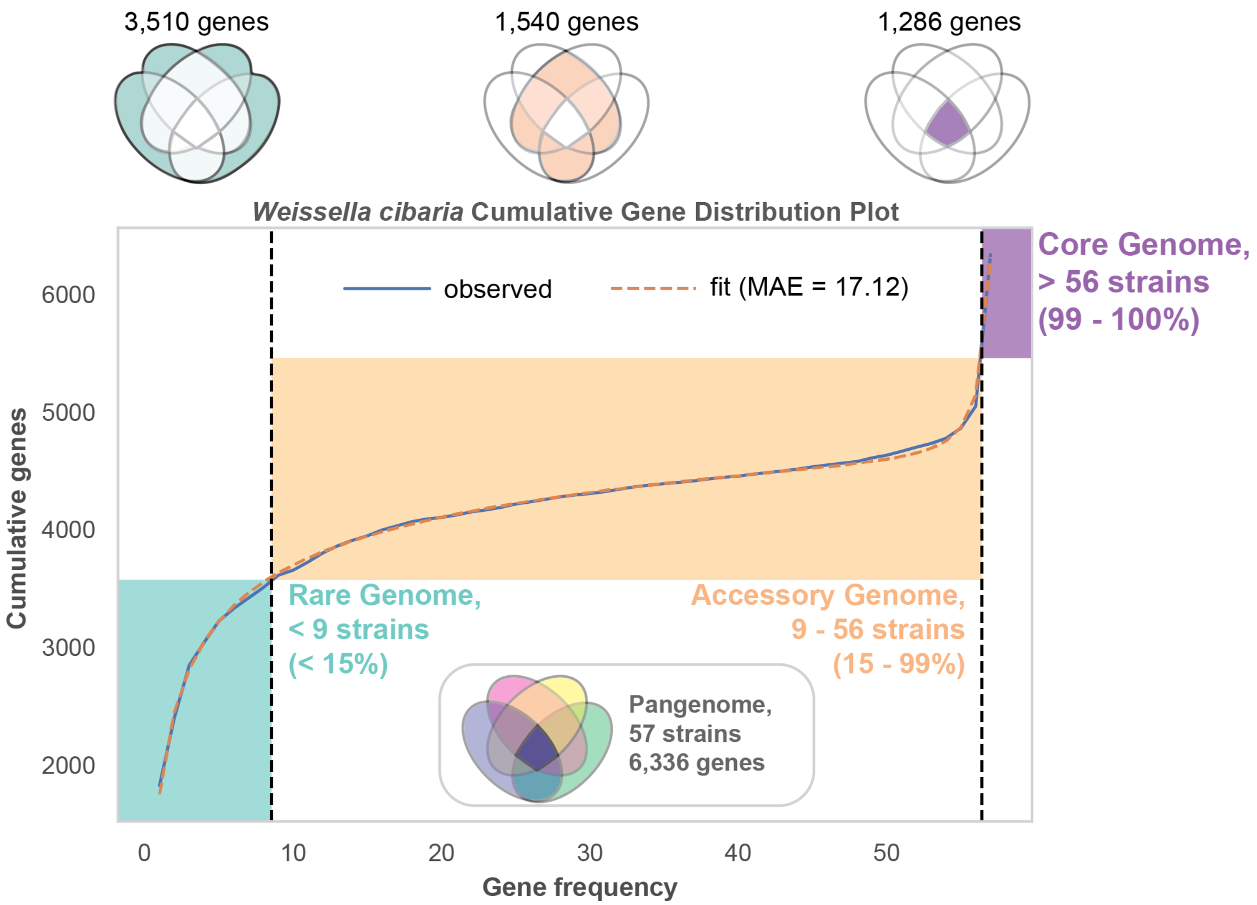


***Supplementary Figure S28: Gene CDF plot for Weissella cibaria.***

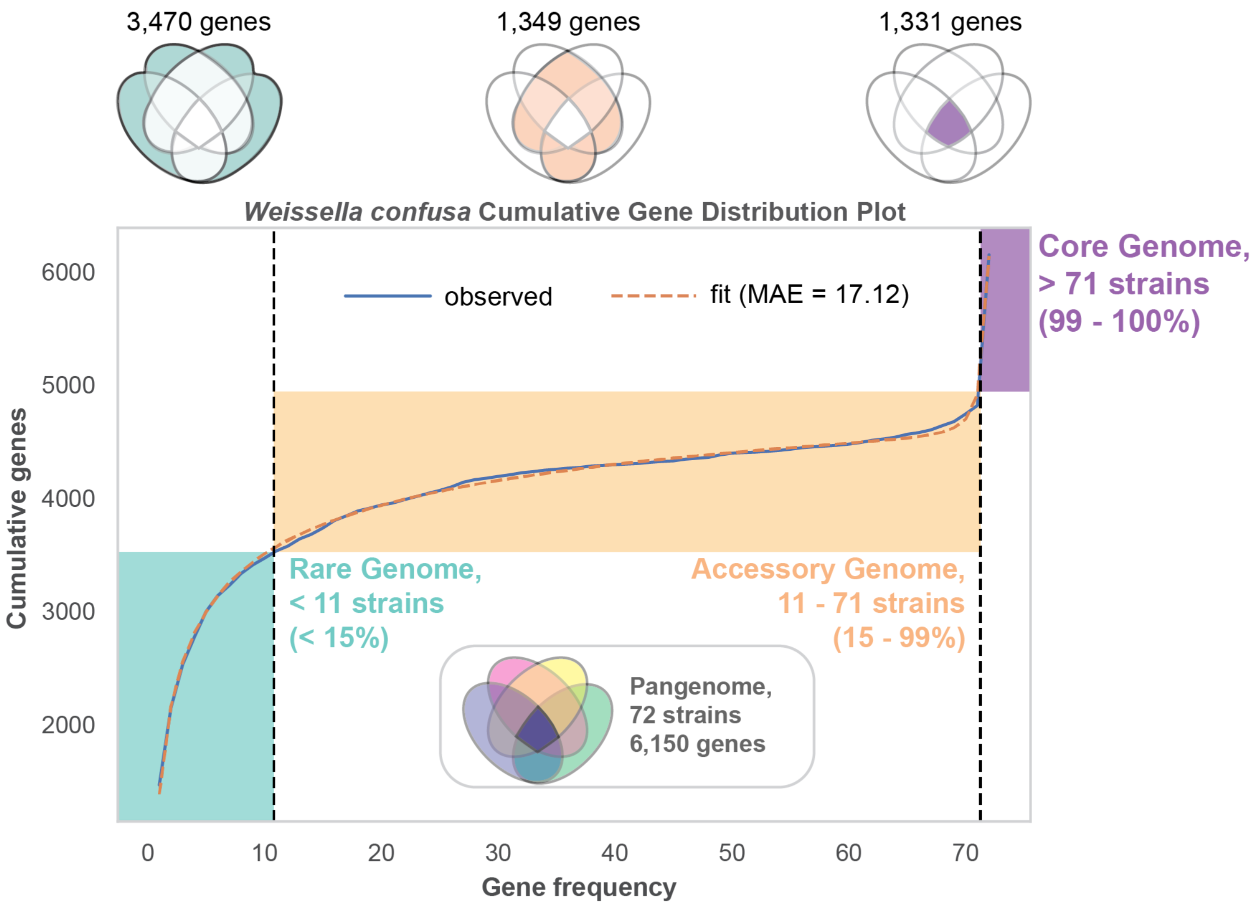


***Supplementary Figure S29: Gene CDF plot for Weissella confusa.***

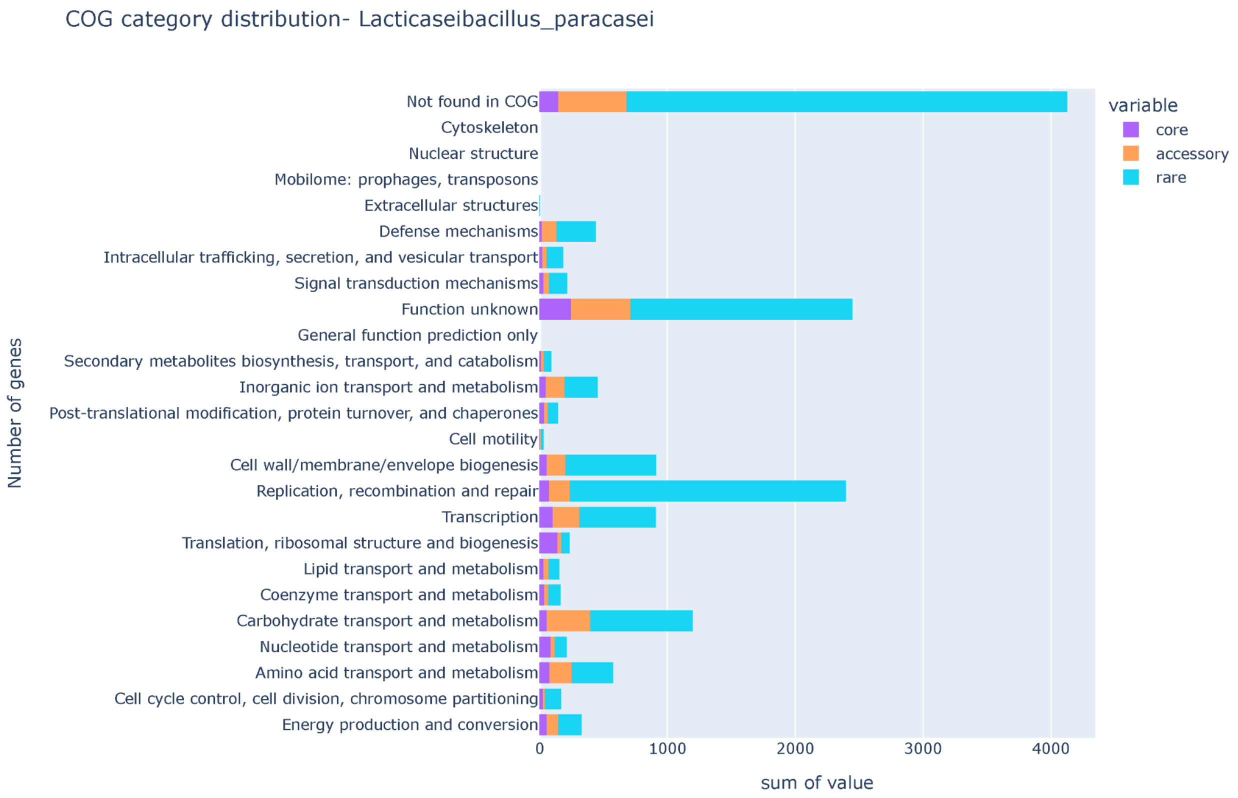


Loading [MathJax]/extensions/MathMenu.js

***Supplementary Figure S30: The barplot depicts the COG functions among core, accessory, and rare pangenome in Lacticaseibacillus paracasei. The barplot is between the number of genes vs COG functions. Note: " - " refers to genes with no COG annotation.***

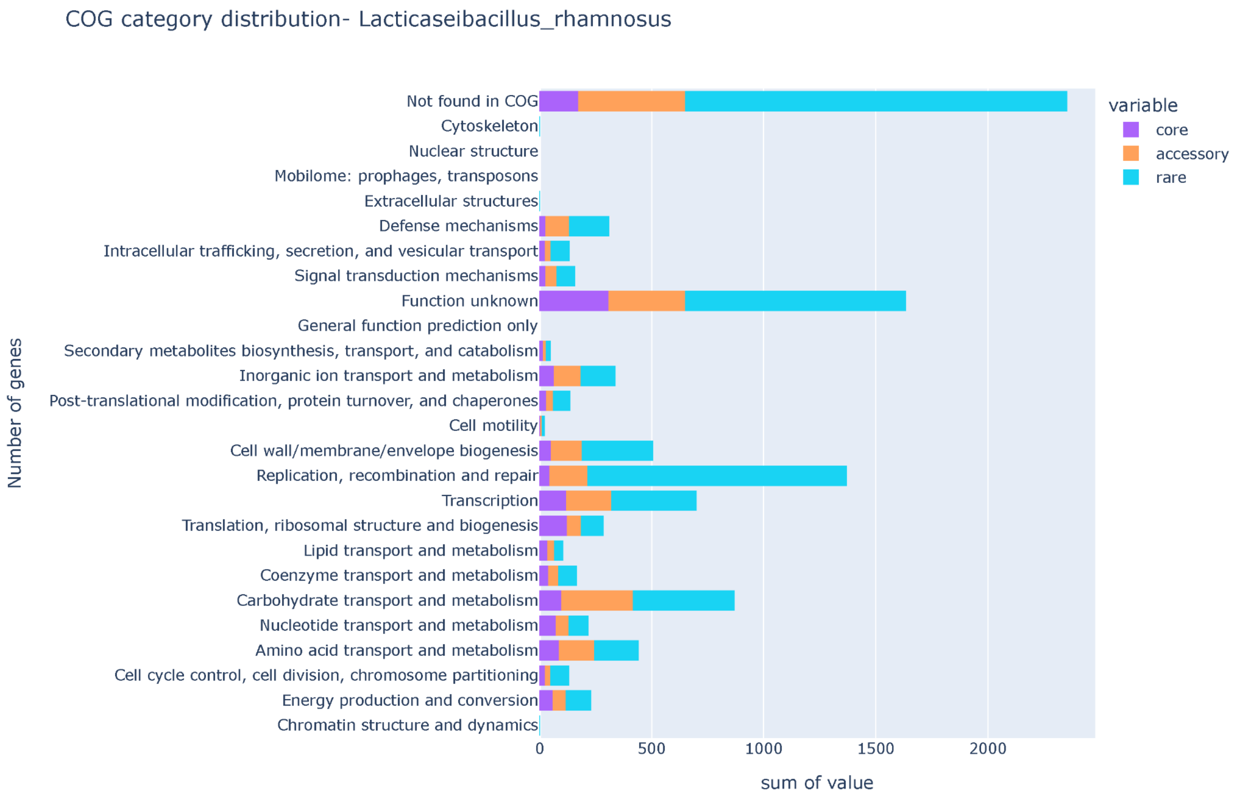


***Supplementary Figure S31: The barplot depicts the COG functions among core, accessory, and rare pangenome in Lacticaseibacillus rhamnosus. The barplot is between the number of genes vs COG functions. Note: " - " refers to genes with no COG annotation.***

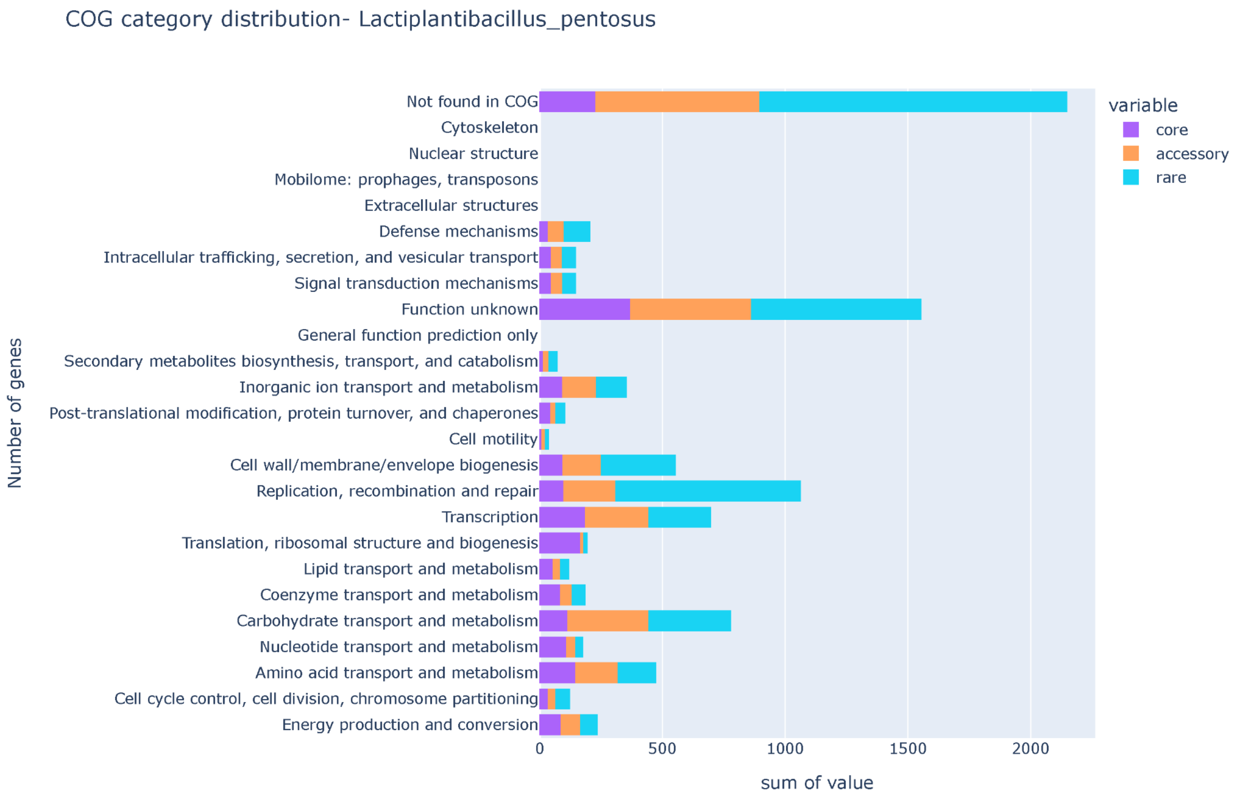


Loading [MathJax]/extensions/MathMenu.js

***Supplementary Figure S32: The barplot depicts the COG functions among core, accessory, and rare pangenome in Lactiplantibacillus pentosus. The barplot is between the number of genes vs COG functions. Note: “ - ” refers to genes with no COG annotation.***

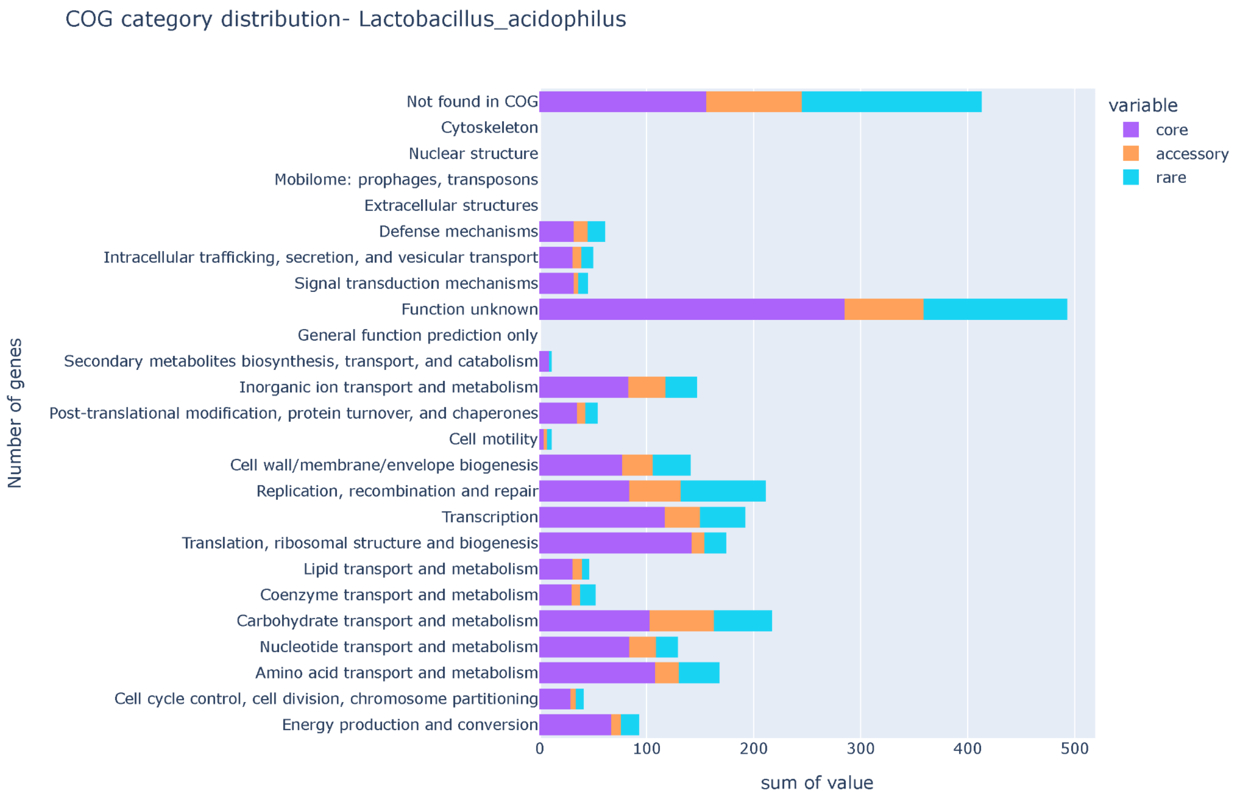


Loading [MathJax]/extensions/MathMenu.js

***Supplementary Figure S33: The barplot depicts the COG functions among core, accessory, and rare pangenome in Lactobacillus acdiophillus. The barplot is between the number of genes vs COG functions. Note: " - " refers to genes with no COG annotation.***

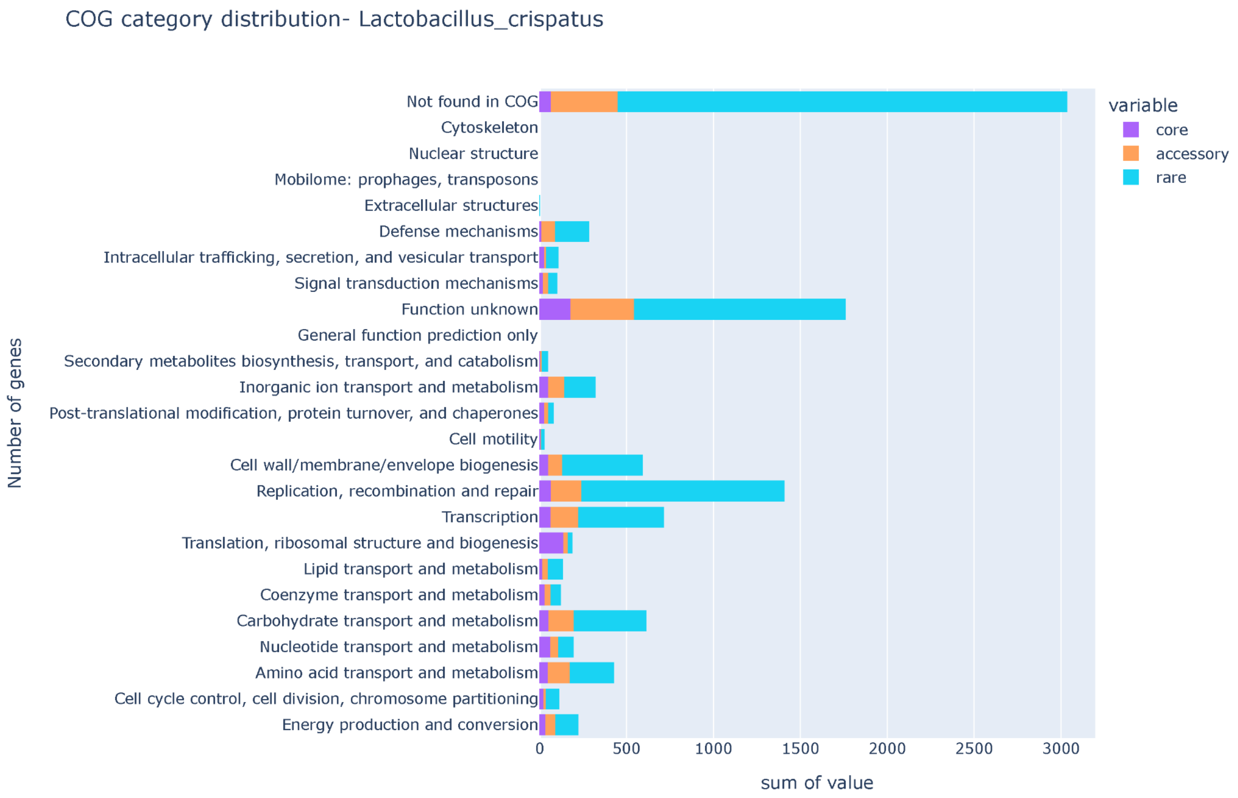


Loading [MathJax]/extensions/MathMenu.js

***Supplementary Figure S34: The barplot depicts the COG functions among core, accessory, and rare pangenome in Lactobacillus crispatus. The barplot is between the number of genes vs COG functions. Note: " - " refers to genes with no COG annotation.***

***Supplementary Figure S35: The barplot depicts the COG functions among core, accessory, and rare pangenome in Lactobacillus delbrueckii. The barplot is between the number of genes vs COG functions. Note: " - " refers to genes with no COG annotation.***

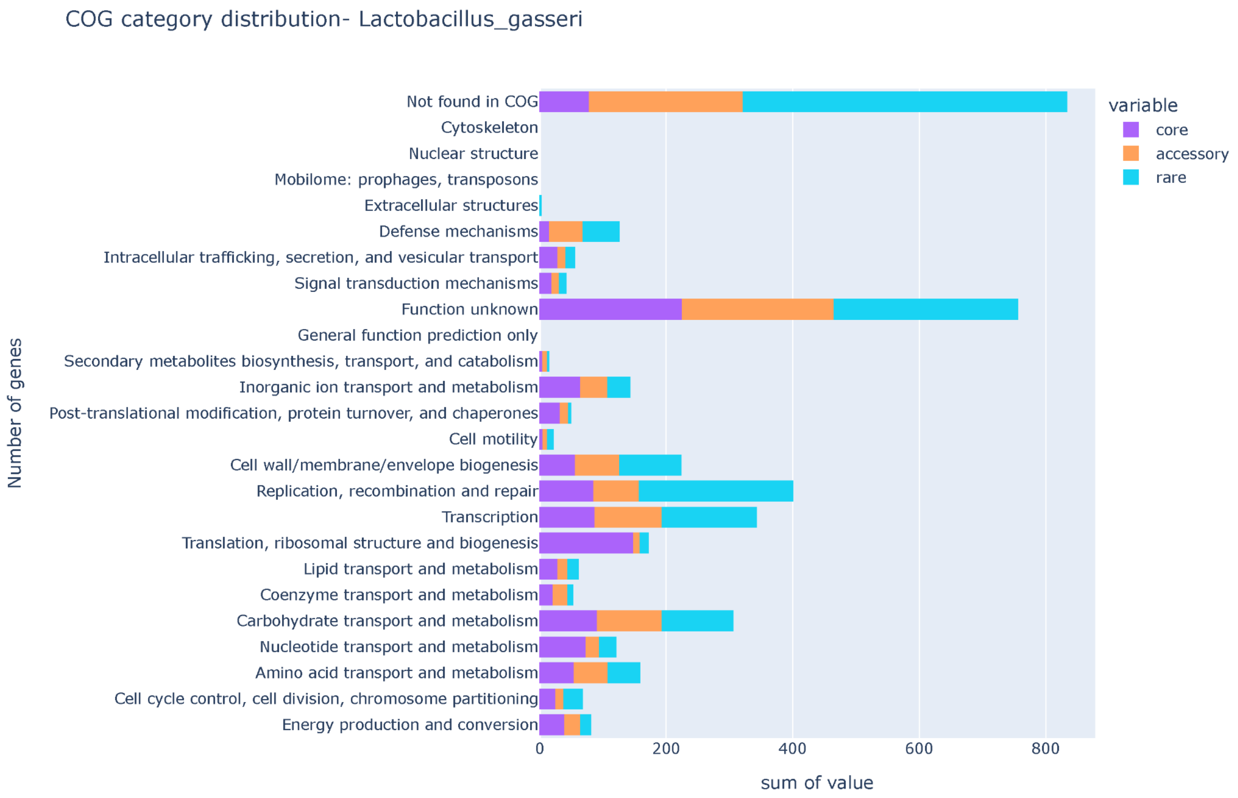


Loading [MathJax]/extensions/MathMenu.js

***Supplementary Figure S36: The barplot depicts the COG functions among core, accessory, and rare pangenome in Lactobacillus gasseri. The barplot is between the number of genes vs COG functions. Note: " - " refers to genes with no COG annotation.***

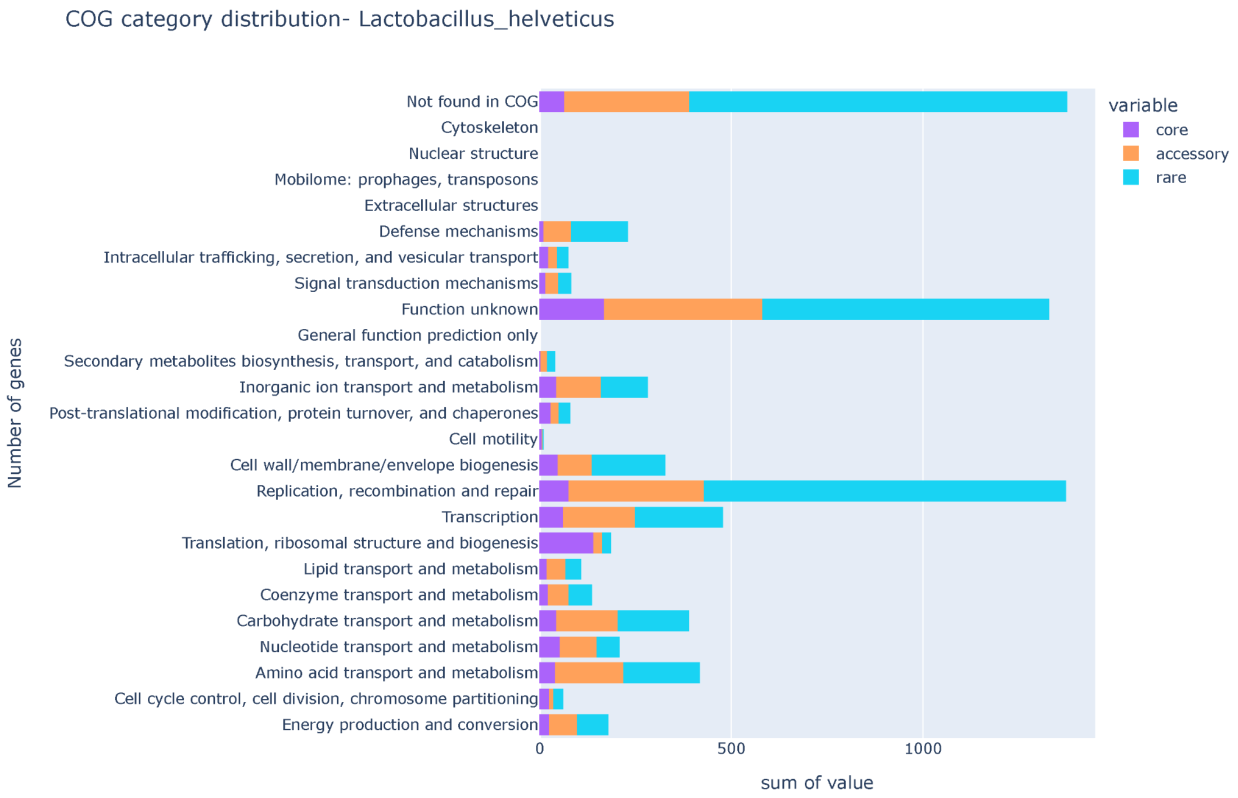


Loading [MathJax]/extensions/MathMenu.js

***Supplementary Figure S37: The barplot depicts the COG functions among core, accessory, and rare pangenome in Lactobacillus helveticus. The barplot is between the number of genes vs COG functions. Note: " - " refers to genes with no COG annotation.***

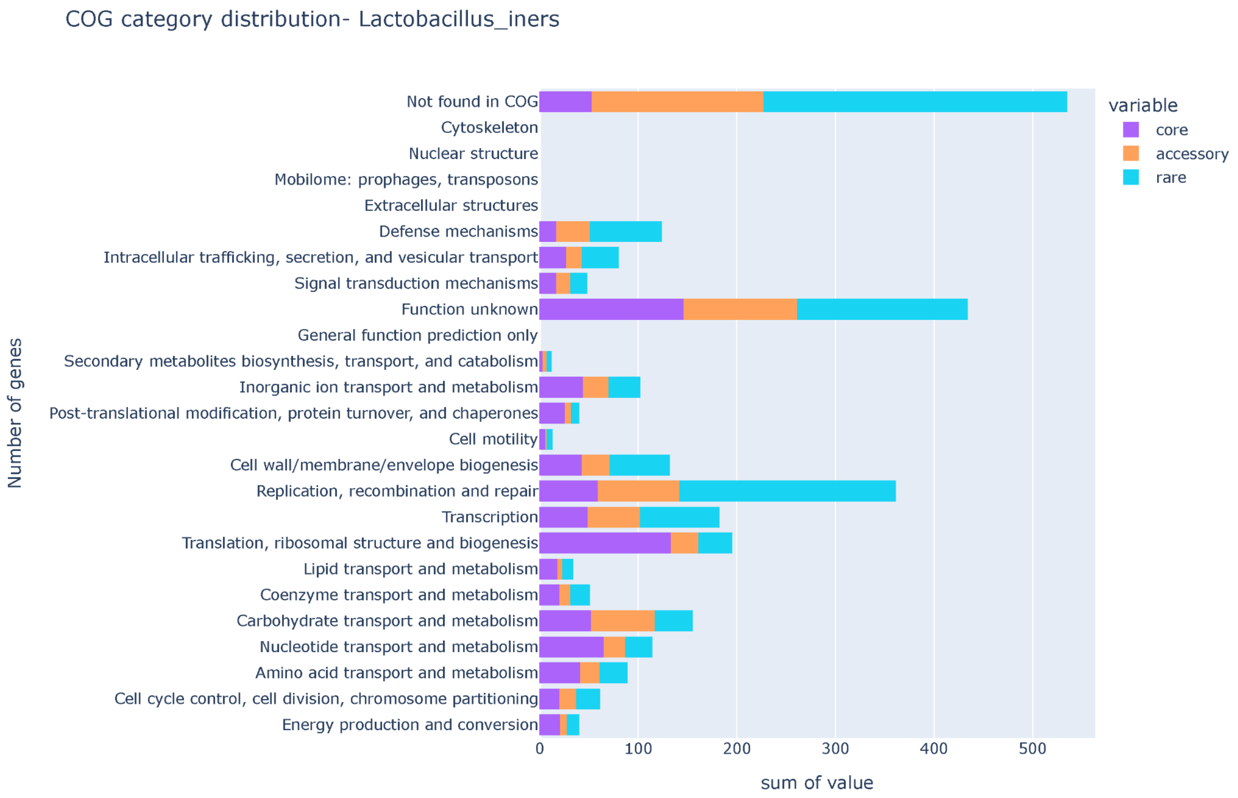


Loading [MathJax]/extensions/MathMenu.js

***Supplementary Figure S38: The barplot depicts the COG functions among core, accessory, and rare pangenome in Lactobacillus iners. The barplot is between the number of genes vs COG functions. Note: " - " refers to genes with no COG annotation.***

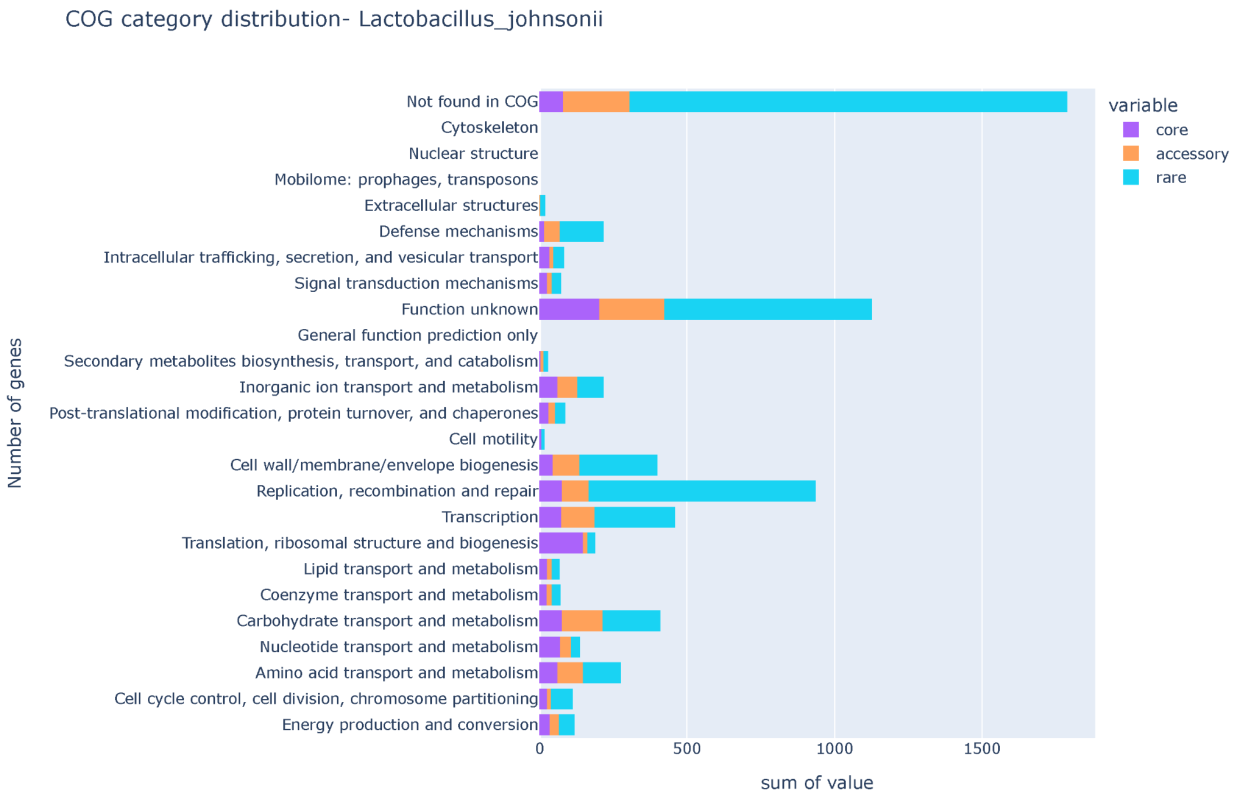


Loading [MathJax]/extensions/MathMenu.js

***Supplementary Figure S39: The barplot depicts the COG functions among core, accessory, and rare pangenome in Lactobacillus johnsonii. The barplot is between the number of genes vs COG functions. Note: " - " refers to genes with no COG annotation.***

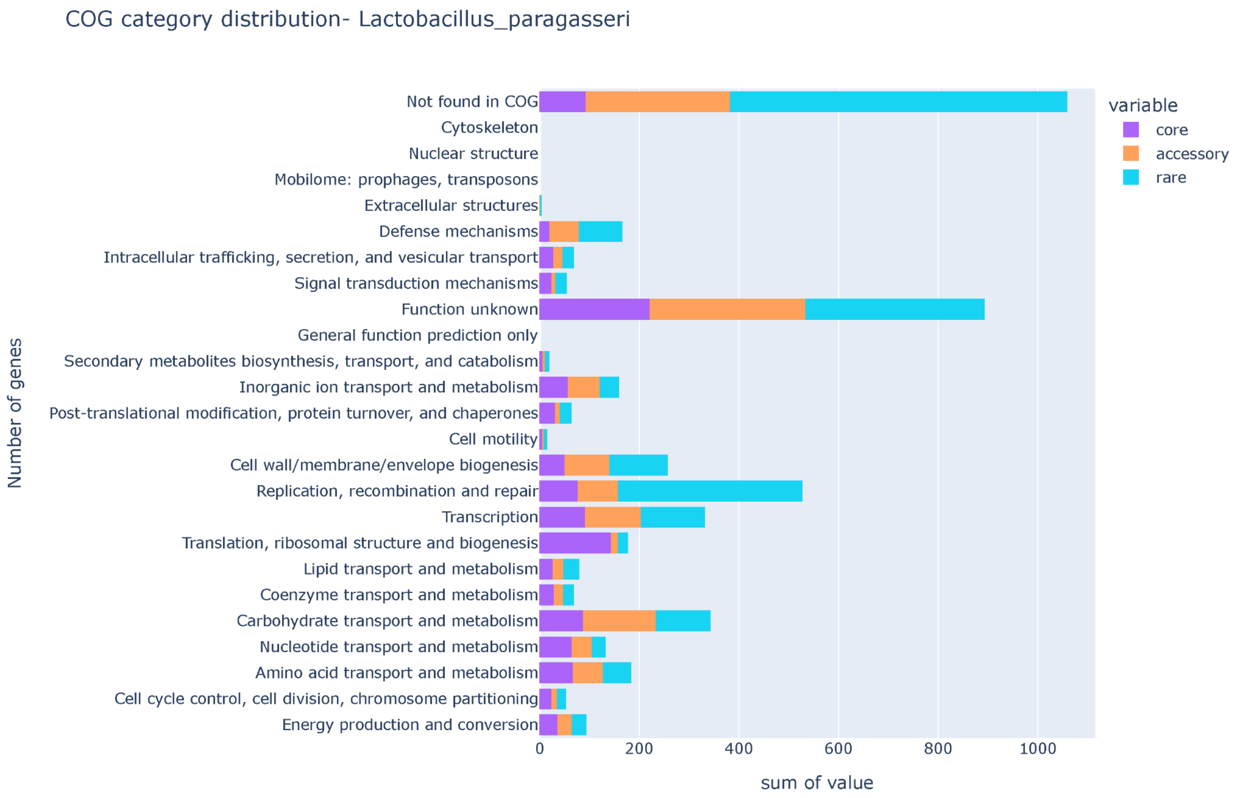


Loading [MathJax]/extensions/MathMenu.js

***Supplementary Figure S40: The barplot depicts the COG functions among core, accessory, and rare pangenome in Lactobacillus paragasseri. The barplot is between the number of genes vs COG functions. Note: " - " refers to genes with no COG annotation.***

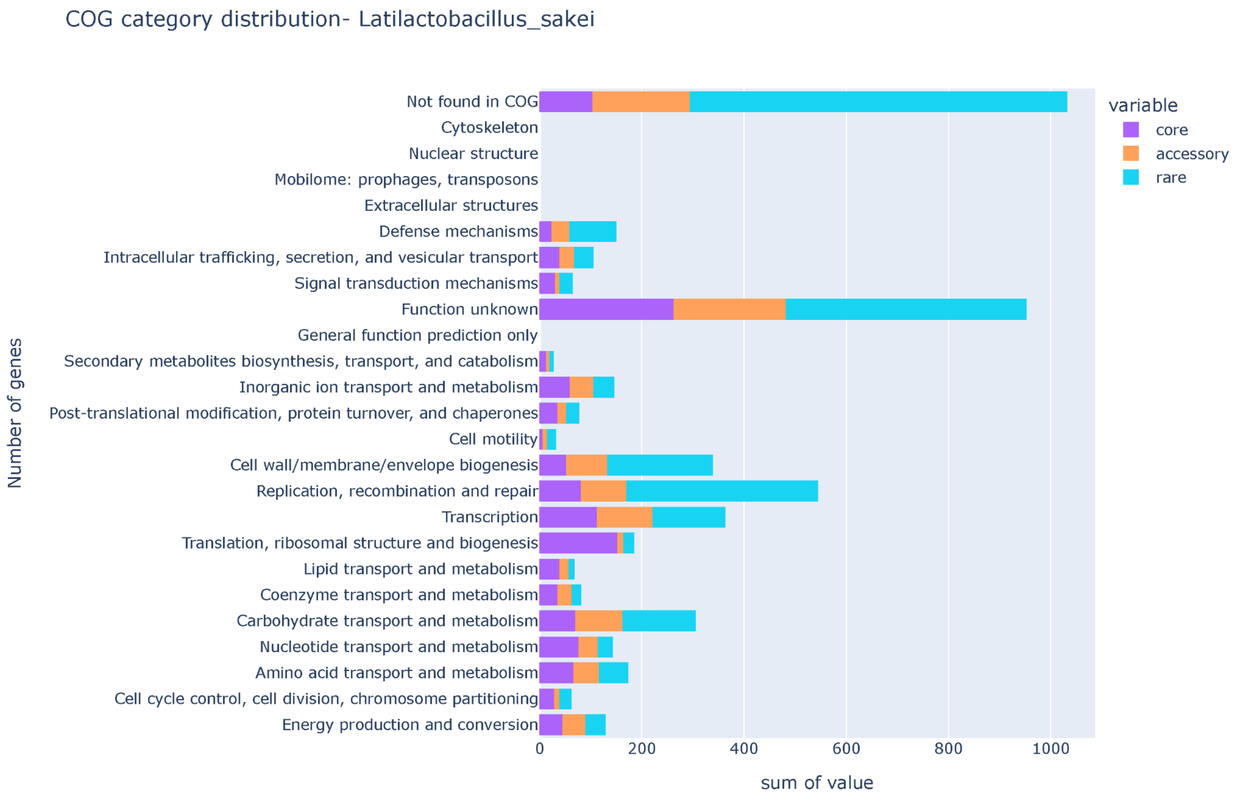


Loading [MathJax]/extensions/MathMenu.js

***Supplementary Figure S41: The barplot depicts the COG functions among core, accessory, and rare pangenome in Latilactobacillus sakei. The barplot is between the number of genes vs COG functions. Note: " - " refers to genes with no COG annotation.***

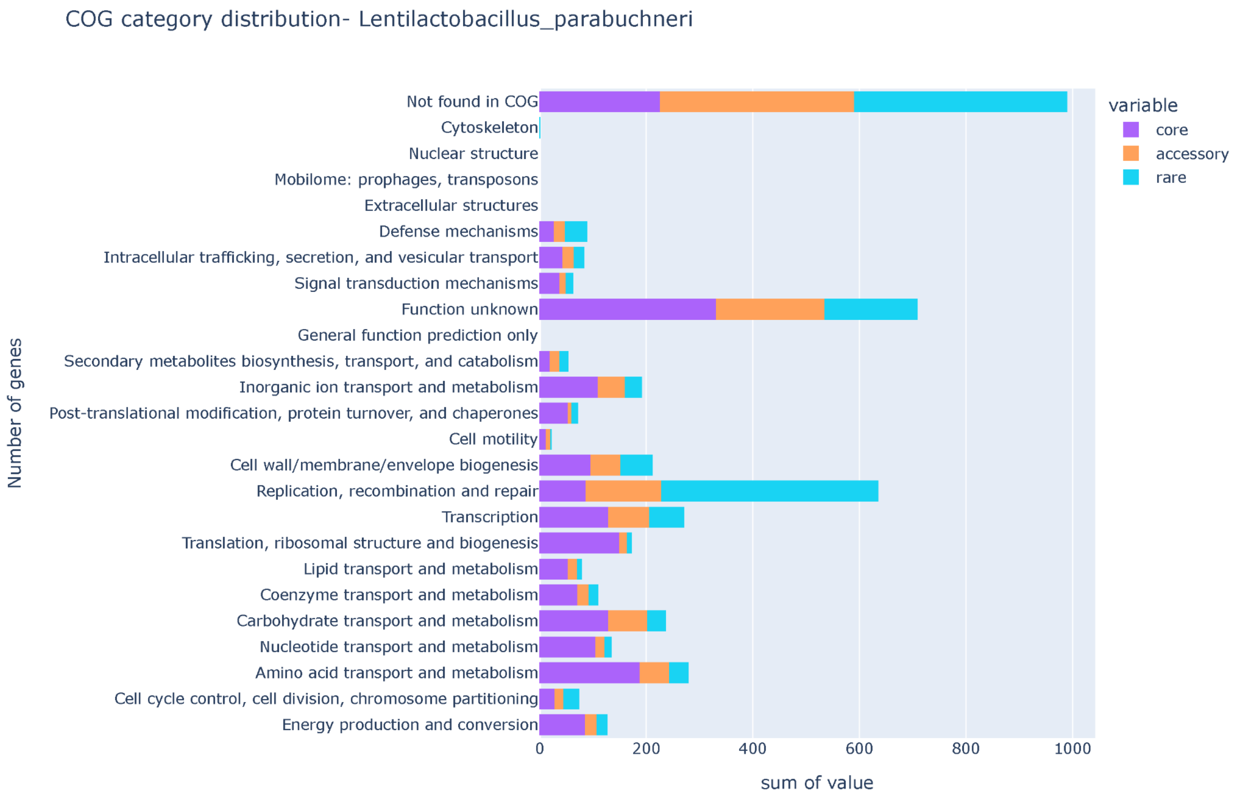


Loading [MathJax]/extensions/MathMenu.js

***Supplementary Figure S42: The barplot depicts the COG functions among core, accessory, and rare pangenome in Lentilactobacillus parabuchneri. The barplot is between the number of genes vs COG functions. Note: " - " refers to genes with no COG annotation.***

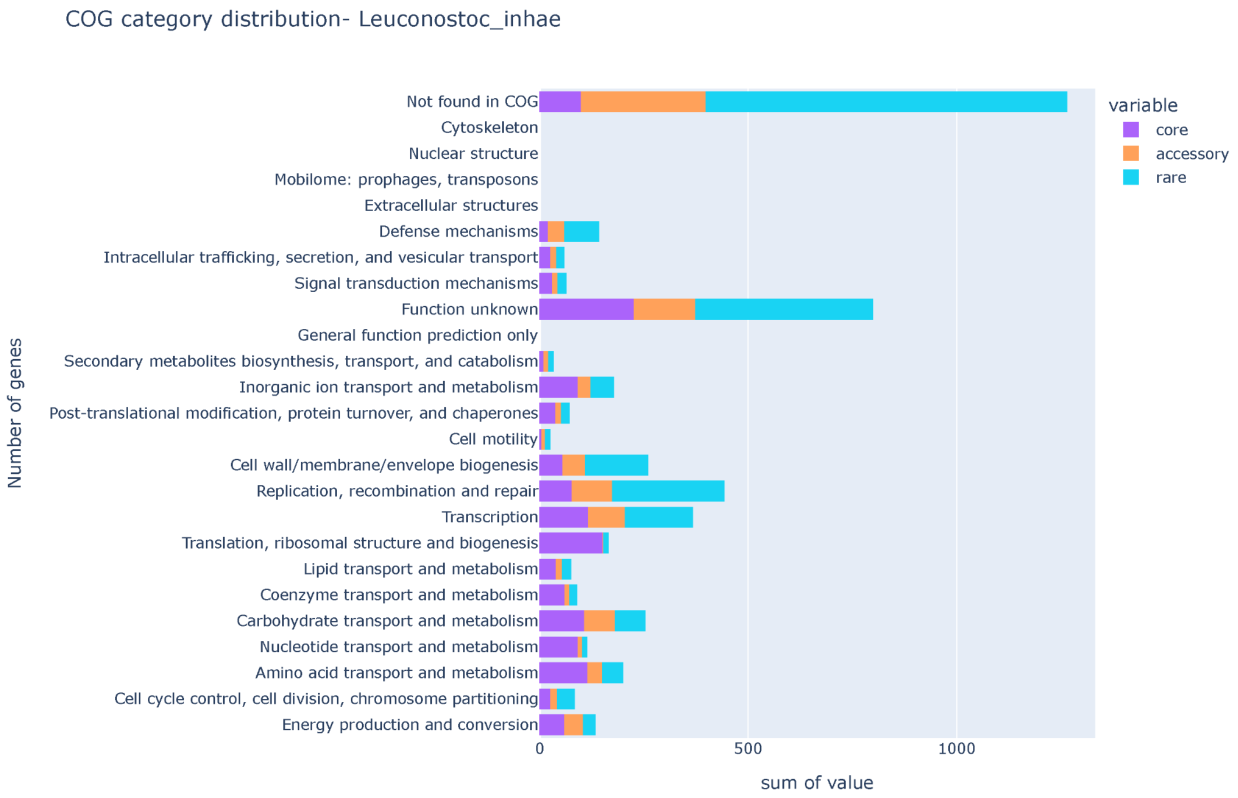


Loading [MathJax]/extensions/MathMenu.js

***Supplementary Figure S43: The barplot depicts the COG functions among core, accessory, and rare pangenome in Leuconostoc inhae. The barplot is between the number of genes vs COG functions. Note: “ - ” refers to genes with no COG annotation.***

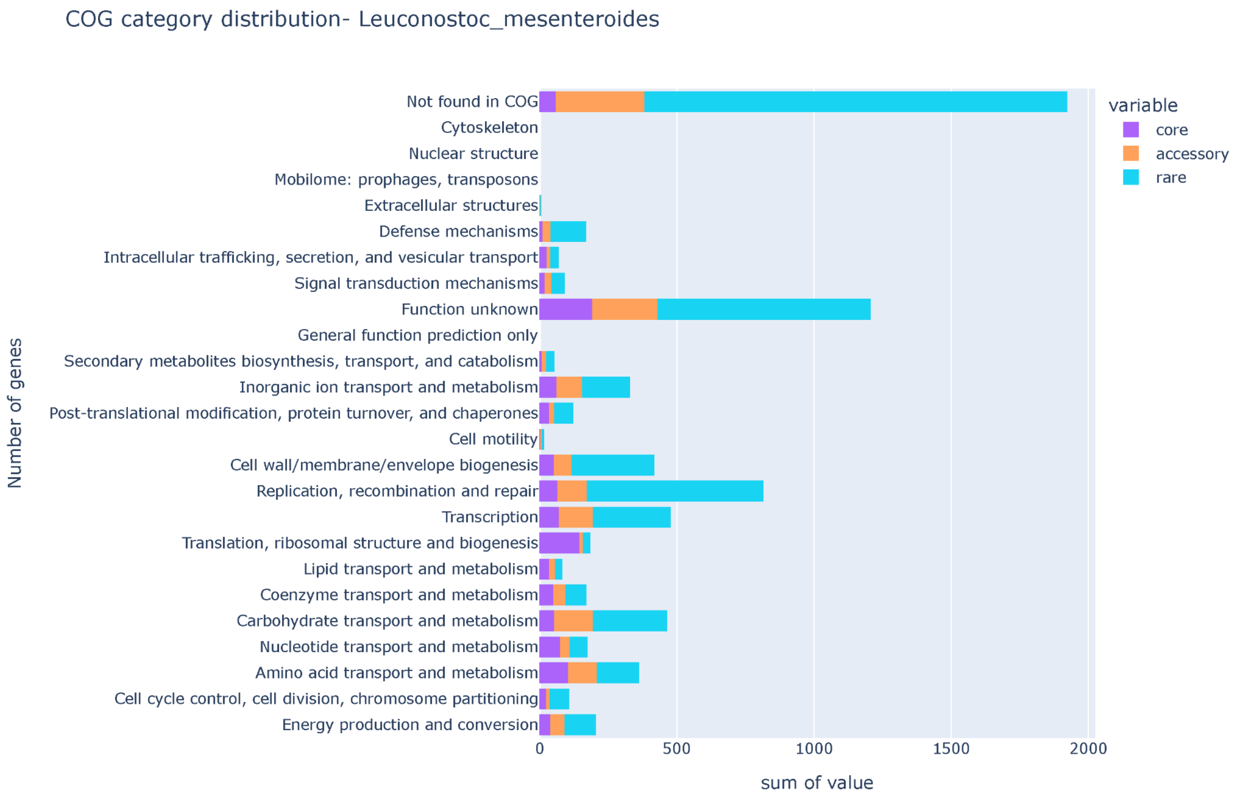


Loading [MathJax]/extensions/MathMenu.js

***Supplementary Figure S44: The barplot depicts the COG functions among core, accessory, and rare pangenome in Leuconostoc mesenteroides. The barplot is between the number of genes vs COG functions. Note: " - " refers to genes with no COG annotation.***

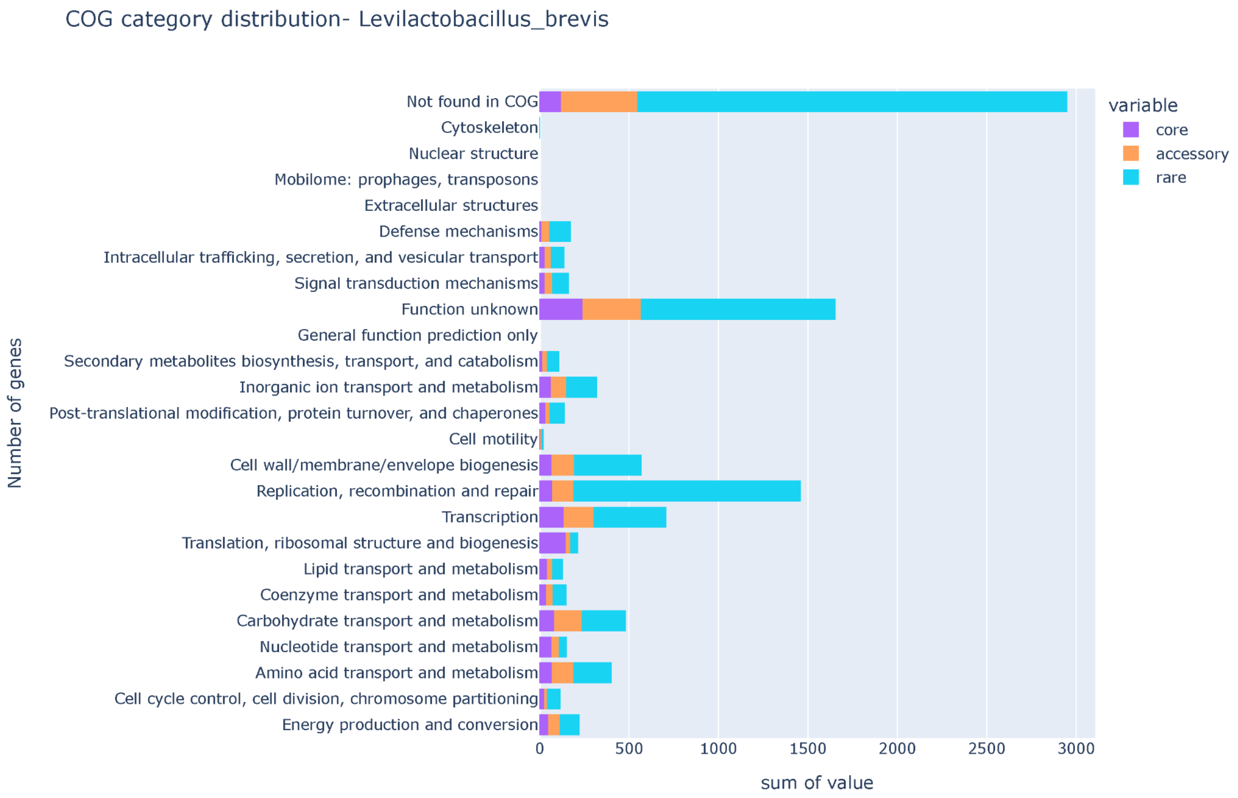


Loading [MathJax]/extensions/MathMenu.js

***Supplementary Figure S45: The barplot depicts the COG functions among core, accessory, and rare pangenome in Levilactobacillus brevis. The barplot is between the number of genes vs COG functions. Note: " - " refers to genes with no COG annotation.***

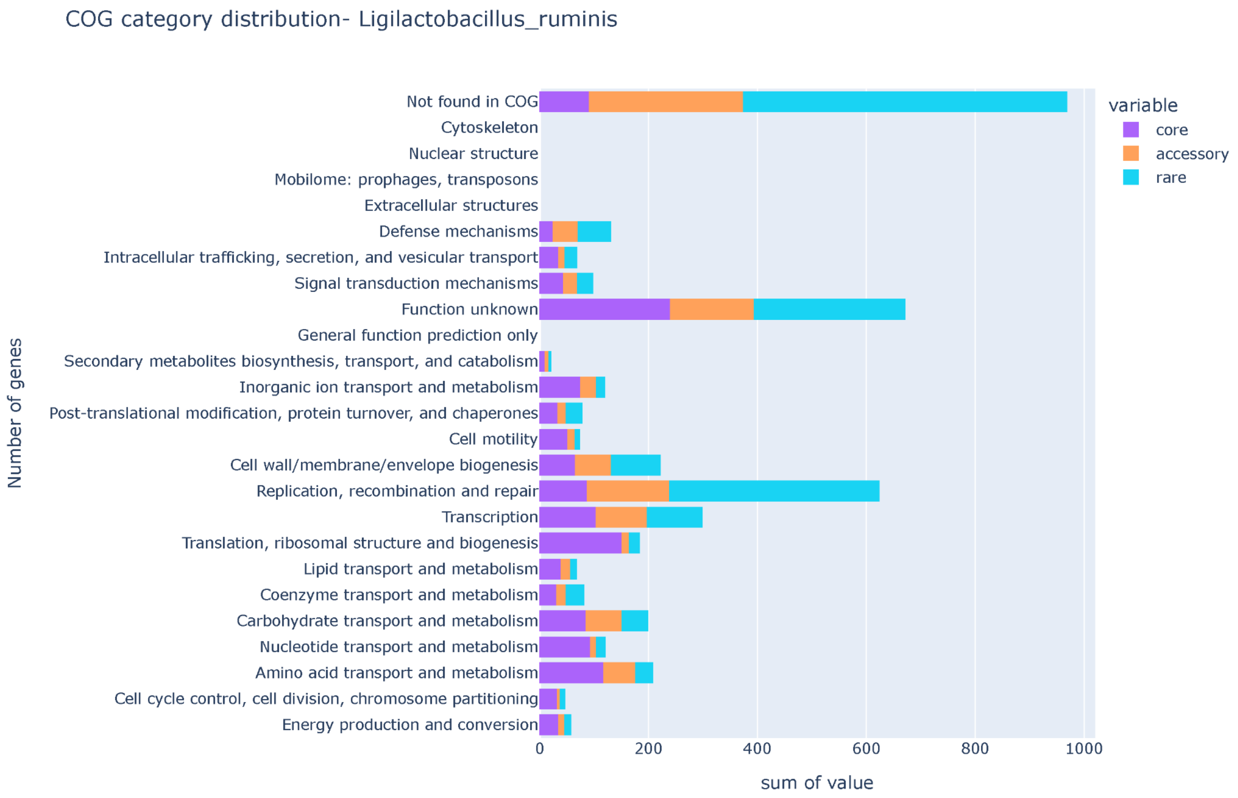


Loading [MathJax]/extensions/MathMenu.js

***Supplementary Figure S46: The barplot depicts the COG functions among core, accessory, and rare pangenome in Ligilactobacillus ruminis. The barplot is between the number of genes vs COG functions. Note: " - " refers to genes with no COG annotation.***

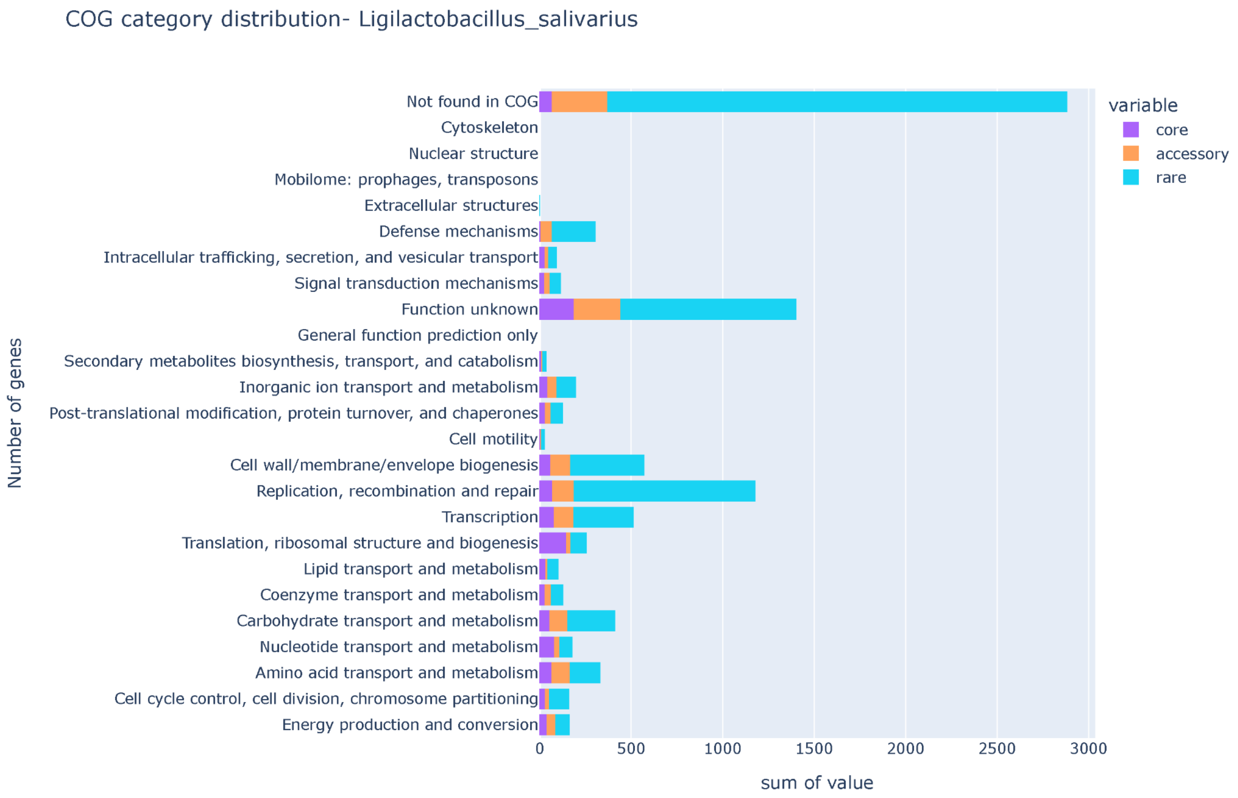


Loading [MathJax]/extensions/MathMenu.js

***Supplementary Figure S47: The barplot depicts the COG functions among core, accessory, and rare pangenome in Ligilactobacillus salivarius. The barplot is between the number of genes vs COG functions. Note: " - " refers to genes with no COG annotation.***

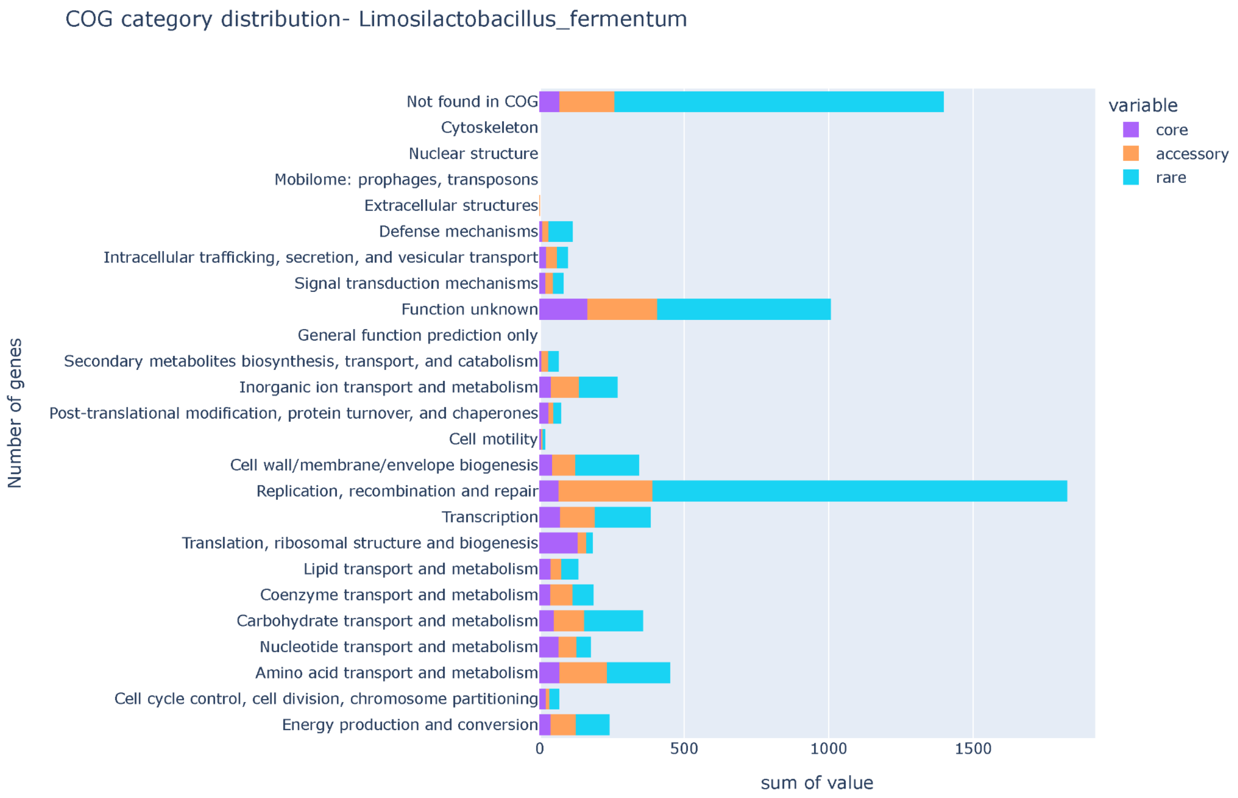


Loading [MathJax]/extensions/MathMenu.js

***Supplementary Figure S48: The barplot depicts the COG functions among core, accessory, and rare pangenome in Limosilactobacillus fermentum. The barplot is between the number of genes vs COG functions. Note: " - " refers to genes with no COG annotation.***

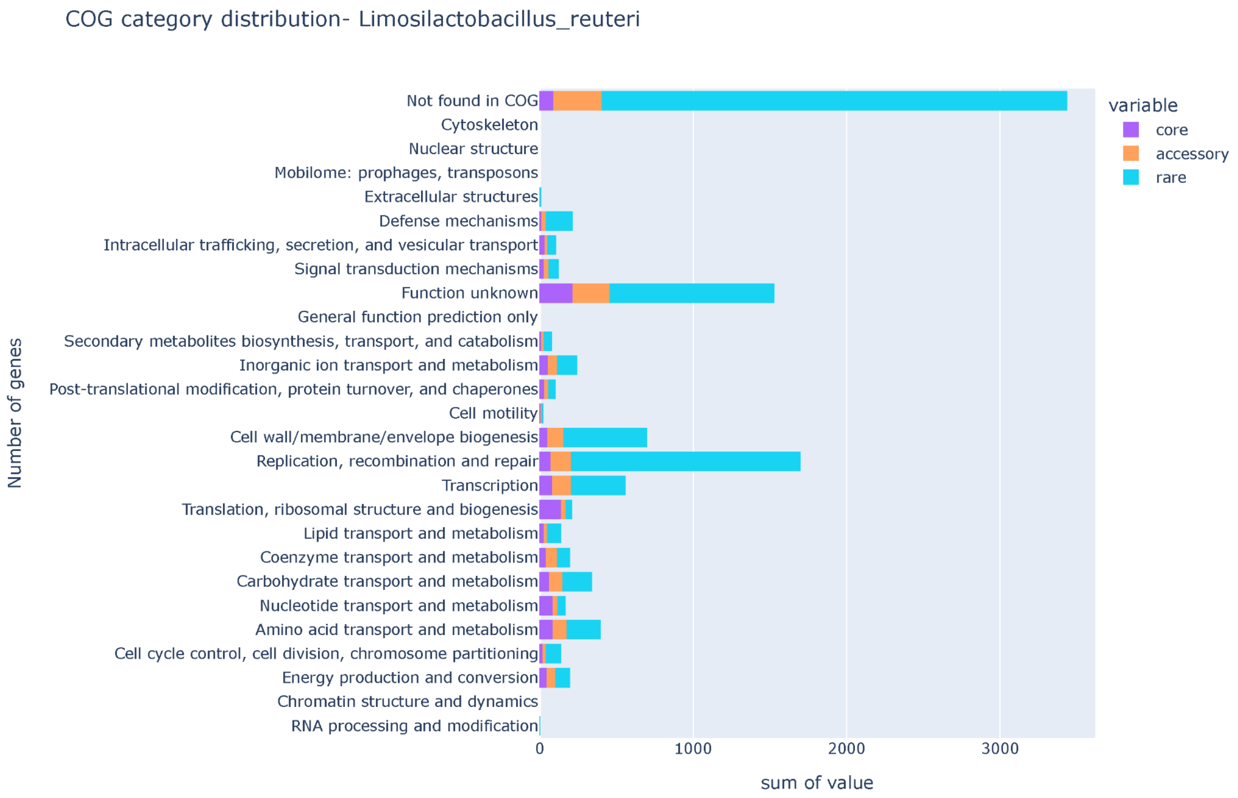


Loading [MathJax]/extensions/MathMenu.js

***Supplementary Figure S49: The barplot depicts the COG functions among core, accessory, and rare pangenome in Limosilactobacillus reuteri. The barplot is between the number of genes vs COG functions. Note: " - " refers to genes with no COG annotation.***

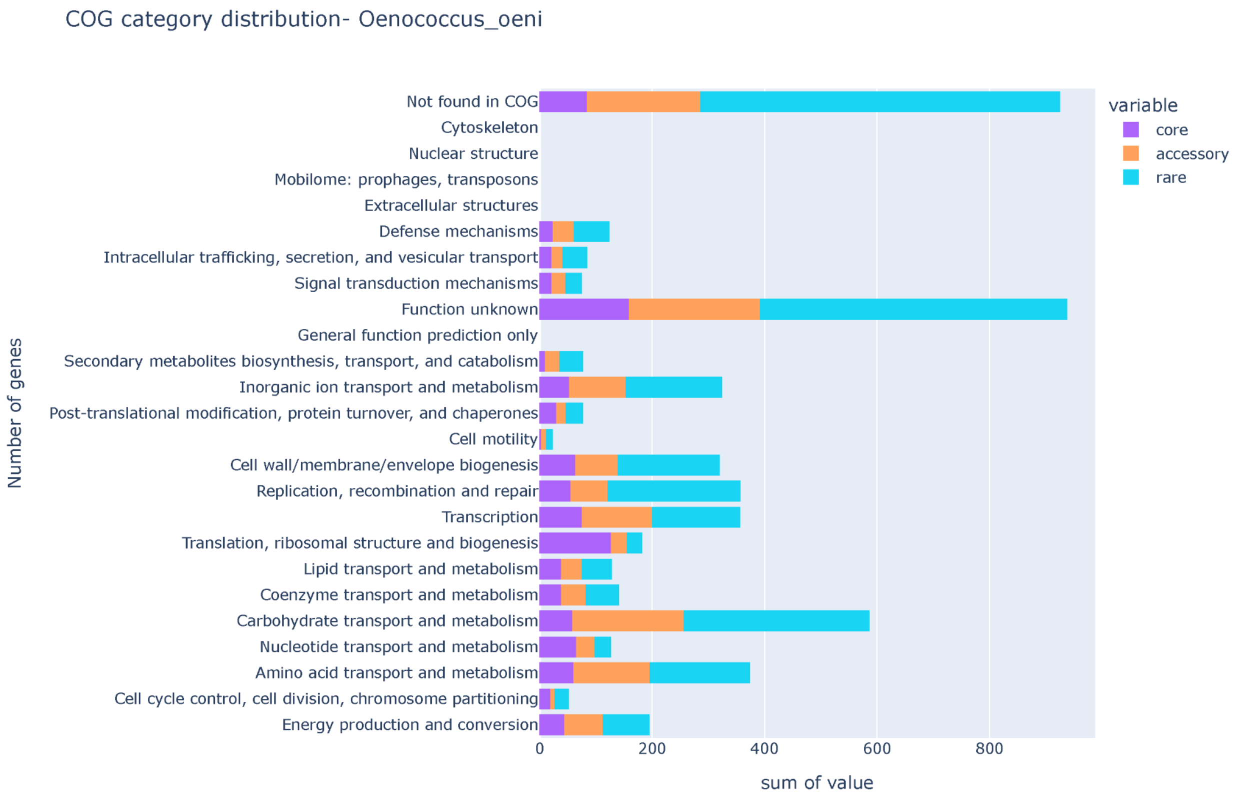


Loading [MathJax]/extensions/MathMenu.js

***Supplementary Figure S50: The barplot depicts the COG functions among core, accessory, and rare pangenome in Oenococcus oeni. The barplot is between the number of genes vs COG functions. Note: " - " refers to genes with no COG annotation.***

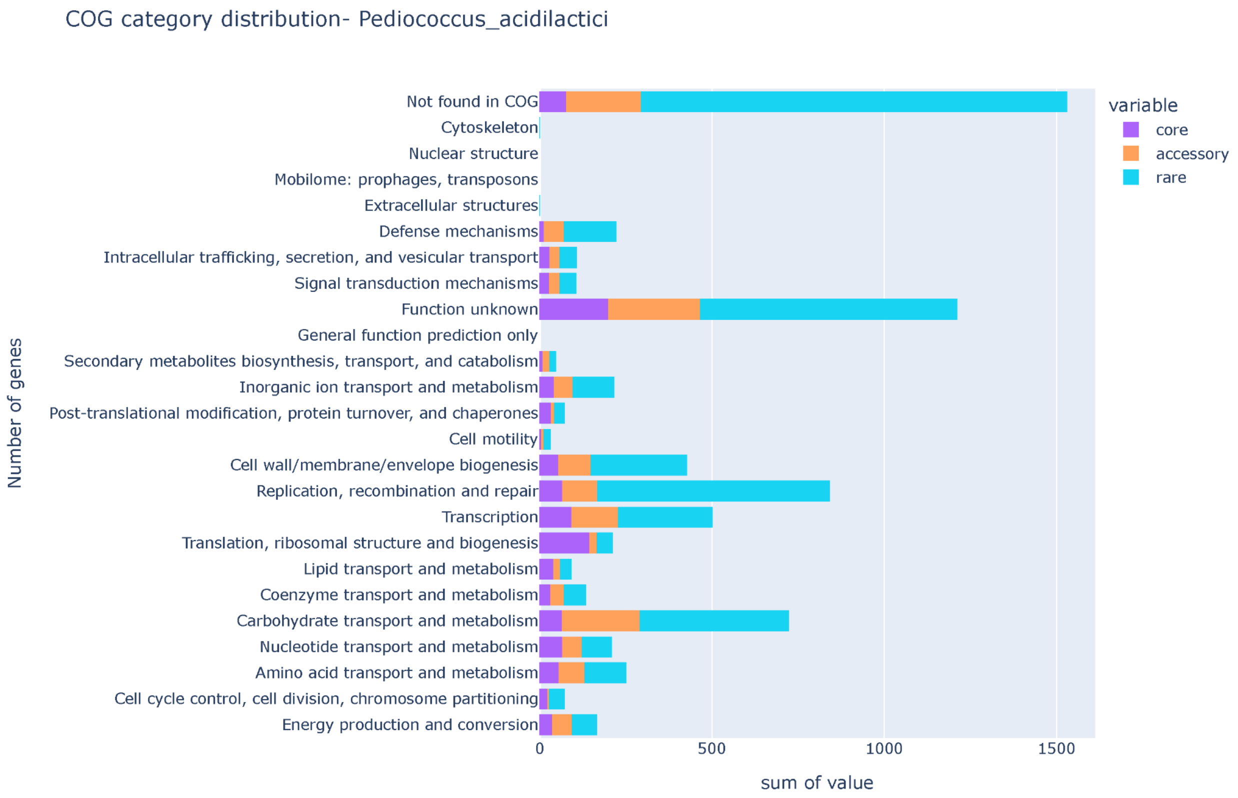


***Supplementary Figure S51: The barplot depicts the COG functions among core, accessory, and rare pangenome in Pediococcus acidilactici. The barplot is between the number of genes vs COG functions. Note: " - " refers to genes with no COG annotation.***

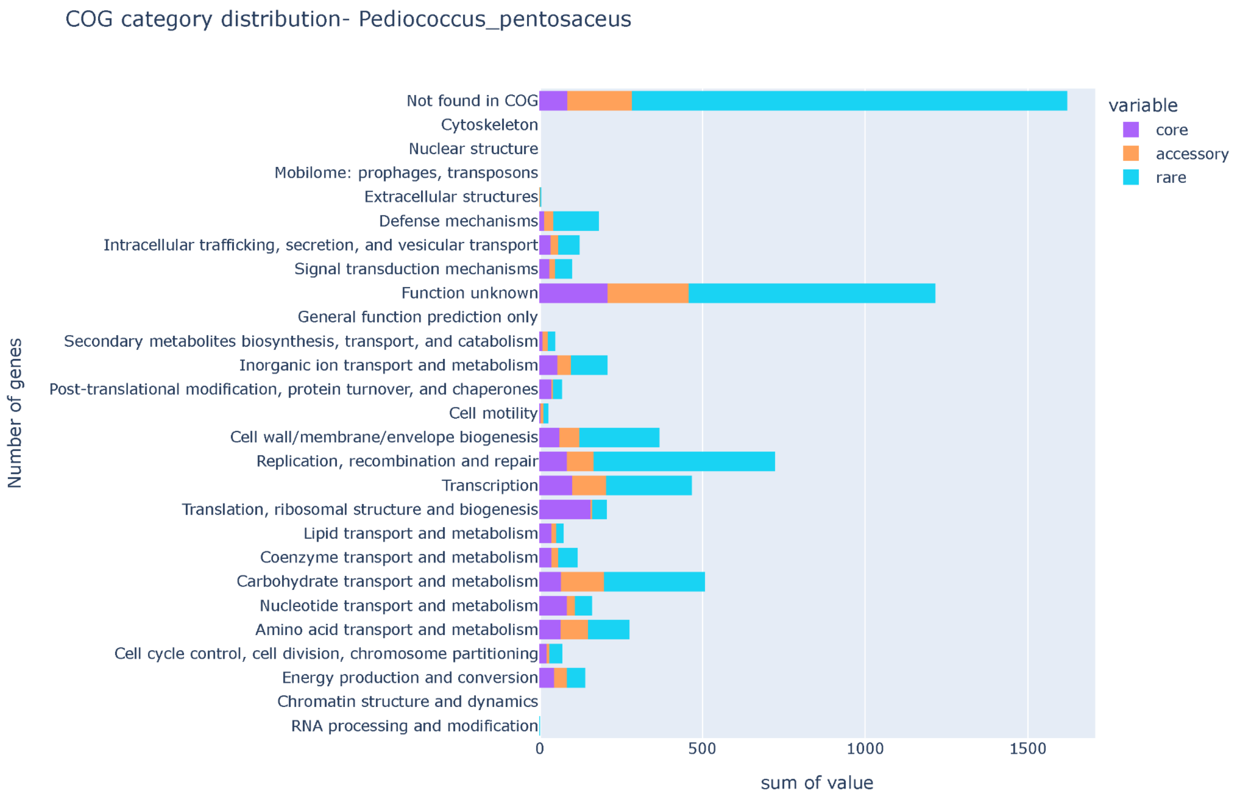


Loading [MathJax]/extensions/MathMenu.js

***Supplementary Figure S52: The barplot depicts the COG functions among core, accessory, and rare pangenome in Pediococcus pentosaceus. The barplot is between the number of genes vs COG functions. Note: " - " refers to genes with no COG annotation.***

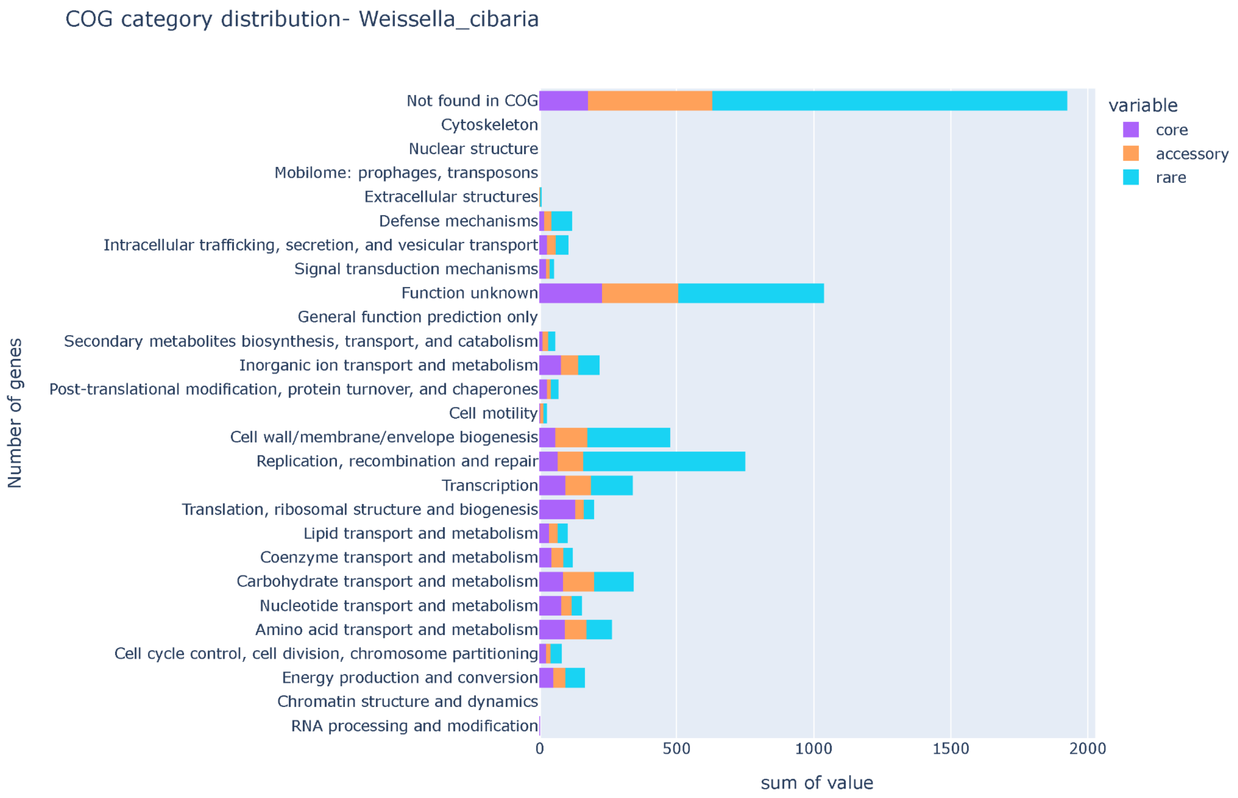


Loading [MathJax]/extensions/MathMenu.js

***Supplementary Figure S53: The barplot depicts the COG functions among core, accessory, and rare pangenome in Weissella cibaria. The barplot is between the number of genes vs COG functions. Note: " - " refers to genes with no COG annotation.***

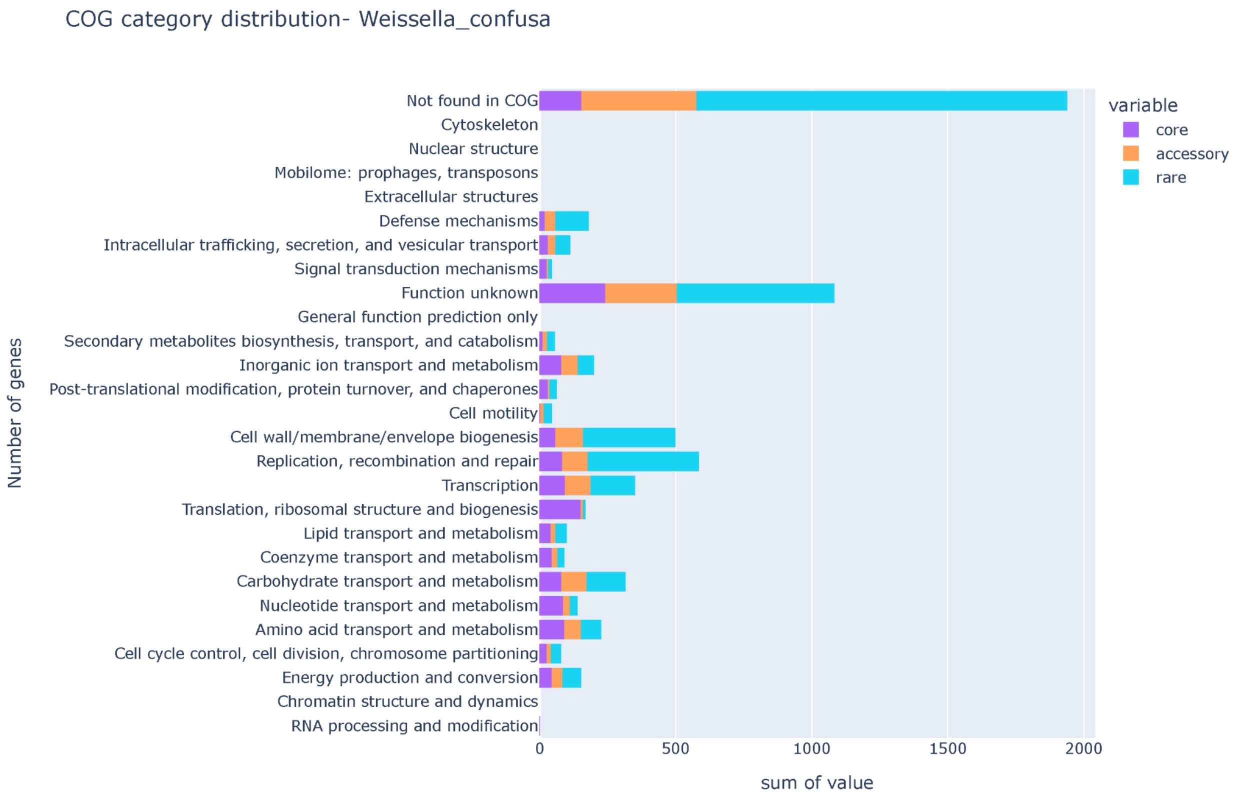


Loading [MathJax]/extensions/MathMenu.js

***Supplementary Figure S54: The barplot depicts the COG functions among core, accessory, and rare pangenome in Weissella confusa. The barplot is between the number of genes vs COG functions. Note: " - " refers to genes with no COG annotation.***

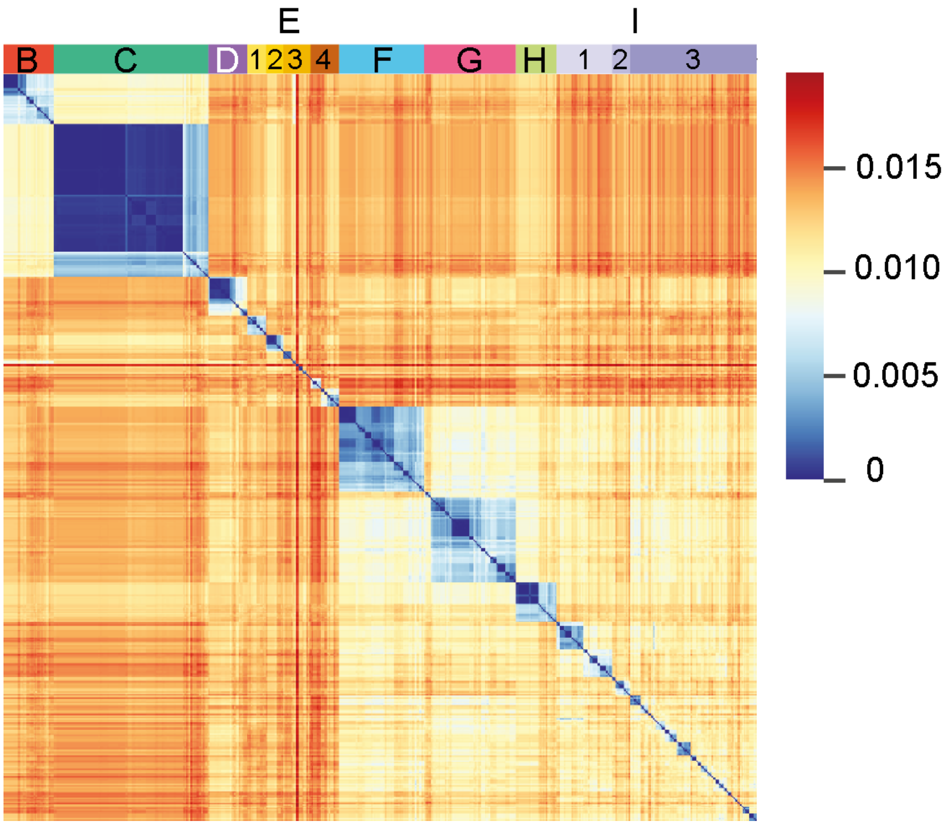


***Supplementary Figure S55: A clustermap of Mash distances for all L. plantarum strains with the phylogenetically distinct Mash Cluster A removed. All the clusters remain stable, with Cluster E splitting into 4 subclusters and Cluster I splitting into 3 subclusters.***

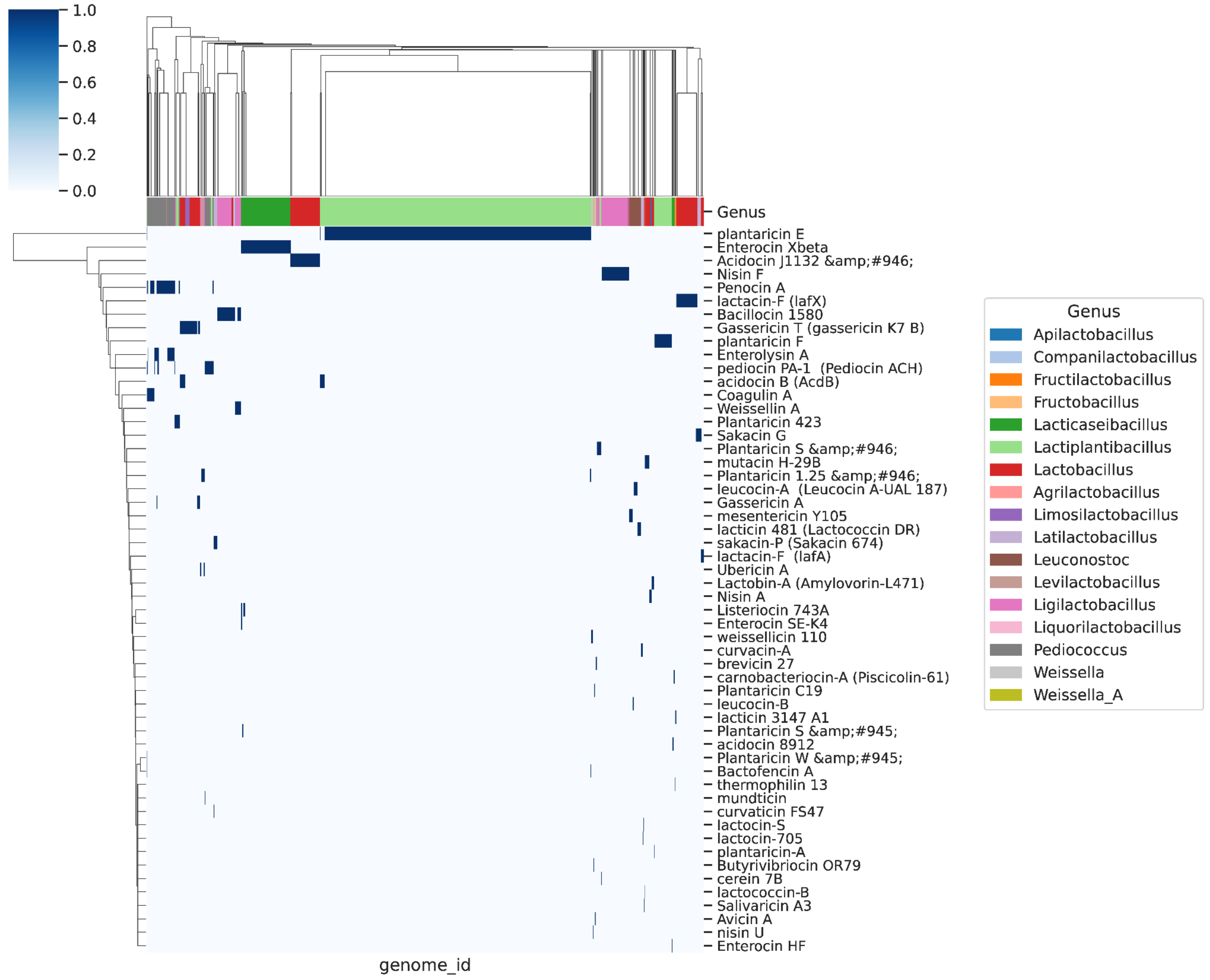


***Supplementary Figure S56: Biosynthetic gene clusters with BLAST hits (50%) in Bactibase.***
